# Activity-targeted metaproteomics uncovers rare syntrophic bacteria central to anaerobic community metabolism

**DOI:** 10.1101/2025.07.15.664484

**Authors:** Skyler Friedline, Elizabeth McDaniel, Matthew Scarborough, Maxwell Madill, Kate Waring, Vivian S. Lin, Rex R. Malmstrom, Danielle Goudeau, William Chrisler, Morten K. D. Dueholm, Leo J. Gorham, Chathuri J. Kombala, Lydia H. Griggs, Heather M. Olson, Sophie B. Lehmann, Nathalie Munoz, Jesse Trejo, Nikola Tolic, Ljiljana Pasa-Tolic, Sarai Williams, Mary Lipton, Steven J. Hallam, Ryan M. Ziels

## Abstract

Syntrophic microbial consortia can contribute significantly to the activity and function of anoxic ecosystems, yet are often too rare to study their *in situ* physiologies using traditional molecular methods. Here, we combined bioorthogonal non-canonical amino acid tagging (BONCAT), stable isotope probing, and metaproteomics to improve the recovery of proteins from active members and track isotope incorporation in an anaerobic digestion community. Both click chemistry-enabled cell-sorting and direct protein pulldown coupled to metaproteomics improved recovery of isotopically labeled proteins during anaerobic acetate oxidation. Resulting labeled protein expression profiles revealed elevated activity of a rare and so-far uncharacterized syntrophic bacterium belonging to the family *Natronincolaceae*. BONCAT-based capture of newly translated proteins provided direct molecular evidence for the expression of a previously hypothesized oxidative glycine pathway for syntrophic acetate oxidation by this microorganism, showcasing the potential of targeted metaproteomics to characterize rare and active cells central to community metabolism in natural and engineered ecosystems.

---

Microorganisms in natural and engineered ecosystems drive matter and energy conversion processes through distributed metabolic networks at the individual, population and community levels of biological organization^1,2^. Understanding the contributions of individual cells and interacting populations to community level metabolic fluxes is a key challenge in microbial ecology that would further our capacity to engineer and manage microbiomes and fully harness their vast metabolic potentials in the context of biotechnology innovation^3^. However, the irreducible complexity and resistance to axenic isolation inherent to natural microbial communities often necessitates *in situ* approaches to decipher their underlying ecological interactions^4^. Obfuscating this task is the observation that members with relatively low biomass abundance can contribute significantly to particular forms of metabolic flux within the community, and that these rare taxa are often overlooked when using traditional multi-omics (DNA, RNA, and protein) methods^5,6^.

One key example of rare but metabolically active community members are syntrophic consortia in anaerobic microbiomes^7,8^. Due to the minimal free energy yields associated with syntrophic energy metabolism^9^, such organisms typically remain low in abundance but can contribute significantly to substrate turnover^10^. For instance, syntrophic acetate oxidizing bacteria (SAOB) are notoriously difficult to isolate in pure culture^11^, and their metabolism yields such little energy (e.g., −10 kJ/mol^11^) that their doubling time is exceedingly slow (e.g., ranging from 3-20 days^11^). As a result, the metabolic pathways used by SAOB remain difficult to differentiate from other functional groups such as acetogens that inhabit the same environments. Yet, in anoxic environments, over 60% of methane is typically generated via acetate^12,13^, highlighting the importance of further understanding such cryptic syntrophic metabolisms. This need is particularly acute in engineered ecosystems producing bioenergy, as well as natural environments closely intertwined with global carbon cycling.

Protein is of particular interest when assessing the active fraction of microbial communities, as it presents contemporaneous information related to community metabolism^14^. Stable isotope probing (SIP)-informed metaproteomics has been employed to track substrate incorporation by syntrophs within complex anaerobic microbiomes by Mosbæk et al.^15^ However, in that study, only 5 ^13^C-labeled peptides were identified after 48 hours of SIP incubation, which is likely attributed to the low biomass abundance of the syntrophs relative to the background community^15^. Because metaproteomic analysis is compositional, the number of identified labeled peptides decreases as the relative abundance of the active biomass decreases^16^, due to unlabeled protein from non-active members overwhelming the proteome. One possible way to circumvent this limitation would be to ‘enrich’ for proteins produced by the most active community members. The substrate analog probing technique of bioorthogonal noncanonical amino acid tagging (BONCAT) is one such enrichment approach that uses amino acid analogues compatible with click chemistry to enable sorting and/or purification of recently translated proteins or the cells containing them^17–20^. As rare and active members like syntrophic microorganisms still synthesize new proteins^21,22^, we hypothesized that the enrichment of their proteins using BONCAT would enable more sensitive detection of labeled peptides from rare members in protein-SIP compared to ‘conventional metaproteomics’ (e.g., without protein enrichment).

In this work, we applied two different BONCAT enrichment techniques to enrich the SIP signal of SAOB in metaproteomics data from an anaerobic microbiome, in order to provide ecophysiological insights into this rare but important metabolism. The first BONCAT enrichment approach was based on sorting active cells using fluorescent activated cell sorting (FACS) prior to protein extraction for mass spectrometry. The second BONCAT technique involved protein extraction from bulk biomass followed by purification of newly-translated proteins using click chemistry and a biotin-streptavidin bead pulldown assay prior to preparation for mass spectrometry. We found that both of these techniques improved the detection of a highly-active and low-abundance syntrophic microorganism that significantly contributed to acetate turnover in microcosms established with sludge from a full-scale anaerobic digestion (AD) bioenergy facility. The BONCAT-SIP approaches described here therefore represent a powerful toolbox for unraveling the contributions of active microorganisms to microbial communities, a facet of microbial ecology that holds great potential for improving our ability to link field processes to microbial agents within natural and engineered ecosystems^23^.

## Results

### Long-read and short-read metagenome sequencing yielded a comprehensive genome database of the anaerobic digester community

Biomass from a full-scale AD bioenergy facility (Supplemental Figure S1) was routinely sampled over the course of 12 months, and extracted DNA was sequenced using a combination of short-read (Illumina; *n* = 8 samples) and long-read (PacBio; *n* = 3 samples) technologies (Figure 1). Using both long- and short-read metagenomic sequencing, assembly, and binning workflows, a dereplicated genome bin set was recovered containing 466 high-quality^24^ (including 22 circular) and 446 medium-quality draft metagenome assembled genomes (MAGs) (Figure 2; Supplemental Tables S1 and S2). This metagenomic sequencing effort added 334 taxonomically distinct high-quality MAGs to the existing collection of 799 high-quality de-replicated MAGs from ADs retrieved across 134 studies by Campanaro et. al.^25^, representing a 42% increase (Figure 2). Our set of 912 MAGs covered 8 phyla that were not represented in the previous global AD MAG database^25^ (Figure 2), possibly owing to the relatively high sequencing depth and multiple sequencing approaches used in this study. Correspondingly, the only circular genomes (*n=*22) for representative AD MAGs originated from this study (Figure 2), and included members of the 8 added phyla, such as the DPANN archaea, *Iainarchaeota*.

**Figure 1:**
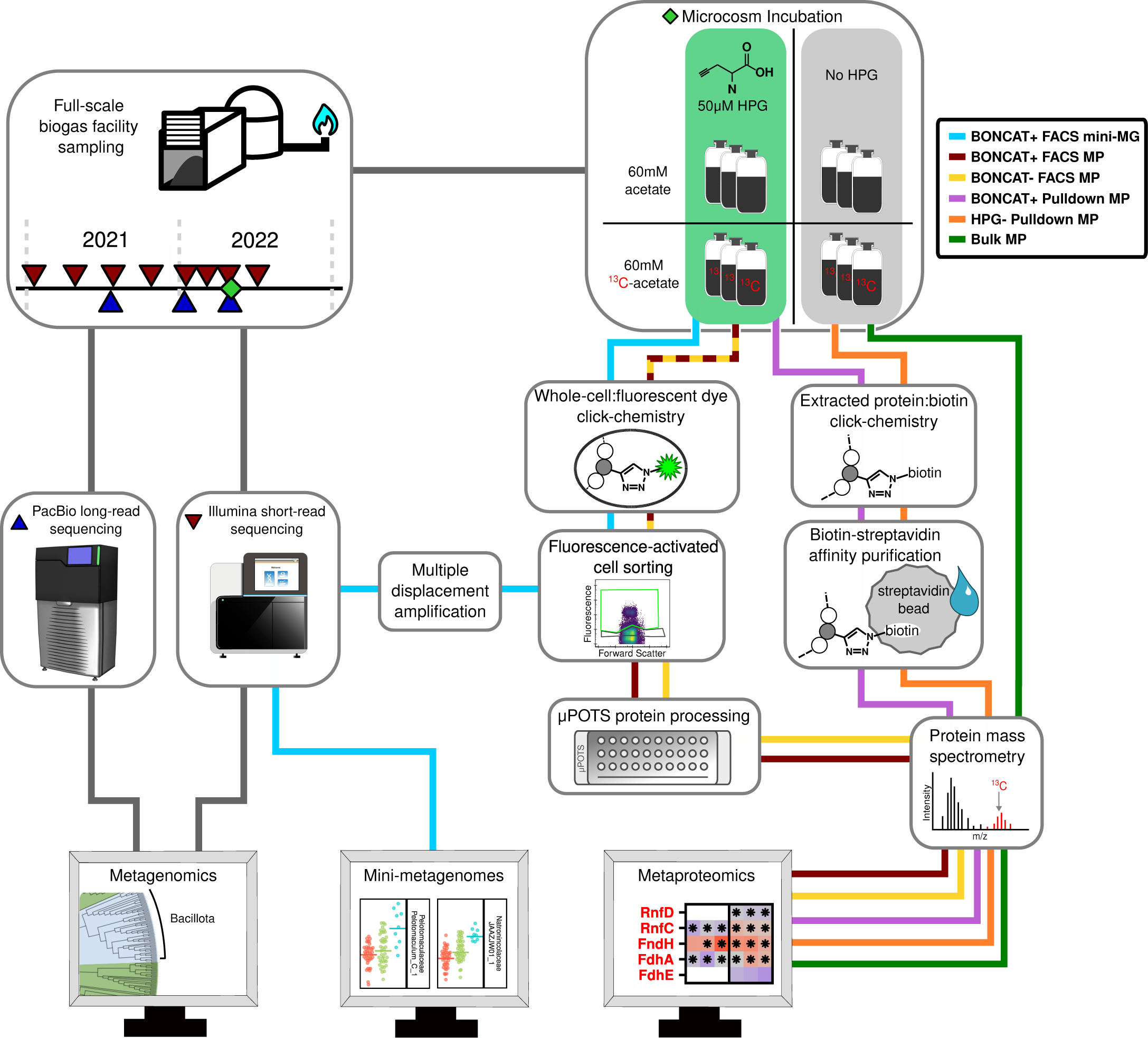
Conceptual overview of experimental design. Liquid digestate samples collected over two years from the Surrey Biofuel Facility were used to prepare a time series metagenome involving multiple sequencing methods. Liquid digestate was also collected to inoculate microcosm incubations with a combination of metabolic labeling substrates. Biomass samples from the microcosms were used to generate mini-metagenomes and metaproteomes.

**Figure 2:**
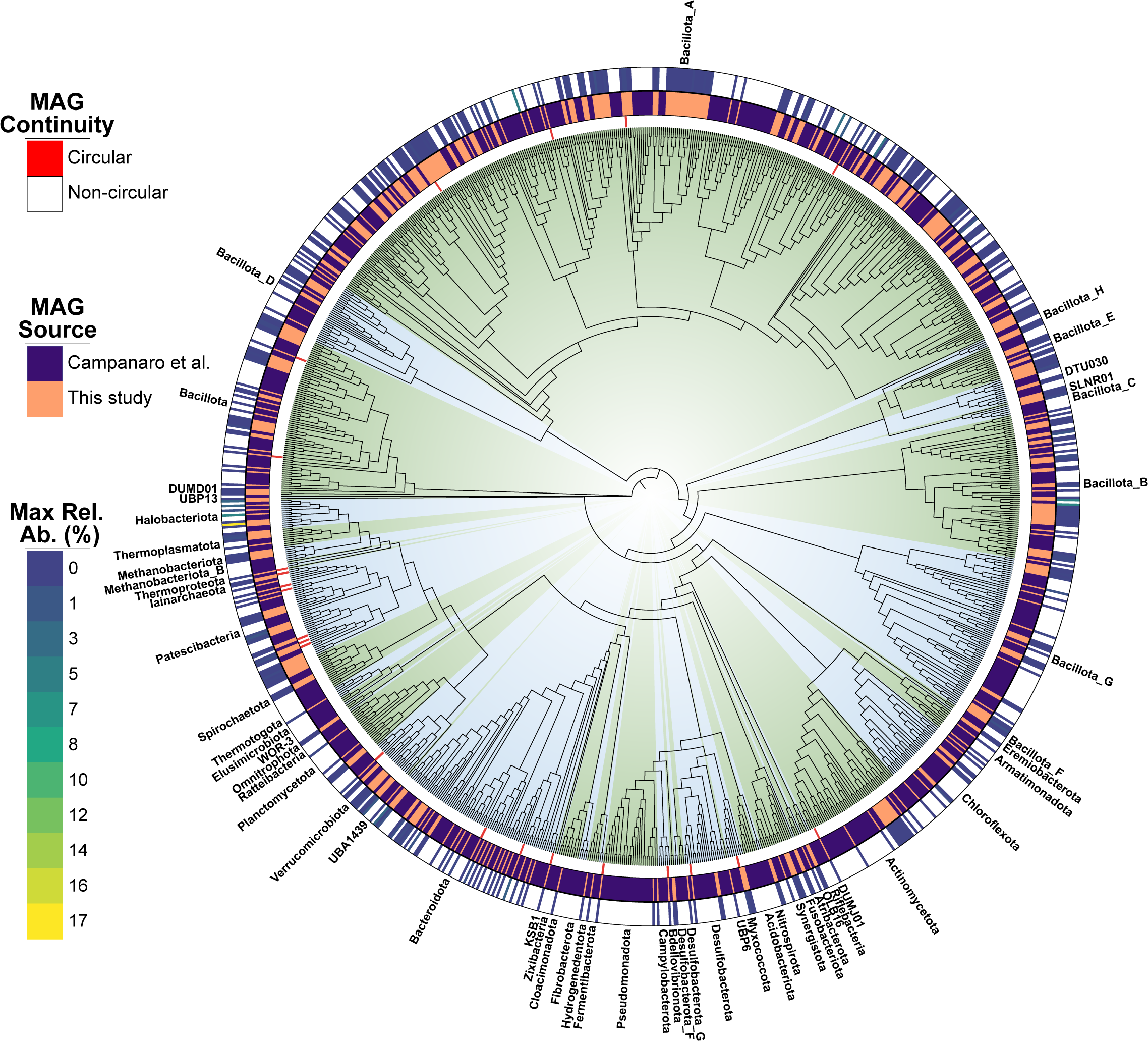
Comprehensive genomic database of the study anaerobic digester. Phylogenomic tree of concatenated single-copy marker genes from high-quality MAGs retrieved from this study along with the collection reported by Campanaro et al.^25^ The inner nodes of the tree are colored by phyla (alternating green and blue), as labeled on the outer ring. The middle rings, moving from inside to outside, correspond to: (1) MAG continuity, (2) MAG source, and (3) the maximum relative abundance (%) observed in the time-series metagenomic survey. The phylogenomic tree was visualized and annotated with iTol.^76^

The mapping rates of short-reads from time-series samples to the de-replicated set of MAGs ranged from 64.0 to 75.4% (Supplemental Table S3). Among the most abundant MAGs (by read mapping depth) in every sample were population genome bins affiliated with *Methanoculleaceae_Methanoculleus_5* (ranging from 5.10 - 17.8% relative abundance, n = 8) and *Cloacimonadaceae_UBA4175_1* (2.97 - 12.3%, n = 8) (Figure 2; Supplemental Figure S2). Closely related members (>95% average nucleotide identity (ANI)) of these two genomes have been previously observed in other AD systems (NCBI Accessions: GCA_012797575.1^25^ and GCA_030058025.1^26^, respectively). Only 39 of the 912 MAGs had greater than 1% relative abundance in any sample (Figure 2; Supplemental Table S3), suggesting that many low-abundance members may be critical for overall community function in this anaerobic ecosystem.

### BONCAT-based activity and methane production were concurrent in anaerobic microcosms

Anaerobic SIP microcosms were carried out to discern the fate of acetate in the anaerobic digestion microbiome and identify active members. SIP incubations were conducted using either universally ^13^C-labeled acetate (i.e., ^13^C-1,2-acetate), methyl-position ^13^C-labeled acetate (i.e., ^13^C-2-acetate) or unlabelled acetate (i.e., ^12^C-1,2-acetate). In addition, homopropargylglycine (HPG) was amended to a set of microcosms to enable BONCAT-based activity profiling in combination with SIP (Supplemental Table S4).

Metabolic activity was approximated by the fraction of cells from the HPG^+^ incubation that had elevated BONCAT fluorescence following a click chemistry reaction using a fluorescent dye (i.e. BONCAT^+^). We measured BONCAT^+^ cell fractions using FACS combined with either metaproteomics or metagenomics (Figure 3a and Supplemental Figures S3 and S4). The proportion of cells identified as BONCAT^+^ increased between the 24-hour (mean ± s.d. 6.5 ± 3.1%, *n =* 6 in the metaproteome sorting experiment and 9.2 ± 0.83%, *n =* 3 in the mini-metagenome sorting experiment) and the 72-hour (metaproteome 13.4 ± 1.5%, *n =* 6 and mini-metagenome 11.7 ± 1.5%, *n =* 3) sampling events before declining again at 96 hours (Figure 3b). In a similar trend, the methane production rate in the microcosms was most rapid within the first 24 hours (mean ± s.d. 26.2 ± 1.84 cm^3^/d, *n* = 48) and declined slowly (20.5 ± 4.5 cm^3^/d, *n* = 36 between 24 and 48 hours, 8.0 ± 2.1 cm^3^/d, *n =* 24 between 48 and 72 hours) before ultimately producing methane at the same rate as the no-substrate controls between 72 and 96 hours (Figure 3c).

**Figure 3:**
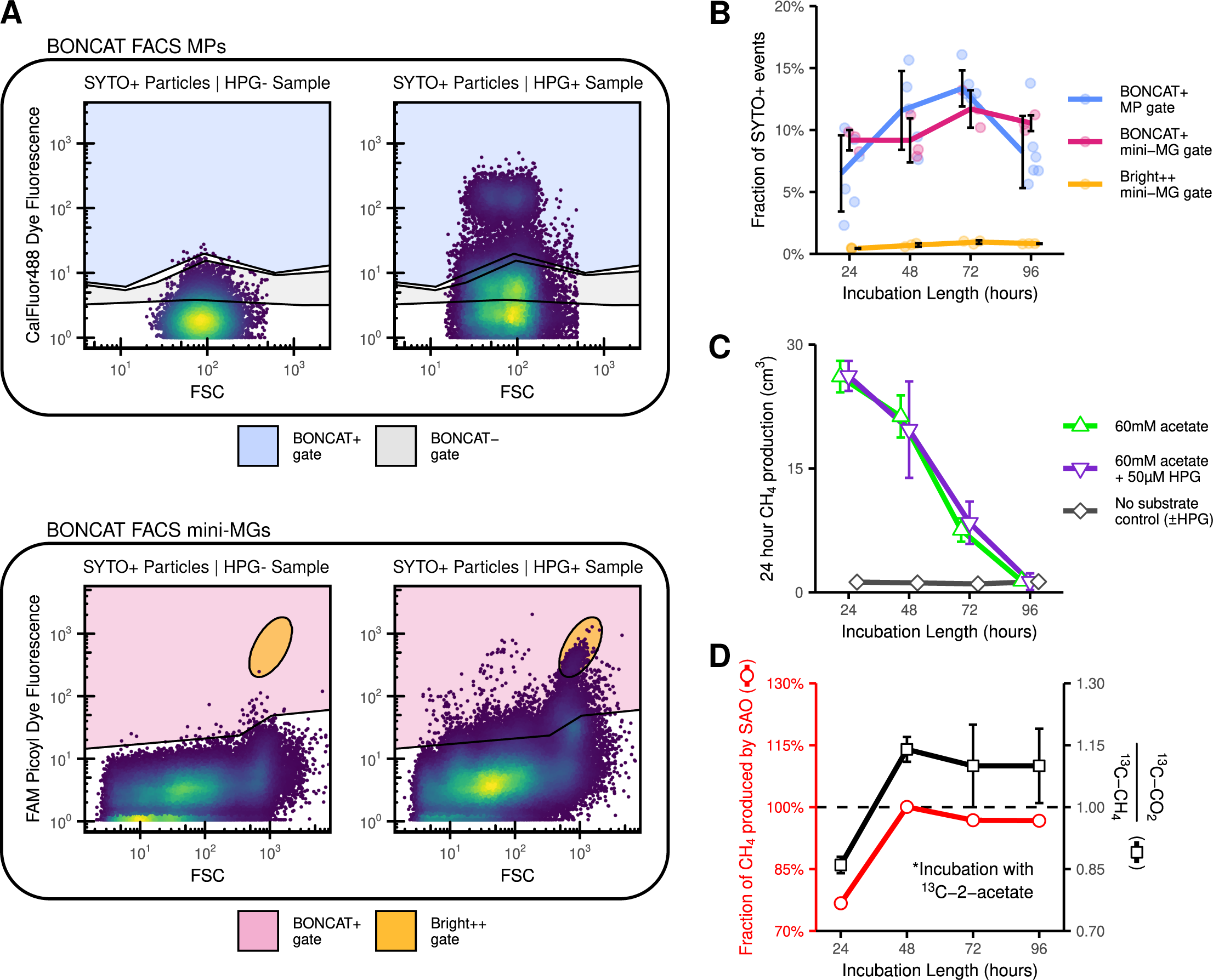
Enrichment of an active sub-population with BONCAT-targeted metaproteomics and mini-metagenomics. (A) The sorting strategy consisted of first identifying whole cells using DNA dye staining (SYTO) fluorescence, which were subsequently assessed for BONCAT fluorescence (CalFluor488, Ex: 488 nm, Em: 520/15 nm). Cells were sorted using a variety of gates in two separate experiments (see Online Methods for details). Only the BONCAT sorting gate is shown in these plots, which was applied to SYTO-positive cells. The left plots come from a no-HPG control condition while the right plots show an example HPG^+^ condition. (B) The fraction of flow cytometry events that were BONCAT+ varied over time and peaked at 72 hours (lines and error bars indicate the mean ± s.d, n = 6). (C) Methane production occurred primarily before 72 hours and was not affected by HPG (points and horizontal error bars indicate the mean ± s.d, n = 6 - 24). (D) Measured isotopic ratio of the headspace methane from an incubation with ^13^C-2-acetate (right axis, in black; points and vertical bars represent mean ± s.d, n = 3) was used to model the relative contributions of the syntrophic acetate oxidation and acetoclastic methanogenesis pathways to total methane production (left axis, in red; points represent model-optimized values with standard error too small to be visible).

Incorporation of noncanonical amino acids for BONCAT has been shown to cause slight changes in metabolism within the study of a single bacterial species^27^. Here, we observed no statistical difference between the methane volume produced in the samples incubated with HPG (i.e., HPG^+^) and those without (i.e., HPG^−^) at any sampling event (Student’s t test, *P* = 0.9, 0.3, 0.3, 0.9 and *n* = 24, 18, 12, 6 at 24, 48, 72, and 96 hours respectively), indicating that HPG did not impact the rate of substrate turnover during the incubations. Moreover, there were no apparent differences in metabolite profiles within HPG^+^ and HPG^−^ samples determined via GC-MS based metabolomics (Supplemental Figure S5), further indicating that HPG did not noticeably impact community metabolism. Based on the theoretical methane potential of the acetate amendment, approximately 91.6% (including both HPG^+^ and HPG^−^ conditions, s.d. 10.3%, n = 24) substrate conversion was achieved in the microcosms by 72 hours (Supplemental Figure S6).

Acetoclastic methanogenesis partitions the carboxyl group of acetate into CO_2_, while SAO partitions both carboxyl and methyl carbon atoms of acetate into CO_2_^28^. We therefore tracked the isotopic ratio of the evolved CO_2_ and CH_4_ in microcosms amended with methyl-^13^C-labeled acetate (e.g., ^13^C-2-acetate) in comparison to universally-^13^C-labeled acetate (Figure 3d and Supplemental Table S5). According to the isotope fractionation model of Mulat et al.^28^, SAO was responsible for the vast majority of acetate conversion in the microcosms, ranging from 76.7% (s.d. 1.7%, n = 10,000, Monte Carlo simulation) of total methane production at 24 hours and increasing to an average of 98% (s.d 3.8%, n = 10,000, Monte Carlo simulation) for the remainder of the incubations (Figure 3d and Supplemental Table S6). The SIP incubations therefore provided an opportunity to investigate members involved in SAO within the full-scale anaerobic digester community.

### BONCAT-targeted mini-metagenomics enriched two highly active bacteria

Three populations of cells were sorted to generate mini-metagenomes from HPG-incubated samples: (1) all cells passing the SYTO^+^ gate (i.e., SYTO^+^), (2) all SYTO^+^ cells passing the BONCAT^+^ gate (i.e., BONCAT^+^), and (3) BONCAT^+^ cells that clustered in a high-fluorescence, high forward scatter region (i.e. Bright^++^) (Figure 3a and Supplemental Figure S3). The Bright^++^ gate was targeted as a subset of the BONCAT^+^ population based on its unique fluorescence pattern and uniformity, indicating that it was a distinct flow cytometric population that was clearly BONCAT-labeled. The fraction of cells passing the Bright^++^ gate followed a similar trend to that of the BONCAT^+^ gate (described above), increasing from 0.45 ± 0.04% (mean ± s.d, *n* = 3) of SYTO^+^ events at 24 hours to 0.96 ± 0.14% at 72 hours before falling to 0.82 ± 0.02% at 96 hours (Figure 3a).

At each sampling time point, an average of 43 ± 1, 43 ± 2, and 9 ± 2 (mean ± s.d., *n* = 3) pools of 300 cells from the SYTO^+^, BONCAT^+^, or Bright^++^ gates were collected separately for metagenome sequencing. Two MAGs were statistically enriched (Wald test with 43 - 85 degrees of freedom, Bonferroni adjusted *P* < 0.001, log_2_ fold change > 2, relative abundance in at least one sample > 0.5%) at every time point in the Bright^++^ population compared to the BONCAT^+^ and SYTO^+^ populations (Figure 4a). Of these, *Pelotomaculaceae_Pelotomaculum_C_1* was additionally enriched at every time point in BONCAT^+^ samples compared to SYTO^+^ samples while *Natronincolaceae_JAAZJW01_1* was only statistically enriched at 24 and 48 hours (Figure 4a). Of these two MAGs, *Pelotomaculaceae_Pelotomaculum_C_1* was not detected in the previous global biogas genome database^25^ (at an ANI cutoff of 98%), while a MAG with 152 contigs was detected the database (GenBank accession: GCA_012519835.1) that was closely related (98.6% ANI) to *Natronincolaceae_JAAZJW01_1*, of which we obtained a circular genome (Supplemental Table S2).

**Figure 4:**
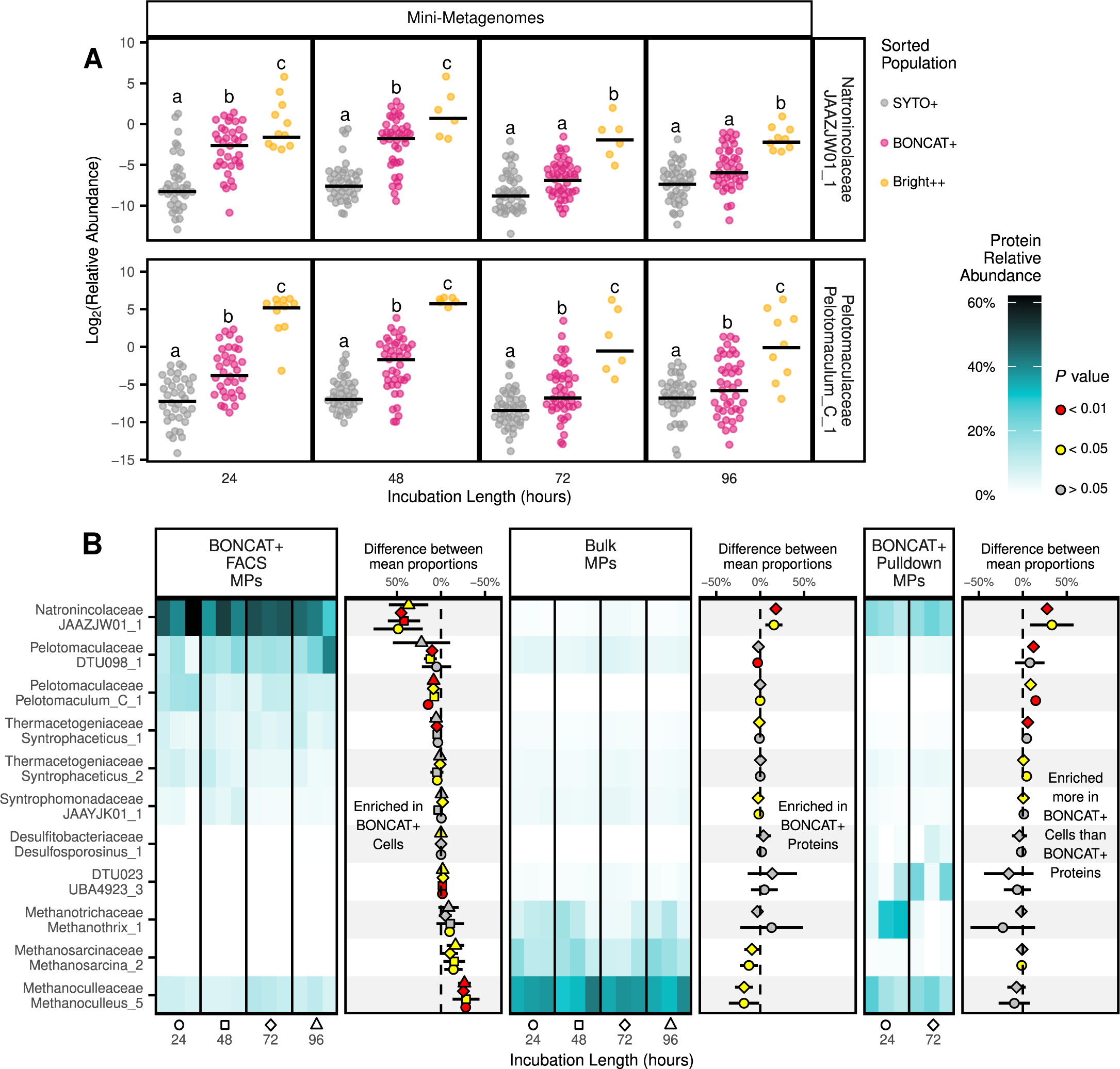
BONCAT-targeted methods enriched active populations in mini-metagenomes and metaproteomes. (A) Mapping reads from sorted cell mini-metagenomes to the time series metagenome revealed two MAGs were statistically enriched (Wald test P < 0.001, log_2_-fold change > 2, relative abundance > 0.5% in at least one sample) in the Bright++ population compared to both the BONCAT+ and SYTO+ populations (bar represents median, letters represent statistically similar groups, n = 6 - 45). (B) Relative protein abundance of the highest-contributing MAGs (>7% relative abundance in at least 1 sample) in the Bulk and BONCAT^+^ MPs. The points between heatmaps indicate the difference in means between the two flanking MPs and the rightmost subpanel references the BONCAT-FACS MP as if it was additionally shown on the far right, in order to compare the two BONCAT-enabled MPs. (point color indicates Student’s t-test P value and horizontal bars indicate 95% confidence interval, missing values were replaced with zeros, n = 3).

### BONCAT-enriched metaproteomes were dominated by an uncharacterized Natronincolaceae bacterium

To gain further insights into active microorganisms metabolizing acetate, the newly-translated proteome was profiled using BONCAT-informed metaproteomics. Metaproteomes (MPs) were generated using three separate methodologies: (1) untargeted metaproteomics of the bulk community (i.e., Bulk MPs); (2) activity-targeted metaproteomics of cells from HPG^+^ incubations sorted by FACS (i.e., BONCAT^+^ FACS MPs); (3) activity-targeted metaproteomics of proteins from HPG^+^ incubations enriched via biotin-streptavidin pulldown assays (i.e. BONCAT^+^ Pulldown MPs) (Figure 1). While little variation in composition was seen over time within the individual approaches, there were clear differences in the profiles obtained between methods (Figure 4b and Supplemental Figure S7). Mass spectra were searched using a protein database generated from the set of 912 dereplicated MAGs described above. Overall, between 89.2 - 99.6% of the total identified protein abundance in each sample was attributed to the 466 high-quality MAGs, indicating that high-quality MAGs were recovered for most of the detectable community biomass.

The relative biomass contributions (i.e., protein relative abundance) of the MAGs in each metaproteome were determined using a total protein approach^29–31^. In the Bulk MP samples, *Archaea* contributed the majority of detected protein abundance (minimum 56.9 ± 6.9% at 72 hours, *n* = 3; maximum 76.7 ± 0.1% at 0 hours, *n* = 2; mean ± s.d.). Across every Bulk MP sample, archaeal proteins were primarily attributed to three MAGs: *Methanoculleaceae_Methanoculleus_5* (minimum 30.1 ± 3.2% at 0 hours, *n* = 2; and maximum 37.0 ± 1.8% at 72 hours)*, Methanosarcinaceae_Methanosarcina_2* (minimum 10.2 ± 3.6% at 72 hours, *n* = 3; and maximum 19.9 ± 2.2% at 0 hours, *n* = 2) and *Methanotrichaceae_Methanothrix_1* (minimum 4.8 ± 2.9% at 72 hours, *n* = 3; and maximum 22.0 ± 1.1% at 0 hours, n = 2). Meanwhile, the bacterial protein fraction of the Bulk MPs was primarily assigned to the phylum *Bacillota* (minimum 18.8 ± 0.2% at 0 hours, *n* = 2; and maximum 33.0 ± 5.2% at 72 hours, *n* = 3). The most abundant *Bacillota* MAG was *Pelotomaculaceae_DTU098_1* (minimum 3.1 ± 0.1% at 0 hours, *n* = 2; and maximum 5.4 ± 1.0% at 72 hours, n = 3) followed by 8 other MAGs that had a mean relative abundance > 1% in at least one time point (Supplemental Figure S7).

Compared to the Bulk MP samples, the BONCAT^+^ FACS MP samples were marked by a large increase in the fraction of protein abundance from *Bacteria*, and specifically from the phylum *Bacillota* (3.4 ± 0.0, 3.4 ± 0.1, 2.7 ± 0.2, 3.5 ± 0.1 fold increases in relative abundance at 24, 48, 72, and 96 hours respectively; mean ± sd, *n* = 3 biological replicates) (Figure 4b). Of all MAGs, *Natronincolaceae_JAAZJW01_1* (phylum = *Bacillota_A*, class = *Clostridia*, Order = *Peptostreptococcales,* Family = *Natronincolaceae*) manifested the largest enrichment, increasing from 0.1 - 2.8% (*n* = 14 samples) relative abundance in the Bulk MPs to 27.9 - 62.0% (*n* = 12 samples) in the BONCAT^+^ FACS MPs (Student’s t test *P* = 0.02, 0.01, 0.00006, 0.02 at 24, 48, 72, and 96 hours respectively, *n* = 3 biological replicates). This MAG was completely absent from the corresponding BONCAT^−^ FACS MPs (Figure 3d and Supplemental Figure S7).

The only archaeal MAG with notable abundance in the BONCAT^+^ FACS MPs was *Methanoculleaceae_Methanoculleus_5*, which ranged from 6.4 - 12.9% (*n* = 12 samples) relative abundance, a significant decline from the Bulk MPs (Student’s t test *P* = 0.00006, 0.01, 0.00005, 0.001 at 24, 48, 72, and 96 hours respectively, *n* = 3 biological replicates) (Figure 4b). Conversely, *Methanotrichaceae_Methanothrix_1* and *Methanosarcinaceae_Methanosarcina_2* were completely absent from both BONCAT^+^ and BONCAT^−^ FACS MP populations, indicating they were excluded from FACS-based analysis. It is conceivable that the filamentous morphology of genus *Methanothrix* (> 100μm long^32^) and the aggregate morphology of genus *Methanosarcina*^33^ prevented their inclusion in FACS analysis due to pre-filtration, and/or to the FACS gating strategies employed here. Examination of the anaerobic digester biomass using fluorescence in situ hybridization (FISH) and microscopy revealed the presence of *Methanosarcina*-like tetrad clusters of archaeal cells, along with *Methanothrix*-like filamentous archaeal cells (Supplemental Figure S8), supporting the notion that these cell aggregates could have been excluded from FACS analysis. Aggregated cell morphologies could have also reduced the extent of BONCAT labeling of these methanogens if HPG concentrations were diffusion-limited into the biofilm, as we correspondingly observed lower BONCAT^+^ cell fractions with lower concentrations of amended HPG (Supplemental Figure S9).

The BONCAT^+^ Pulldown MPs showed more variation in total signal intensity and taxonomic composition between timepoints and replicates than observed in the Bulk MPs and BONCAT^+^ FACS MPs (Figure 4b). The fraction of total protein assigned to *Archaea* at 24 hours was 44.8 ± 6.7% (mean ± s.d, *n* = 3) in the BONCAT^+^ Pulldown MPs, which decreased to 24.0 ± 5.0% (*n* = 3) at 72 hours. This decrease in archaeal protein relative abundance was concurrent with the disappearance of *Methanotrichaceae_Methanothrix_1* (22.7 ± 14.8% at 24 hours, *n* = 3; and 1.7 ± 2.0% at 72 hours, *n* = 3; mean ± s.d.) (Figure 4b). In agreement with the BONCAT^+^ FACS MPs, the MAG experiencing the highest increase in relative abundance between the Bulk MPs and BONCAT^+^ Pulldown MPs was *Natronincolaceae_JAAZJW01_1*, which comprised 16.6 ± 3.9% (*n* = 3) and 20.2 ± 2.8% (*n* = 3) of the total protein abundance at 24 and 72 hours respectively (mean ± s.d.) (Figure 4b). This MAG was also entirely absent from the HPG^−^ Pulldown MPs (Supplemental Figure S7).

### BONCAT enrichment consistently selects for SIP-tagged proteins

We quantified ^13^C isotope abundance in peptides to examine whether the BONCAT-informed protein pools were more concentrated in the heavy isotope signal. Over the course of the SIP experiment, the peptide labeling ratio (LR) in Bulk MPs showed a modest increase, reaching a peak median value of 15.9% (Interquartile Range [IQR] = 22.6%; 261 ^13^C-containing peptides) at 96 hours (*n*=3) (Figure 5a). The quantity and extent of ^13^C labeling was higher at every time point in the BONCAT^+^ FACS MPs, which started with a median peptide LR of 18.2% (IQR = 18.7%; 296 labeled peptides) at 24 hours and increased at every time point to reach 58.0% (IQR = 23.3%; 905 labeled peptides) by 96 hours. The BONCAT^+^ Pulldown MPs maintained a nearly constant and elevated peptide LR at both time points (median = 69.3%, IQR = 30.0%, 545 labeled peptides at 24 hours; and median = 67.6%, IQR = 34.1%; 267 labeled peptides at 72 hours) (Figure 5a and Supplemental Table S7).

**Figure 5:**
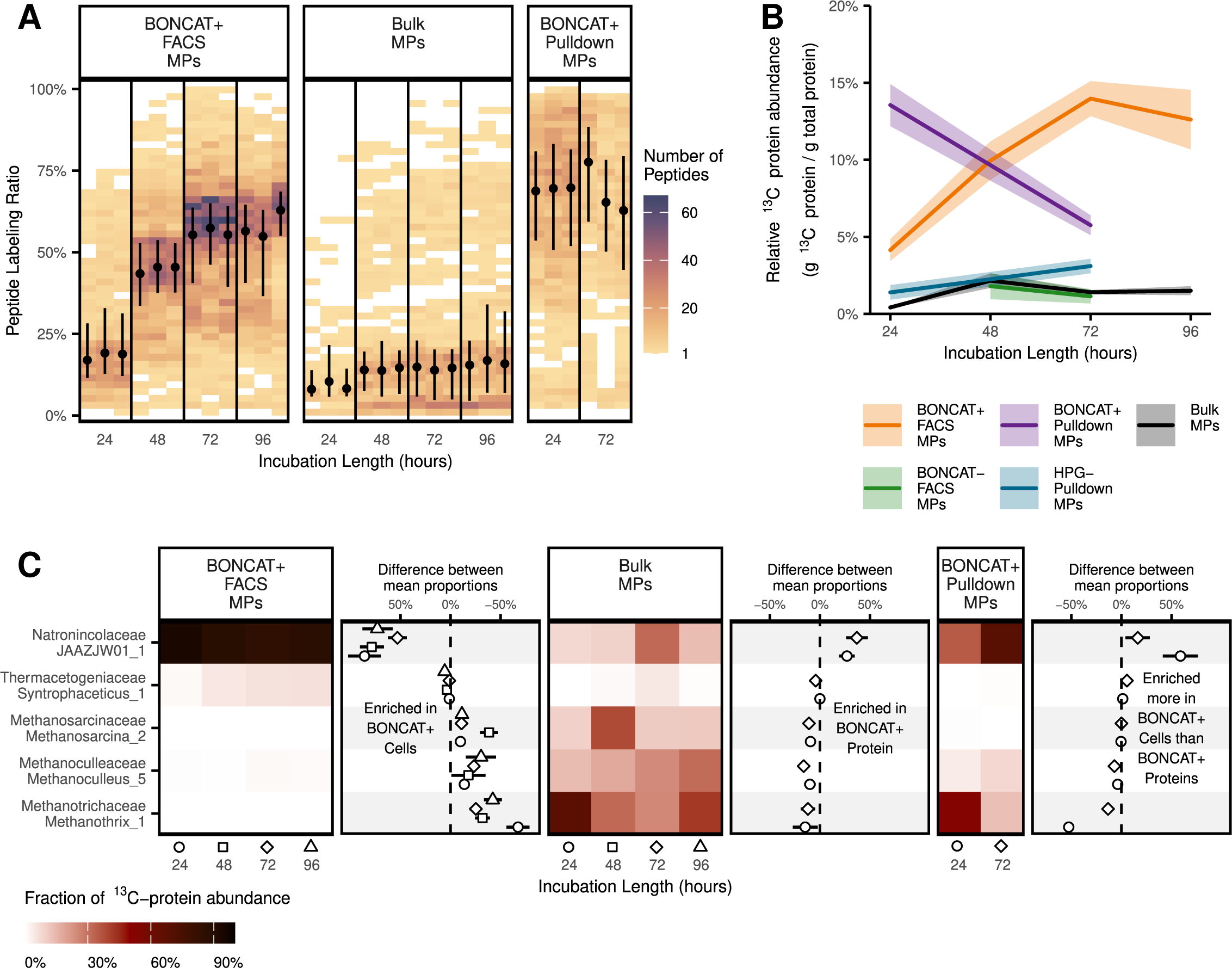
Tracking ^13^C incorporation in the metaproteome. (A) Tiled heatmap showing peptide counts (fill color) with different peptide labeling ratios (LRs) over time with the various metaproteome approaches (black circles represent median and error bars indicate interquartile range). (B) ^13^C relative protein abundance over time with the various metaproteome approaches (line indicates the mean and shaded area represents standard deviation, n = 3). (C) Tiled heatmap colored by the mean fraction of the ^13^C-labeled protein abundance among the most labeled MAGs (i.e. summed ^13^C-protein fraction in bins with > 5% in at least one sample). The points between heatmaps represent the difference in observed means between the flanking plots, and the rightmost subpanel references the BONCAT-FACS MP as if it was additionally shown on the far right, in order to compare the two BONCAT-enabled MPs. (Point shapes indicate time point and horizontal bars indicate bootstrapped 95% confidence interval from Monte Carlo simulation, missing values were replaced with zeros, n = 10,000).

The ^13^C-protein relative abundance (i.e. g ^13^C-protein/g total protein) in the Bulk MPs reached a maximum of 1.5 ± 0.3% at 96 hours (Figure 5b). The labeled proteins predominantly came from three methanogens (*Methanoculleaceae_Methanoculleus_5*, *Methanosarcinaceae_Methanosarcina_2,* and *Methanotrichaceae_Methanothrix_1)* and *Natronincolaceae_JAAZJW01_1* (Figure 5c and Supplemental Figure S10). While *Natronincolaceae_JAAZJW01_1* contributed very little to the total protein relative abundance in the Bulk MPs (minimum 0.2 ± 0.0% at 0 hours; and maximum 2.4 ± 0.5% at 72 hours; mean ± s.d.), this MAG contributed disproportionately to the ^13^C-protein relative abundance in the same samples (minimum 8.2 ± 3.1% of ^13^C-tagged protein abundance at 24 hours; and maximum 31.9.2 ± 5.7% at 72 hours; mean ± s.d.; Figure 5c).

Compared to the Bulk MPs, the ^13^C-protein relative abundance was higher at every time point in both the BONCAT^+^ FACS MP samples (10.0 ± 0.2, 4.6 ± 0.3, 10.0 ± 0.1, 8.4 ± 0.2-fold increases at 24, 48, 72, and 96 hours respectively) and BONCAT^+^ Pulldown MP samples (32.3 ± 0.2 and 4.0 ± 0.2-fold increases at 24 and 72 hours respectively) (Figure 5b). The ^13^C-protein relative abundance in the BONCAT^+^ FACS MPs increased continuously between 24 and 72 hours (maximum ^13^C-protein relative abundance 14.0 ± 1.1%; mean ± s.d.) before falling slightly at 96 hours (Figure 5b) and was attributed primarily to *Natronincolaceae_JAAZJW01_1* (minimum 85.0 ± 8.1% of ^13^C-tagged protein abundance at 72 hours; and maximum 94.1 ± 16.9% at 24 hours) (Figure 5c). The BONCAT^+^ Pulldown MPs showed a different trend over time (13.4 ± 1.4% ^13^C-protein relative abundance at 24 hours falling to 5.6 ± 0.6% at 72 hours) (Figure 5b), but were similarly dominated by *Natronincolaceae_JAAZJW01_1* (35.2 ± 7.7% of ^13^C-tagged protein abundance at 24 hours and 68.8 ± 10.0% at 72 hours) (Figure 5c).

Both the BONCAT^−^ FACS MPs and the HPG^−^ Pulldown MPs (e.g. negative controls) contained a similar ^13^C-protein relative abundance as the Bulk MPs (Figure 5b). The ^13^C-tagged protein abundance in the BONCAT^−^ FACS MPs was dominated by two MAGs from genus *Syntrophaceticus* as well as *Methanoculleaceae_Methanoculleus_5,* while that in the HPG^−^ Pulldown MPs was largely attributed to *Methanotrichaceae_Methanothrix_1* (Supplemental Figure S10). The *Syntrophaceticus* MAGs therefore incorporated ^13^C-acetate, but were less active than *Natronincolaceae_JAAZJW01_1* based on their lower ^13^C signal in the Bulk MPs and BONCAT^+^ MPs (Supplemental Figure S10).

Proteins from several MAGs were enriched in BONCAT-based MPs, but lacked significant ^13^C-labeling. *Pelotomaculaceae_DTU098_1* and *Pelotomaculaceae_Pelotomaculum_C_1* each had elevated abundance in the BONCAT^+^ FACS MPs and BONCAT^+^ FACS mini-MGs, but incorporated very little ^13^C-labeled acetate (maximum 4.1 ± 0.7% and 1.5 ± 0.4% of ^13^C-protein abundance at any time point, respectively). This phenomenon can be explained by previous reports of the *Pelotomaculaceae* family performing syntrophic propionate oxidation (SPO)^34–36^. In agreement with this, proteins related to the methylmalonyl-CoA pathway used for propionate oxidation were among the highest expressed in *Pelotomaculaceae_DTU098_1* (Supplemental Figure S12), and this organism has been previously proposed to perform SPO^37^. *Pelotomaculaceae_Pelotomaculum_C_1* did not encode a recognizable methylmalonyl-CoA pathway, but it did contain five separate propionate CoA-transferase genes (three of which had detectable protein expression), and multiple genes indicative of syntrophic behavior, such as heterodisulfide reductase^38^ and carbon monoxide dehydrogenase^39^ (Supplemental Figure S12).

We also observed differences in BONCAT labelling between protein Pulldown MPs and FACS MPs for certain organisms, and in particular methanogenic archaea. For instance, both *Methanotrichaceae_Methanothrix_1* and *Methanosarcinaceae_Methanosarcina_2* were metabolically active, as evidenced by elevated ^13^C-content in the Bulk MPs, but were excluded from the FACS MPs and FACS mini-metagenomes (e.g., no protein quantification and near-zero read mapping in mini-metagenomes) (Figure 5c). These organisms were detected, however, in the BONCAT Pulldown MPs, both in the BONCAT^+^ and HPG^−^ fractions (Figure 5c; Supplemental Figure S7). This observation further supports the notion that the filamentous and aggregated morphologies of these methanogens excluded their inclusion in FACS-based analyses, whereas they could be detected using the direct protein pulldown enrichment approach.

### Uncultured Natronincolaceae bacterium expressed a glycine-mediated pathway for acetate oxidation

The substantial incorporation of ^13^C by *Natronincolaceae_JAAZJW01_1* and its repeated enrichment in BONCAT-informed metagenomics and metaproteomics prompted further interrogation of its genome and protein expression profile. Despite its low abundance in the time-series metagenome sequencing data (maximum 0.412% relative abundance across twelve months, *n* = 8; Supplemental Table S3), a circular, high-quality draft MAG for this organism was recovered from the long-read sequencing assembly, with 98.8% completeness and 1.6% contamination (Supplemental Table S2). Genome annotation and reconstruction revealed the presence of genes relevant to SAO and related energy conservation machinery (Figure 6a). While this MAG did not encode the CODH/ACS complex necessary for the oxidative Wood-Ljungdahl pathway, it expressed enzymes from the glycine cleavage system relevant to the reductive glycine pathway^40^. Given the evidence for acetate oxidation in the SIP microcosms (Figure 3b), it is expected that this organism employs the reverse pathway, hereby termed the oxidative glycine pathway (OGP).

**Figure 6:**
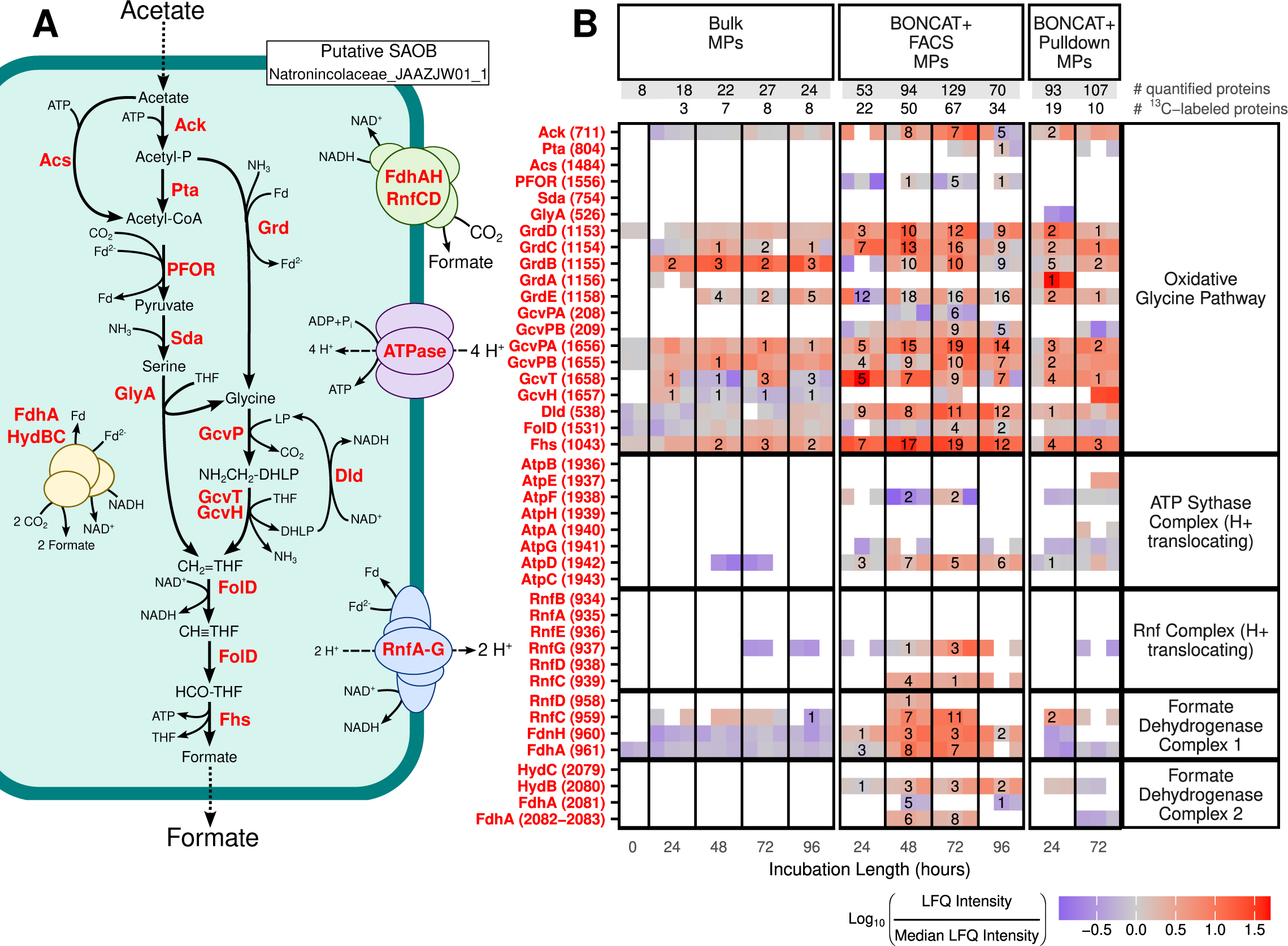
Metabolism of acetate oxidation in *Natronincolaceae_JAAZJW01_1* (i.e., *Syntrophacetatiphaga salishiae*). (A) Illustration of the oxidative glycine pathway including a non-exhaustive set of energy conservation mechanisms (B) Detection of proteins related to acetate metabolism in *Natronincolaceae_JAAZJW01_1*. Relative abundance values for each protein were rescaled by dividing by the median relative abundance among proteins from this MAG in each sample. Numbers indicate how many unique peptides from the given protein were ^13^C-labeled in any of the ^13^C-incubation samples (n = 3) at the given time point. Protein abbreviations are as follows: Ack = Acetate kinase, Acs = Acetyl-CoA synthetase, Dld = Dihydrolipoyl dehydrogenase, Fdh = Formate dehydrogenase, Fhs = Formate-tetrahydrofolate ligase, FolD = Methylenetetrahydrofolate cyclohydrolase, GcvH = Glycine cleavage system H protein, GcvP = Glycine cleavage system P protein, GcvT = Glycine cleavage system T protein, GlyA = Glycine hydroxymethyltransferase, Grd = Glycine reductase complex, Hyd = Bifurcating [FeFe] hydrogenase, PFOR = Pyruvate:ferredoxin oxidoreductase, Pta = Phosphate acetyltransferase, Sda = Serine deaminase, Rnf = Na^+^-translocating ferredoxin:NAD^+^ oxidoreductase complex.

The OGP involves the conversion of acetate into glycine by two possible routes: (1) acetate is converted to acetyl-P via acetate kinase (Ack) coupled to acetyl-CoA formation by phosphoacetyltransferase (Pta), or acetyl-CoA is directly generated from acetate by acetyl-CoA synthetase (Acs), followed by conversion of acetyl-CoA to pyruvate via pyruvate:ferredoxin oxidoreductase (PFOR), amination of pyruvate to serine by serine dehydratase (Sda), and conversion of serine into glycine by glycine hydroxymethyltransferase (GlyA); (2) acetate is converted to acetyl-P by Ack, and acetyl-P is oxidized to glycine by the glycine reductase complex (Grd) (Figure 6a). Glycine is then converted by the glycine cleavage system (GcvP, GcvT, GcvH, Dld) into 5,10-methylene-tetrahydrofolate, which is used by methylenetetrahydrofolate dehydrogenase (FolD) to form N^10^-formyltetrahydrofolate that is finally converted into formate by formate-tetrahydrofolate ligase (Fhs). OGP-related enzymes accounted for a large fraction of ^13^C-labeled peptides from *Natronincolaceae_JAAZJW01_1* in Bulk MPs (23 of the 25 peptides, *n* = 14), BONCAT+ Pulldown MPs (30 of 48 peptides, *n* = 6), and BONCAT^+^ FACS MPs (186 of the 524 peptides, *n* = 12) (Figure 6b). Also present in the *Natronincolaceae_JAAZJW01_1* genome were genes encoding multiple enzyme complexes related to energy conservation relevant to syntrophic organisms. Among those with protein expression data observed for at least two of their sub-components were a proton-translocating ferredoxin:NAD^+^ oxidoreductase (Rnf) complex, a proton-translocating ATP synthase, and two formate dehydrogenase complexes (Figure 6a). The BONCAT-informed metaproteomic enrichment methods provided a higher level of evidence for the activity of the OGP versus the conventional Bulk MPs workflow via the detection of more enzymes in the OGP pathway and energy conservation, along with detection of more ^13^C-labeled peptides from these enzymes (Figure 6b). Despite the greater detection of metabolic enzymes from *Natronincolaceae_JAAZJW01_1* with BONCAT-enabled MPs, expression of Sda by this organism was not observed in any metaproteome, and PFOR, Pta, and GlyA showed weak expression in the BONCAT-FACS MPs. This observation suggests that acetate was primarily converted to glycine via the Grd complex, which had high protein expression in the BONCAT-enabled MPs (Fig. 6b). To further investigate the possible activity of Ack/Grd over Pta/PFOR/Sda/GlyA, possible flux distributions were predicted for growth of *Natronincolaceae_JAAZJW01_1* on acetate using parsimonious flux balance analysis constrained by the amount of ^13^C protein produced (Figure 6; Figures 7 and 8). This metabolic model of *Natronincolaceae_JAAZJW01_1* predicted that all glycine was formed by Ack/Grd rather than Pta/PFOR/Sda/GlyA, supporting the proteomic expression levels of these alternative pathways (Figures 7 and 8).

**Figure 7.**
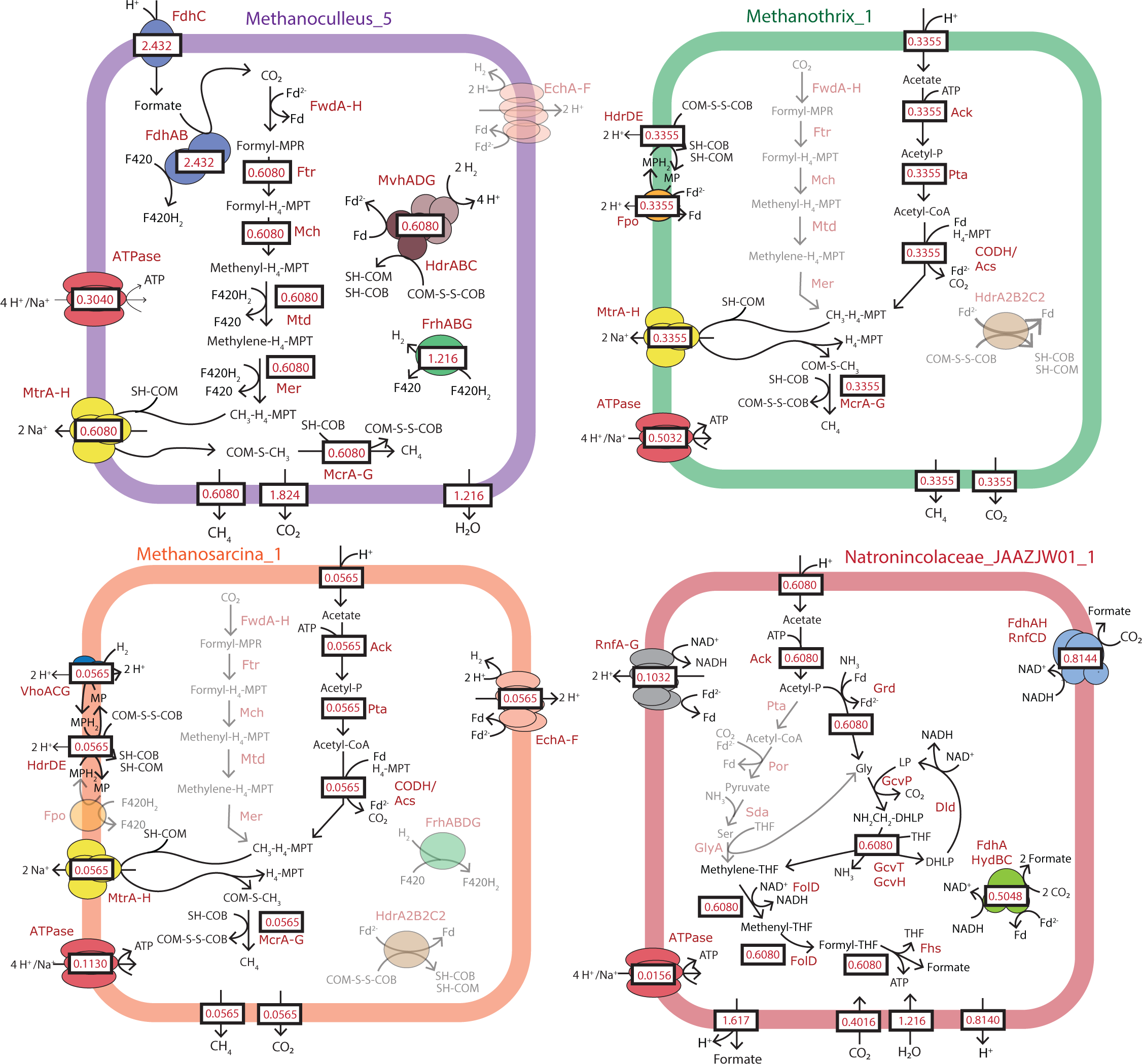
Metabolic fluxes predicted for the five most ^13^C labeled members at 24 hr. Model fluxes are based on consumption of one mol of acetate, and thus represent the stoichiometry across the metabolic networks at the discrete time point of 24 hr of the SIP incubation. Values were predicted with parsimonious flux balance analysis while constraining each organism’s ATP production to be proportional to the mass of ^13^C labeled proteins at 24 hr. Ack = Acetate kinase, Acs = Acetyl-CoA synthetase, CODH = carbon monoxide dehydrogenase, Dld = Dihydrolipoyl dehydrogenase, Ech = Energy-converting hydrogenase, Fdh = Formate dehydrogenase, Fhs = Formate-tetrahydrofolate ligase, FolD = Methylenetetrahydrofolate cyclohydrolase, Fpo = F_420_H_2_ dehydrogenase, Frh = F_420_-reducing hydrogenase, Ftr = Formylmethanofuran:tetrahyromethanopterin formyltransferase, Fwd = Formylmethanofuran dehydrogenase, GcvH = Glycine cleavage system H protein, GcvP = Glycine cleavage system P protein, GcvT = Glycine cleavage system T protein, GlyA = Glycine hydroxymethyltransferase, Grd = Glycine reductase complex, Hdr = heterodisulfide reductase, Hyd = Bifurcating [FeFe] hydrogenase, Mch = Methenyltetrahydromethanopterin cyclohydrolase, Mcr = Methyl-CoM reductase, Mer = 5,10-Methylenetetrahydromethanopterin reductase, Met = 5,10-Methylenetetrahydrofolate reductase, Mtd = F_420_-dependent methylenetetrahydromethanopterin dehydrogenase, Mtr = 5-Tetrahydromethanopterin:CoM-S-methyltransferase, Mvh = F_420_-nonreducing hydrogenase, PFOR = Pyruvate:ferredoxin oxidoreductase, Pta = Phosphate acetyltransferase, Sda = Serine deaminase, Rnf = Na^+^-translocating ferredoxin:NAD^+^ oxidoreductase complex, Vho = F_420_-nonreducing hydrogenase.

**Figure 8.**
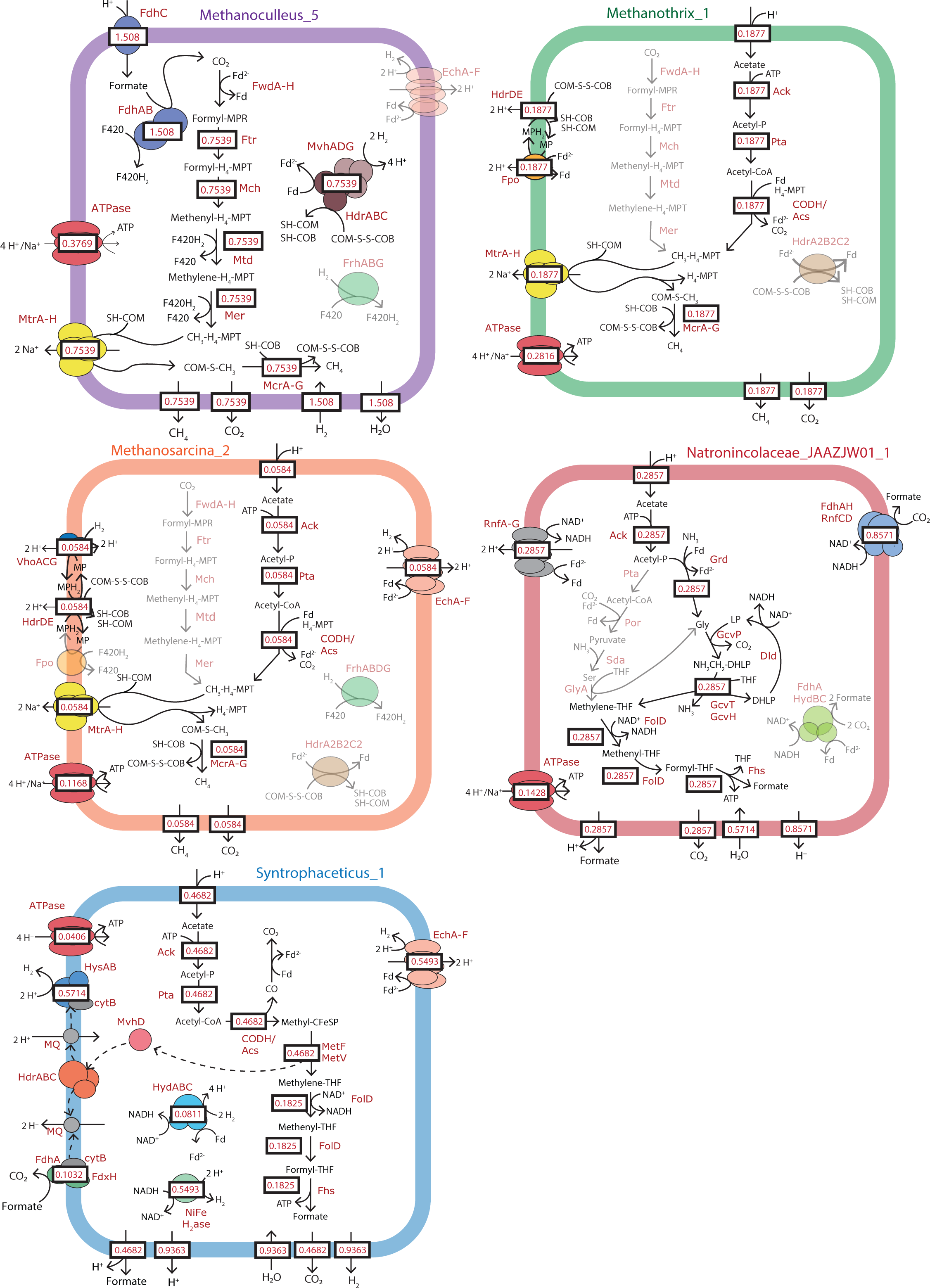
Metabolic fluxes predicted for the five most ^13^C labeled members at 72 hr. Model fluxes are based on consumption of one mol of acetate, and thus represent the stoichiometry across the metabolic networks at the discrete time point of 72 hr of the SIP incubation. Values were predicted with parsimonious flux balance analysis while constraining each organism’s ATP production to be proportional to the mass of ^13^C labeled proteins at 72 hr. Ack = Acetate kinase, Acs = Acetyl-CoA synthetase, CODH = carbon monoxide dehydrogenase, Dld = Dihydrolipoyl dehydrogenase, Ech = Energy-converting hydrogenase, Fdh = Formate dehydrogenase, Fhs = Formate-tetrahydrofolate ligase, FolD = Methylenetetrahydrofolate cyclohydrolase, Fpo = F_420_H_2_ dehydrogenase, Frh = F_420_-reducing hydrogenase, Ftr = Formylmethanofuran:tetrahyromethanopterin formyltransferase, Fwd = Formylmethanofuran dehydrogenase, GcvH = Glycine cleavage system H protein, GcvP = Glycine cleavage system P protein, GcvT = Glycine cleavage system T protein, GlyA = Glycine hydroxymethyltransferase, Grd = Glycine reductase complex, Hdr = heterodisulfide reductase, Hyd = Bifurcating [FeFe] hydrogenase, Mch = Methenyltetrahydromethanopterin cyclohydrolase, Mcr = Methyl-CoM reductase, Mer = 5,10-Methylenetetrahydromethanopterin reductase, Met = 5,10-Methylenetetrahydrofolate reductase, Mtd = F_420_-dependent methylenetetrahydromethanopterin dehydrogenase, Mtr = 5-Tetrahydromethanopterin:CoM-S-methyltransferase, Mvh = F_420_-nonreducing hydrogenase, PFOR = Pyruvate:ferredoxin oxidoreductase, Pta = Phosphate acetyltransferase, Sda = Serine deaminase, Rnf = Na^+^-translocating ferredoxin:NAD^+^ oxidoreductase complex, Vho = F_420_-nonreducing hydrogenase.

Expression of the OGP by *Natronincolaceae_JAAZJW01_1* would oxidize acetate into formate by two separate formate dehydrogenase enzyme complexes, a membrane bound FdhAH-RnfCD complex using NADH as an electron donor to produce formate and an electron bifurcating FdhA-HydBC that accepts electrons from both NADH and reduced ferredoxin (Figure 6a). As this metabolism is predicted to be energetically unfavorable under standard conditions (Supplemental Tables S8 and S9), formate would need to be maintained at low concentrations to create favorable thermodynamics for this metabolism^9^. The likely syntrophic partner of *Natronincolaceae_JAAZJW01_1* was the archaeon *Methanoculleaceae_Methanoculleus_5*, which accounted for more than 50% of the total protein abundance at every time point in the Bulk MPs (Figure 4b) and was among the highest ^13^C-labeled MAGs in every sample (Figure 5c and Supplemental Figure S10). Relevant to its role as a syntrophic partner to *Natronincolaceae_JAAZJW01_1* was high expression of multiple formate dehydrogenase complexes, of which expression was not detected in any other archaeal MAG. This methanogen also encoded a formate transporter (*fdhC*) that is required for growth on formate in methanogens^41,42^ (Supplemental Figure S13). Parsimonious flux balance analysis predicted that *Methanoculleaceae_Methanoculleus_5* accounted for 60% and 75% of the methane production at 24 hr and 72 hr, respectively (Figures 7 and 8, Supplemental Tables S8 and S9). Based on the model-predicted yield of 2.43 mol formate per 0.6 mol acetate consumed by *Natronincolaceae_JAAZJW01_1* (Supplemental Table S8), a formate concentration of 3.4 μM would enable a free energy release of −30 kJ per mol of ATP produced for both *Natronicolaceae_JAAZJW01_1* and *Methanoculleus_5* (assuming acetate at 0.05 M and CH_4_ and CO_2_ at 0.5 atm), which is similar to what has been observed for ATP phosphorylation measured in cells performing acetogenesis^43^. Formate has been previously measured in SAO communities at low micromolar levels^42^, indicating the important role of methanogenic partners in overcoming the thermodynamic barrier of this metabolism by maintaining low concentrations of electron shuttles.

We also observed likely competition for acetate between *Natronincolaceae_JAAZJW01_1* and both an obligate acetoclastic methanogen (*Methanotrichaceae_Methanothrix_1*), and a mixotrophic methanogen encoding for the acetoclastic pathway (*Methanosarcinaceae_Methanosarcina_2*) (Figure 6; Figures 7 and 8). In the BONCAT^+^ Pulldown MPs, the relative abundance of ^13^C-labeled proteins from MAG *Methanotrichaceae_Methanothrix_1* declined precipitously between 24 and 72 hours (43.6 ± 4.8% and 11.1 ± 3.5% of total ^13^C-tagged protein abundance, respectively; mean ± s.d., *n* = 3; Figure 5c), which corresponded with a decline in total protein abundance for this MAG over the same time period (Figure 4b). Both the ^13^C-labeled and total protein abundance in the HPG^−^ Pulldown MPs also increased for *Methanotrichaceae_Methanothrix_1* by 72 hr (Supplemental Figures S7 and S10). This decrease in labeling and total protein abundance of *Methanotrichaceae_Methanothrix_1* occurred in concert with the increase in methane generation via the SAO pathway predicted by gas-phase metabolomics (Figure 3b), indicating that *Methanotrichaceae_Methanothrix_1* likely was more active within the first 24 hours of the SIP incubation and had a decreasing activity as acetate was consumed. The community scale metabolic model corroborated this observation, predicting that 34% of the acetate flux went to *Methanotrichaceae_Methanothrix_1* at 24 hr, which decreased to 19% at 72 hr (Figures 7 and 8). *Methanosarcinaceae_Methanosarcina_2* was primarily detected in the Bulk MPs, and had a relatively stable protein relative abundance over time with a peak in ^13^C-protein relative abundance at 48 hours (Figure 5c). This methanogen encoded complete pathways for methane production from a wide variety of substrates (including H_2_/CO_2_, formate, methanol, acetate, and methylated amines) but showed the highest expression of proteins involved in the methanogenesis from acetate, methanol, and methylated amines (Supplemental Figure S14). It is therefore possible that the lower relative ^13^C labeling of *Methanosarcinaceae_Methanosarcina_2* could have been due to co-growth on acetate and (unlabeled) methanol and/or methylated amines during the SIP incubations. The community scale metabolic model supported this notion by predicting only 6% acetate flux went to *Methanosarcinaceae_Methanosarcina_2*, and was stable between 24 hr to 72 hr (Figures 7 and 8). Despite the lower fluxes of acetate for these two acetoclastic methanogens relative to *Natronincolaceae_JAAZJW01_1*, their predicted ATP yields were higher (Supplemental Tables S8 and S9). This suggests that *Natronincolaceae_JAAZJW01_1* could have outcompeted these organisms due to faster kinetics, or possibly its greater tolerance to the elevated free ammonia levels in this AD ecosystem^44^ (Supplemental Figure S15).

In addition to the two acetoclastic methanogens, we also observed possible competition for acetate by a putative SAOB, *Thermacetogeniaceae_Syntrophaceticus_1*, based on its ^13^C-labeling in the Bulk MPs and BONCAT-FACS MPs (Supplemental Figure S10) as well as its 99% ANI (Supplemental Table S2) to the isolated SAOB, *Syntrophaceticus schinkii*^45^. This organism encoded for an oxidative Wood-Ljungdahl pathway^46^ (Supplemental Figure S11), demonstrating that alternative metabolisms for SAO co-existed in this AD ecosystem. While metabolic modeling predicted no acetate flux for this organism at 24 hr (due to no ^13^C-labeled peptides observed), its acetate flux was predicted to increase to 46% by 72 hr (Figure 8). Modeling also suggested that the primary electron shuttle from *Thermacetogeniaceae_Syntrophaceticus_1* was H_2_, in contrast to *Natronincolaceae_JAAZJW01_1* that only produced formate (Supplemental Table S9; Figure 8). The H_2_ produced by *Thermacetogeniaceae_Syntrophaceticus_1* was predicted to help the growth of *Methanoculleaceae_Methanoculleus_5* between 24 hr and 72 hr, which could oxidize the H_2_ to reduce ferredoxin rather than sacrifice F_420_H_2_ to produce H_2_ internally (Figures 7 and 8). Correspondingly, with *Thermacetogeniaceae_Syntrophaceticus_1* active at 72 hr, the model-predicted total microbial community ATP is 23% higher than the 24 hr scenario (0.58 vs. 0.71 mol/rxn, respectively; Supplemental Tables S8 and S9), suggesting a community–wide benefit of the coexisting SAO pathways.

Given the substantial ^13^C labeling of *Natronincolaceae_JAAZJW01_1* during growth on acetate in the SIP incubations, we suspected it could also play an important role in other AD systems worldwide. A global survey of publicly available metagenomes revealed that this organism has been predominantly observed in engineered anoxic environments, specifically landfills and anaerobic digesters, with a maximum relative abundance of 4.3% in such systems (Figure 9). As the circular genome of *Natronincolaceae_JAAZJW01_1* contained three copies of full-length 16S rRNA genes (Supplemental Table S2), we also leveraged the MiDAS database of 16S rRNA genes within AD systems^47^ to query its distribution and observed it was abundant globally in such systems, with a maximum relative abundance of 3.35% (Figure 9). Notably, it was detected more frequently and at higher relative abundances in mesophilic AD systems versus thermophilic. Interestingly, we found that the 16S rRNA gene(s) of *Natronincolaceae_JAAZJW01_1* matched at 100% identity to a previously observed 16S rRNA operational taxonomic unit, named *OTU_M_H_C* (GenBank accession: MG356789), that was enriched over 10-fold (from 2-3% to 26–44% relative abundance) in continuous methanogenic bioreactors fed with sodium acetate (7.5 g/L) at 37°C^48^. Based on that observation, this phylotype was “strongly suggested” as a novel SAOB candidate by Westerholm et al.^48^. Here, we provide direct evidence of acetate metabolism and ^13^C labeling of this organism, supporting the previous hypothesis of its SAOB behavior. Given its global distribution in engineered anoxic ecosystems, we expect that the circular genome we obtained for *Natronincolaceae_JAAZJW01_1* containing multiple copies of full-length 16S rRNA genes will further enable its molecular identification and continue to expand upon its ecophysiological role as an SAOB in such systems. Based on the widespread abundance of *Natronincolaceae_JAAZJW01_1* in AD systems worldwide and its observed role as an SAOB, we hereby propose it as a type strain according to the SeqCode Registry^49^, with the species name of *Syntrophacetatiphaga salishiae* (registered as: seqco.de/r:nx_72bs4; etymology available in Supporting Table S10).

**Figure 9.**
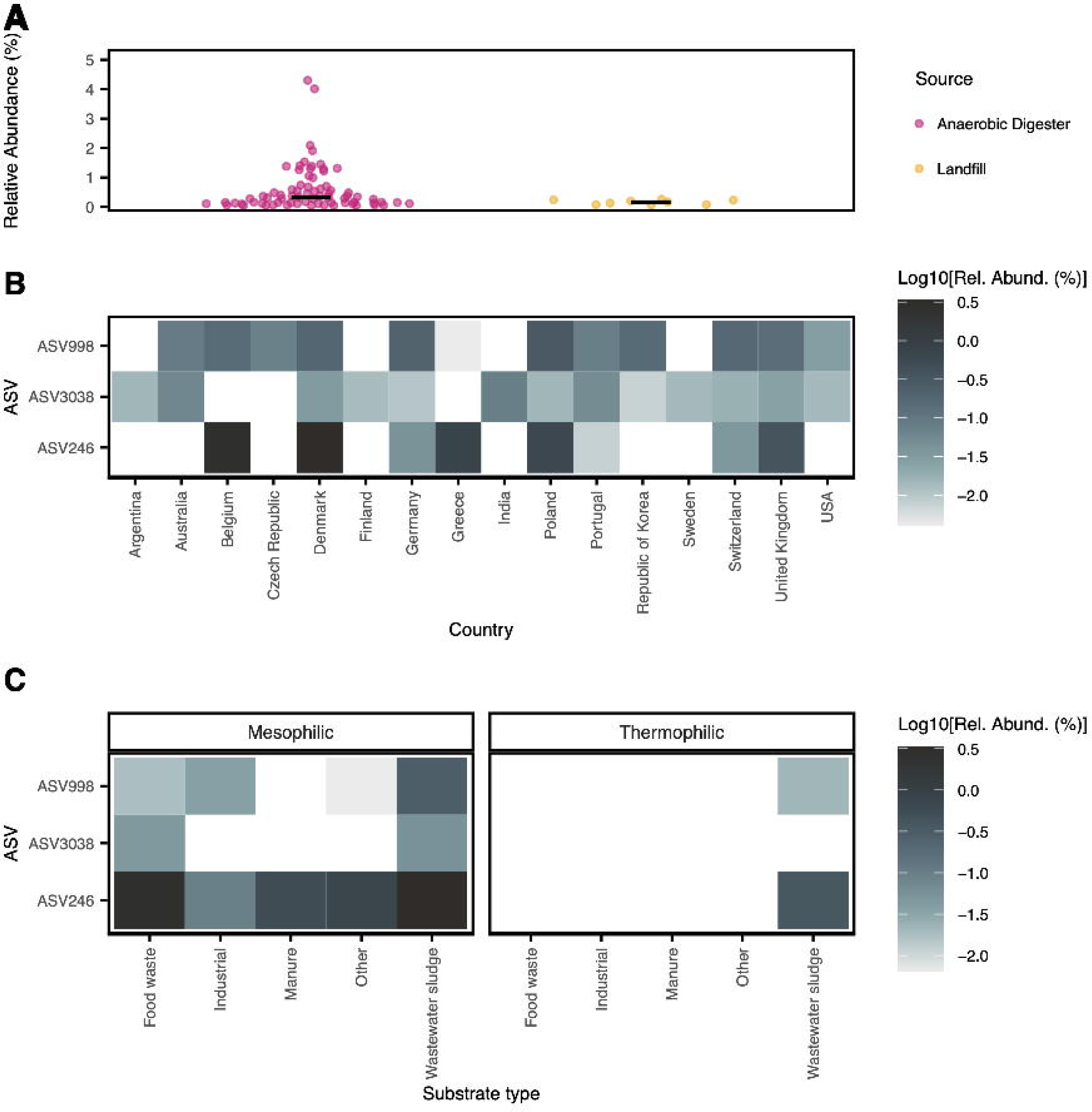
Global distribution of *Natronincolaceae_JAAZJW01_1* (i.e., *Syntrophacetatiphaga salishiae*). (A) Relative abundance in publicly available metagenomes on NCBI (database generated on 20 Feb, 2025) based on Sandpiper 1.0.1. metagenomic profiling. (B,C) Maximum observed relative abundance across anaerobic digesters organized by (B) country and (C) substrate type and temperature based on 16S rRNA gene amplicon sequencing data from the MiDAS global survey of anaerobic digesters^47^. V4 region amplicon sequence variants (ASVs) representing *Natronincolaceae_JAAZJW01_1* (>97% sequence identity) were identified by mapping the ASVs against the 16S rRNA genes from the MAG using usearch^116^ with the command `-usearch_global -maxrejects 0 -maxaccepts 0 -id 0 -top_hit_only`.

## Discussion

Through the combination of SIP, BONCAT, and metaproteogenomics, we identified an uncultured and previously undescribed microorganism, *Natronincolaceae_JAAZJW01_1* (hereby named *Syntrophacetatiphaga salishiae*), that was both relatively rare but significantly involved in acetate turnover within microcosm enrichments using biomass from a full-scale bioenergy facility. This species, affiliated with the phylum *Bacillota_A*, has been observed in other AD studies^25,48^ but never specifically characterized. In this study, long-read and short-read metagenomics recovered a circular genome for this organism containing multiple copies of 16S rRNA that can facilitate its future identification via amplicon sequencing approaches. It is notable that 16S rRNA gene sequences are currently not available for any MAG within this genus (*JAAZJW01*) in the Genome Taxonomy Database (GTDB; Release 226). BONCAT-based enrichment methods amplified the metagenomic and protein abundance of *Syntrophacetatiphaga salishiae* and revealed its high level of ^13^C-acetate incorporation in the SIP microcosms. The amplified information pertaining to its protein expression profile afforded by BONCAT metaproteomics allowed the detection of a complete metabolism for acetate oxidation to formate via the oxidative glycine pathway. While the involvement of the glycine cleavage system in SAO has been proposed for several taxa^50–52^, this work provides the first protein-level confirmation for the activity of the oxidative glycine pathway in an SAOB. This study therefore confirms the oxidative glycine pathway as an alternative to the oxidative Wood-Ljungdahl Pathway in organisms capable of SAO. Interestingly, metabolic modeling predicted a greater community ATP yield when both of these alternative pathways for SAO were active rather than just the former, which was attributable to their different electron shuttles (e.g., formate versus H_2_). The coexistence of these seemingly competitive SAO pathways within this anaerobic ecosystem may therefore reflect a community strategy to circumvent thermodynamic constraints by diversifying electron shuttles to methanogenic partners^42,53^.

This study demonstrated a novel approach for generating targeted metaproteomes of active microbial community members, without the need for prior identification or isolation. We employed BONCAT-enabled click chemistry to identify newly translated proteins in combination with ^13^C-based SIP to track carbon-utilization *in situ*. Two separate techniques were employed to isolate BONCAT-enriched proteins, either by cell sorting or direct protein pulldown. Overall, the two BONCAT-targeted metaproteomics approaches provided similar results, but each with a set of unique features and associated tradeoffs. The BONCAT Pulldown MPs targeted only proteins that were BONCAT-labeled (i.e., using biotin), while the FACS MPs targeted whole-cell pools that were BONCAT-labeled (i.e., using fluorescent dye). We therefore expected the BONCAT Pulldown MPs to contain a higher ^13^C abundance during SIP if translation and carbon-assimilation were coupled, as any unlabeled (i.e. ‘old’) cellular proteins would be removed. While this trend was observed at 24 hr of incubation with ^13^C-acetate, after 48 hr the relative abundance of ^13^C-proteins was lower in the Pulldown MPs compared FACS MPs, even though the ^13^C labeling ratio was similar. This observation suggests that the BONCAT Pulldown MPs could be impacted by potential carryover of ‘background’ non-BONCAT labeled proteins. On the other hand, a tradeoff associated with FACS-based enrichments was its sensitivity to aggregated morphologies. This was particularly observed for methanogenic archaea belonging to *Methanothrix* and *Methanosarcina*, of which we observed filamentous and tetrad-like cellular aggregates that were likely removed from the samples via filtration prior to FACS analysis. It should be noted that both genera have been successfully sorted with flow cytometry in other studies^54,55^, and thus this tradeoff should be surmountable. Additionally, the FACS-based BONCAT-SIP approach provided lower overall protein detection than Bulk MP; yet this was offset by its ability to identify more ^13^C peptides. These two approaches therefore offer alternative pathways to analyze proteins from active community members, and can be individually tailored to particular research questions or needs.

In conclusion, we demonstrated that targeting and capturing newly-translated proteins by BONCAT enhanced our ability to track ^13^C carbon into rare but active microorganisms during protein-SIP. The potential of functionally targeted metaproteomics may well be expanded by the use of other features besides BONCAT. For example, if the genome of a particular microorganism is available, near-pure-culture levels of proteomic information could be resolved using phylogenetic staining methods such as fluorescence *in situ* hybridization (FISH) in combination with activity labeling and cell sorting^56–58^. The higher-resolution of *in situ* physiologies provided by functionally targeted metaproteomics therefore opens many possibilities to deepen our understanding of the interconnected microbial metabolic networks driving matter and energy transformations within natural and engineered ecosystems.

## Online Methods

### Sample collection

The Surrey Biofuel Facility (SBF) is a closed-loop organic waste facility located in Surrey, British Columbia that treats organic waste originating from local households, industries, and commercial operations (Supplemental Figure S1). The SBF operates as a “dry” AD process, in which the incoming waste is shredded, piled into large anaerobic tunnels, and sprayed with a concentrated liquid solution of microbes–called “digestate”. Digestion occurs in the tunnels at 37°C over the course of approximately 30 days, and the liquid released by the digesting waste pile is collected and pumped to a continuously stirred tank reactor (CSTR) for re-application as digestate to later batches. Biogas composed primarily of methane produced in this step is collected and upgraded before it is injected into the natural gas grid and sold as renewable natural gas (RNG).

In order to observe changes in the microbial community inhabiting the SBF over time and generate a genomic database, liquid digestate from the continuous stirred-tank reactor (CSTR) at SBF was collected at various time points over the course of 19 months in sterile glass bottles and transported to the UBC campus in sealed, insulated containers. In the laboratory, nutrient analyses were performed immediately. Samples (2 x 1 mL) for DNA sequencing were centrifuged at 16,000 x g for 5 minutes to pellet cells. The supernatant was removed by decanting and the screw cap tubes containing the biomass pellets were stored at −80°C. Measurements of total ammonia nitrogen were performed per APHA method 4500-NH3-H on a flow injection analyzer (Lachat Instruments QuikChem FIA+ 8000 Series). Free ammonia concentrations were calculated from total ammonia nitrogen, temperature, and pH based on equations provided in Anthonisen et al., 1976^59^.

### Nucleic Acid Extraction

Biomass samples to be used for long-read DNA sequencing were thawed by adding 150 μL DNA/RNA Shield (Zymo Research, cat. R1100) and incubating at room temperature. DNA for long-read sequencing was extracted from each sample using the MagAttract PowerSoil Pro DNA kit (Qiagen, cat. 47119) following manufacturer instructions. The sample collected on 27 July 2021 was shipped on dry ice to the Joint Genome Institute (Berkeley, USA) for sequencing, while the samples collected on 13 January 2022 and 2 May 2022 were shipped on dry ice to the DNA Sequencing Center at Brigham Young University (Salt Lake City, USA). All samples were assessed by Femto Pulse (Agilent, cat. M5330AA) to have a mean fragment length greater than 10,000 bp. Each DNA sample was used to prepare a SMRTbell library (PacBio, cat. 102-141-700) and subsequently sequenced on a SMRT Cell 8M (PacBio, cat. 101-820-200) in a PacBio Sequel II in Circular Consensus Sequencing (CCS) mode.

Biomass samples to be used for short-read DNA sequencing were thawed at room temperature and DNA extraction was performed using the Qiagen Powersoil Pro MagAttract Kit (Qiagen, USA). The concentration of all DNA extracts was determined using the Qubit dsDNA BR Assay Kit on a Qubit 4 Fluorometer (Life Technologies, USA), while purity was assessed on a Nanodrop 2000 (Thermo Scientific, USA). Libraries were constructed for and sequenced (2×151) on the Illumina NovaSeq S4 platform. Sequencing reads were processed with BBMap v39.01 (https://sourceforge.net/projects/bbmap/) to trim ends and remove low-quality and contaminant reads.

Details of all metagenome sequencing runs, including NCBI accession numbers, can be found in Supplemental Table S1.

### Metagenome assembly and analysis

A set of representative genomes was generated by combining and de-replicating genome bins retrieved from both long-read sequencing data and short-read sequencing data.

HiFi long-reads were assembled and binned using the mmlong2-lite v1.0.2 pipeline (https://github.com/Serka-M/mmlong2-lite). Briefly, the three HiFi sequencing samples were co-assembled using Flye v2.9.2^60^ in metagenome mode. All HiFi and Illumina sequencing reads were mapped to the assembly with minimap2 v2.26^61^ to inform differential coverage binning. Automatic binning was performed using MetaBAT2 v2.15^62^, SemiBin v1.5^63^, and GraphMB v0.1.5^64^ and the resulting bins were de-replicated with DAS_Tool v1.1.3^65^ upon quality assessment with CheckM2 v1.0.2^66^.

Reads from each short-read Illumina sequencing sample were corrected and normalized using bbcms v38.90 (a.k.a. BBNorm; https://sourceforge.net/projects/bbmap/) and assembled independently using metaSPAdes v3.15.2^67^. The uncorrected reads from all eight short-read sequencing samples were then mapped to each individual assembly using bowtie2 v2.5.1^68^ to inform differential coverage binning. Automated binning was performed for each assembly using MetaBAT2 v2.15^62^.

Multiple binning approaches were employed to ensure an exhaustive set of genomes was generated, which would provide the most complete set of proteins to be used for top-down proteomics. To reduce redundancy of genomes and proteins in the database, the bins were de-replicated by sequence identity and quality. Genome bins retrieved from the HiFi co-assembly and each of the eight Illumina assemblies were assessed with CheckM2 v1.0.2^66^ to determine completeness and contamination, and with Infernal v1.1.4^69^ to determine the presence of structural RNA genes. The superset of all genome bins was then clustered by nucleotide similarity using dRep v3.4.2^70^. In short, dRep initially performed a pairwise comparison of all bins with Mash v1.1.1 and an ANI threshold of 94% followed by secondary comparison within primary clusters with FastANI v1.33 and an ANI threshold of 98%. The “best” bin from each of the resulting secondary clusters was selected using a conditional scoring algorithm adapted from Parks, et. al. 2020^71^ to prioritize more contiguous bins (e.g. bins with fewer contigs) as well as high quality bins (i.e. completeness >90% and contamination <5%). If a bin had > 90% completeness and < 5% contamination, the score was calculated as:

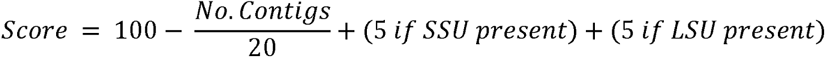

If a bin was less complete or more contaminated than this threshold, it was scored as follows:

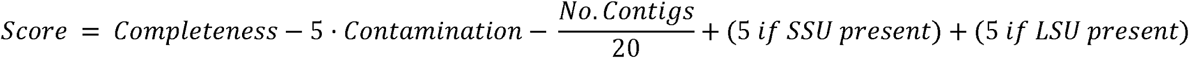

The de-replicated set of bins was taxonomically classified using GTDB-Tk v2.3.2^72^ and release 214 of the Genome Taxonomy DataBase (GTDB)^71^. Bins were renamed as [*Family*]_[*Genus*]_[sequential #] for interpretability. MAGs *Methanosarcinaceae_Methanosarcina_2 and Methanosarcinaceae_Methanosarcina_4* were later determined to be > 99.9% similar using skani v0.2.2^73^. As a result, the later MAG was removed from subsequent analysis. Details of all 912 medium and high-quality de-replicated MAGs, including their NCBI accession numbers, can be found in Supplemental Table S2.

DNA-based relative abundance of the genome bins over time was evaluated by mapping sequencing reads from each Illumina short-read time-series metagenome to the representative MAG using CoverM v0.7.0 (https://github.com/wwood/CoverM) and a minimum identity threshold of 98%.

In order to place our set of MAGs within the context of existing biogas metagenomic studies, the 466 high-quality MAGs from this study were compared to the 799 high-quality MAGs collected from a survey of 134 anaerobic digesters by Campanaro et. al.^25^. A non-overlapping set of high-quality MAGs was found using dRep with a secondary clustering threshold of 98%, and Anvi’o v8^74^ was used to perform phylogenomic placement using a concatenated alignment of ribosomal proteins according to the tree of life proposed by Hug et. al.^75^ The results were visualized with iTOL v6^76^.

Prodigal v2.6.3^77^ was used to predict open reading frames (ORFs) in the de-replicated MAGs from this study. The resulting protein sequences were functionally annotated using DRAM v1.5.0^78^ to perform HMM profile searches of the KOfam^79^ and Pfam v36.0^80^ databases. Proteins of particular interest were manually curated using NCBI services BLASTp and conserved domain search^81^. Finally, a non-redundant protein database was generated by clustering the ORFs at 100% identity using CD-hit v4.8.1^82^. This final protein database was appended to the “common repository of adventitious proteins” (https://www.thegpm.org/crap/) for the purpose of filtering contaminant proteins from the results.

Bins considered to be of particular interest based on protein expression in the microcosm experiment (the potential SAOB: *Natronincolaceae_JAAZJW001_1* and the most abundant MAGs from the main methanogenic genera: *Methanoculleaceae_Methanoculleus_5, Methanotrichaceae_Methanothrix_1*, and *Methanosarcinaceae_Methanosarcina_2*) were subsequently submitted to the MicroScope Annotation Platform^83^ for detailed functional protein prediction. The protein sequences called by MicroScope were associated with their corresponding sequences in the Prodigal-derived protein database using CD-hit-2D v4.8.1^82^ and existing functional annotations were superseded by the MicroScope annotations. Metabolic pathways in bins of particular interest were further investigated with Pathway Tools v27.5^84^ including the use of automated gap-filling^85^ to identify proteins not annotated by other methods.

### SIP Microcosm Incubation

SIP microcosms were established with SBF digestate and incubated with various labeling substrates to assess the active acetate-utilizing microbial community and its metabolic potential. Liquid digestate was collected from the CSTR at SBF in sterile glass bottles and placed in a sealed, insulated container during transportation. In the laboratory, the digestate was immediately submerged in a 37°C water bath. The bottles were then mixed thoroughly by manually shaking and 1.5 L of digestate was sieved to 1 mm under a constant flow of N_2_ gas into a sterile 2 L glass bottle. The sieved digestate was diluted 1:1 with anaerobic minimal media^86^ supplemented with 3.8 g/L ammonium chloride to match the average total ammonia nitrogen (TAN) concentration observed in the SBF (1.16 g TAN/L) (Supplemental Figure S15). Based on the reactor temperature of 37°C and the average measured TAN and pH (8.02) of the digestate, the free ammonia nitrogen (FAN) concentration in the microcosms was calculated to be 200 mg FAN/L (Supplemental Figure S15). The headspace was then flushed 5 times with alternating vacuum and N_2_ gas to ensure anoxic conditions. The 2 L bottle was then sealed with a butyl rubber septa and threaded cap, and a tedlar gas bag was attached to the septa to allow for pressure release. The inoculum was then incubated at 37°C for 72 hours with shaking at 125 rpm. This step is referred to as “pre-starvation” and served to remove excess carbon-containing compounds from the inoculum matrix. The purpose of the pre-starvation period was to exhaust any background growth substrates to maximize ^13^C incorporation during SIP, as well as to reduce any background translational activity to focus the BONCAT signal on acetate metabolizing organisms.

Under constant N_2_ flow and magnetic stirring, 41 mL of sieved, diluted digestate was transferred via a peristaltic pump into 61 mL microcosm serum bottles. Microcosm bottles were immediately sealed with rubber septa and aluminum crimps, and the headspace was flushed 5 times with N_2_ gas with alternating vacuum. The bottles were returned to the 37°C environment with shaking at 125 rpm for an additional 24 hours of ‘pre-starvation’ incubation.

Under constant N_2_ flow and magnetic stirring, microcosm bottles were opened individually and acetate and L-homopropargylglycine (HPG) were added to achieve the final concentrations of 60 mM and 50 μM, respectively, depending on the experimental condition (Supplemental Table S4). This concentration of HPG for the SIP microcosms was determined experimentally by incubating biomass from the SBF in the presence of 50 mM acetate and various levels of HPG and measuring the fraction of BONCAT positive cells via flow cytometry (Supplemental Figure S9). A total of 57 microcosms were established, including no-substrate controls, no-HPG controls, ^12^C-acetate controls, and ^13^C-acetate treatments (Supplemental Table S4). Universally labeled (^13^C-1,2-acetate) and methyl-C labeled (^13^C-2-acetate) sodium acetate were obtained from Cambridge Isotope Laboratories (Cat. No. CLM-440 and CLM-381; >99% atom purity) and HPG was obtained from Click Chemistry Tools (Cat. No. 1067). Acetate amendments were made using 1 mL of a 2.52 M sodium acetate solution. HPG amendments were made with 100 μL of a 21 mM HPG solution. The bottles were immediately resealed, flushed, and returned to the incubator with shaking at 125 rpm. Headspace pressures were equalized with the atmosphere using a needle attached to an empty gas bag. “Time 0” liquid samples were collected at this point, as described above.

Headspace pressure and methane content were measured in each microcosm bottle every 24 hours to monitor biogas production. A manometer (Traceable model 3462, VWR, USA) fitted with a 20 g needle was pierced through the rubber septum of each bottle to record the pressure. Methane content in the headspace was analyzed by manual collection of 6 μL via a gas syringe followed by injection into a gas chromatograph equipped with a flame ionization detector (HP5890 Series ii Gas Chromatograph, Hewlett Packard, USA and J&W HP-5 GC Column, Hewlett Packard, USA). The peak area was recorded and compared to a four-point calibration curve (0%, 10%, 50%, 80%, and 100%) prepared on the same day to determine the methane content. The biogas produced between sampling points was calculated using the ideal gas law, as described in Valero et. al. 2016^87^. After each measurement, the pressure in each headspace was equalized with the atmosphere using an empty gas bag prior to returning to the incubator. The gas volume was adjusted to standard temperature and pressure (STP) for conversion into molar quantities according to the ideal gas law. The “theoretical methane yield” was calculated as the number of moles of methane produced (minus methane produced in the no substrate control) divided by the number of moles of acetate added to the medium^87^. We also calculated the expected methane yield after accounting for a biomass yield of 0.02 gCOD_biomass_/gCOD_acetate_ for SAO consortia^42^ (Supplemental Figure S6).

At each sampling time point, 3 replicate microcosms from each of the primary experimental conditions were sacrificed under constant N_2_ flow for liquid sample collection. Samples (0.5 mL) for BONCAT-FACS analyses were combined 10:1 with GlyTE buffer (100mM Tris-HCl, 10mM EDTA, 55% v/v glycerol) for a final concentration of 5% glycerol as a cryopreservative^88^. They were sealed in screw cap tubes and flash frozen in a bath of 100% ethanol and dry ice. Once frozen they were stored at −80°C. Samples (8 mL) for bulk and protein-pulldown proteomics were centrifuged at 12,000 x g for 30 minutes in 15 mL conical tubes to pellet cells. The supernatant was removed by decanting and the pellets were stored at −80°C.

### IRMS and Methane Production Modeling

Gas-phase metabolomics using methyl labeled acetate was used to infer the relative contributions of acetoclastic methanogenesis versus SAO to methane formation^28,42^. The ^13^C/^12^C ratio of CO_2_ and CH_4_ in the microcosm headspace was calculated for three biological replicates incubated with 60mM ^13^C-2-acetate (i.e., methyl-C labeled) and three biological replicates incubated with 60 mM ^13^C-1,2-acetate (i.e., universally labeled) (Supplemental Table S4). At each sampling time point, headspace gas (500 µL) was injected into 3 mL exetainers (LabCo. 828W) previously flushed with N_2_. Compound specific isotope analysis (CSIA) was performed to analyze δ^13^C values of CO_2_ and CH_4_ in the samples by GC-C-IRMS (gas chromatograph combustion isotope ratio mass spectrometry). δ^13^C is defined as:

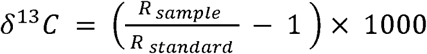

where: R represents the ratio of the heavy to light isotopes (^13^C/^12^C) for the sample and standard, with values denoted in δ-notation relative to Vienna Pee Dee Belemnite (VPDB) as per mil values (‰).

A Thermo Trace GC Ultra was coupled to a Thermo-Finnigan Delta V Plus IRMS using a GCC interface. Mixed gas molecules were separated using an Agilent J&W CarbonPLOT (Porous Layer Open Tubular) column (30 m x 0.53-mm ID x 3.00-µm d_f_), an oven temperature program and helium carrier gas (Supplemental Table S11). Sample gasses were injected into a splitless injection port using a Hamilton gastight syringe before being passed to the GC column to separate gas molecules. Depending on molecular concentration of CO_2_ and CH_4_, between 10 and 240 µL of sample gas were injected for each analysis. Samples were then passed through a micro combustion reactor packed with nickel and platinum wires at 940°C. To aid combustion, 1% O_2_ was added to the helium carrier gas stream prior to reactor introduction. The gas flow was then passed through a water removal column (Nafion membrane) before introduction into the IRMS for δ^13^C analysis of CO_2_ and CH_4_.

Samples were measured in triplicate across 4 time points (n = 24). Samples were analyzed using continuous flow IRMS, wherein CO_2_ reference gas pulses were introduced to the IRMS before and after each sample run. Samples were run interspersed between external standards of CO_2_ (Oztech, −10.4‰) and CH_4_ (LabCorp, −59.1‰). Instrument precision for the δ^13^C of external standards is < 0.4‰ for CH_4_ and 0.2‰ for CO_2_. The Atom % ^13^C fore each sample was calculated as:

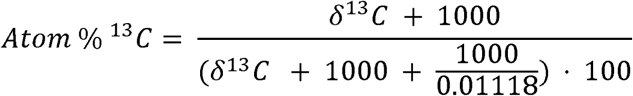

where the natural abundance of ^13^C is 0.01118.

Biological replicates (separate microcosms) and technical replicates (separate MS injections) were pooled within each time point for subsequent analysis. At each time point, the mean ^12^C-CO_2_ and ^12^C-CH_4_ atom fractions from the universally labeled condition were subtracted from those in the methyl-C labeled condition to obtain values corrected for ^12^C-CO_2_ and ^12^C-CH_4_ production from background total inorganic carbon in the inoculum. The percentage of CH_4_ produced via the SAO pathway was calculated using the model described previously^42^ (Supplemental Table S6). A Monte-Carlo simulation was used to assess the uncertainty of this prediction. For each timepoint, the ^13^C-CO_2_ to ^13^C-CH_4_ ratio in the methyl-C condition was sampled 10,000 times from a normal distribution given the mean and standard deviation of the measurements. Given the SAO% at each time point calculated by the above model as initial guesses, the *optimize()* function from the *stats* package in R^89^ was used to find values for the SAO% that minimized error across the simulated ^13^C-CO_2_ to ^13^C-CH_4_ values.

### BONCAT-FACS Mini-Metagenomes

We carried out BONCAT-enabled mini-metagenomics as a supplemental approach to view the active community, as well as to provide supporting evidence of the BONCAT-metaproteomics methods attempted in this study. Cryopreserved biomass from the microcosm incubation was shipped to the Joint Genome Institute in Berkeley, California, USA where whole-cell click chemistry, cell sorting, and DNA sequencing for the BONCAT mini-metagenomes were carried out. The click chemistry reaction was based on that of Hazenpichler and Orphan^90^ utilizing FAM Picoyl Azide as the fluorescent dye as it was observed that CalFluor488 caused inhibition of the multiple displacement amplification (MDA) reaction (Supplemental Figure S16). Accordingly, the cells were washed more stringently following the click reaction to account for the higher background signal of FAM Picoyl dye (Supplemental Table S12). Cells were counter-stained with SYTO59 and sorted using a BD Influx FACS (BD, USA). Details of the click reaction, sample handling, and sorting parameters and how they differ from the BONCAT-FACS metaproteome protocol may be found in Supplemental Table S12.

Optimization and calibration of the FACS was performed before each analysis using 3 μm Ultra Rainbow Fluorescent Particles (Spherotech, Lake Forest, IL). Particles were detected by triggering off the forward scatter (FSC) signal from a 488 nm laser equipped with a small particle detector. SYTO59-stained cells were detected using the red channel (excitation: 640 nm, emission: 655–685 nm) and BONCAT labeling with FAM Picolyl dye was measured on the green channel (excitation: 488 nm, emission: 510-550 nm). The BONCAT^+^ gate was drawn based on the fluorescence of two control samples from each time point that went through the click reaction but were not incubated with HPG. Less than 0.1% of the total cells in the control samples landed within the primary BONCAT+ gate. A second gate (i.e. Bright^++^) was drawn as a subset of the first that encompassed a tight population of larger and brighter cells within the BONCAT^+^ gate. Cells were sorted onto a 384-well plate, with 300 cells deposited per well. This included 16 wells of SYTO59^+^ cells, 16 wells of BONCAT^+^ cells, and 4 wells of Bright^++^ cells. A subset of wells from each sorted population was used for sequencing.

Following sorting, 5 µL MDA reactions were performed using Phi29 DNA Polymerase (Watchmaker Genomics, USA) as described previously^88^. DNA sequencing was performed on a NovaSeq (Illumina, USA) in 2 × 150 bp paired-end mode.

Sequencing reads from each mini-metagenome were mapped to the representative MAG set using CoverM v0.7.0 (https://github.com/wwood/CoverM) and a minimum identity threshold of 98%. The number of reads mapped to each MAG was used as count data for DESeq2 v1.38.3^91^ analysis. For this analysis, biological replicates were pooled for each time point and analyzed as a single condition for group comparisons. Briefly, size factors for each MAG were determined by calculating the median ratio of the read count in one sample to the geometric mean of read counts across all samples with non-zero values. Dispersion was calculated using a local regression. The difference between counts for each MAG between the cytometry gates were tested for significance by fitting to a negative binomial general linear model and calculation of the Wald statistic (without Cook’s cutoff). Significance was determined as having a Benjamini-Hochberg adjusted *P* value < 0.001, a log_2_-fold change > 2, and a relative abundance (as calculated by CoverM) greater than 0.5% in at least one sample.

### Bulk SIP-Metaproteomics

Untargeted metaproteomes were generated from the bulk digestate and microcosm incubations to provide datasets to which comparisons could be made using the BONCAT-enabled metaproteomes. Cell pellets (200 μL) from the SIP microcosms were transferred to a 1.5 mL Eppendorf Biopur tube with an equal amount of 0.1 mm Zirconia/Silica beads and 2× sample volume of 100 mM ammonium bicarbonate buffer, pH 8.0. Samples were lysed in a Bullet Blender (NextAdvance, Troy, NY) for 3 minutes at 4°C, speed of 8. The lysate was collected by puncturing the base of the 1.5 mL tube with a 26g needle and nesting that tube into a 15 mL centrifuge tube followed by centrifuging at 4°C, 4500 × g for 5 minutes. A BCA protein assay (ThermoFisher Scientific, Waltham, MA) was performed to determine approximate protein concentration of each sample and the volumes of the samples were normalized. Urea and dithiothreitol were added at 8 M and 5 mM (respectively) and the samples were incubated on a

Thermomixer (Eppendorf) for 30 minutes at 60°C with shaking at 850 rpm. Samples were diluted eight-fold with 1.14 mM CaCl_2_ in 50 mM ammonium bicarbonate and trypsin (Promega, Madison, WI) was added in a 1:50 (trypsin:protein, w:w) ratio. The samples were incubated for 3 hours at 37°C with shaking at 850 rpm, then frozen and stored at −70°C. Solid phase extraction (SPE) was performed on the samples using 1 mL/50 mg C18 columns (Phenomemex, Torrance, CA). Columns were conditioned with 3 mL of methanol followed by 2 mL of 0.1% trifluoroacetic acid (TFA) in water. The samples were applied to individual columns followed by 4 mL of 95:4.9:0.1 H_2_O:MeCN:TFA. Samples were eluted into 1.5 mL microcentrifuge tubes with 1 mL of 80:19.9:0.1 MeCN:H_2_O:TFA and concentrated in a vacuum concentrator to 75 μL. Another BCA protein assay was performed to obtain peptide concentration and a portion of each sample was diluted to 0.1 μg/μL for LC-MS/MS analysis.

The peptide solution was separated with a Dionex Ultimate HPLC system (Thermo Scientific, USA) and analyzed using a Q Exactive HF-X Orbitrap MS (Thermo Scientific, USA). Specifications of the LC separation and MS analysis are presented in Supplemental Table S13.

### BONCAT FACS SIP-Metaproteomics

One of the BONCAT-enabled metaproteomics approaches described in this study enriched whole cells using fluorescence-labelling of recently-translated proteins followed by FACS. Whole-cell click chemistry for the BONCAT-FACS metaproteomes was carried out at UBC in Vancouver, Canada where the microcosm incubation experiment took place. The click chemistry reaction was based on that of Hatzenpichler and Orphan^90^ but with CalFluor488 azide used as the fluorescent dye due to its lower background signal and tolerance for less stringent washing. The clicked cells were frozen at −80°C in 5% glycerol and shipped on dry ice to the Environmental Molecular Sciences Laboratory in Richland, Washington, USA where cell sorting and MS were carried out. As glycerol interferes with MS, the cells were washed briefly after thawing, a step not performed in the mini-metagenome method. Cells were counter-stained with SYTO59 and sorted using a BD Influx FACS (BD, USA). Details of the click reaction, sample handling, and sorting parameters and how they differ from the BONCAT-FACS mini-metagenome protocol may be found in Supplemental Table S12.

Optimization and calibration of the FACS was performed before each analysis using 3 μm Ultra Rainbow Fluorescent Particles (Spherotech, Lake Forest, IL). Particles were detected by triggering off the FSC signal from a 488 nm laser. Samples were excited with a 640 nm laser targeting SYTO59 to identify in-tact cells and gate out cellular debris. The singlet population of cells was identified using the Trigger Pulse Width versus the SYTO59 positive 670/30 nm signal. From the population passing this gate, BONCAT^+^ and BONCAT^−^ cells were identified by excitation with a 488 nm laser using the forward scatter detector versus the 520/15 nm detector. The BONCAT^+^ gate was drawn to exclude the background fluorescence of control samples that went through the click reaction but were not incubated with HPG. The BONCAT^−^ gate was drawn around an area that was more densely populated in the HPG^+^ vs. HPG^−^ samples but did not differentiate from the background fluorescence. Although this gate was named “BONCAT^−^” it is likely that some HPG-incorporating cells were missed by the stringent cutoff. We chose to prioritize reducing false positive results at the expense of a presumably elevated false negative rate. Approximately 50,000 BONCAT^+^ and BONCAT^−^ cells were separately sorted and stored at −80°C until processing by μPOTS.

Low-input proteomics using the microliter processing of trace samples in one pot (µPOTS) protocol, described by Xu et. al. 2019^85^, was employed to account for limited biomass available from sorting only 50,000 cells per metaproteome sample. The method is briefly outlined here. The sorted pools of cells (5 µL) were thawed and gently combined with 10 µL of 50 mM ammonium bicarbonate (ABC) buffer containing 0.1% *n*-Dodecyl β-d-maltoside and 2 mM dithiothreitol. The samples were transferred to a thermomixer at 90°C and mixed at 1000 rpm for one hour. After mixing, the samples were cooled to room temperature and then centrifuged at 1000 rpm for one minute. Next,1µL of an aklyation buffer containing 60mM chloroacetamide (CAA) in 50mM ABC buffer was added to each tube and was mixed at 1000 rpm at room temperature, in the dark for 30 minutes. For protein digestion, 10 μL of 50 mM ABC buffer containing 16 ng/μL trypsin and 8 ng/μL Lys-C was added to each tube and left to incubate at 37°C and 1500 rpm for 10 hours. To quench digestion, 1 μL of 5% formic acid was added to each tube. From each sample, 20 μL of peptide solution was removed and transferred to a polypropylene vial coated with 1% n-dodecyl β-D-maltoside (DDM) and stored at −20°C until analysis.

The peptide solution was separated with a Nano-Acquity dual-pumping ultra performance liquid chromatography (UPLC) system (Waters, USA) and analyzed using an Orbitrap Fusion Lumos Tribrid MS (Thermo Scientific, USA) outfitted with a high field asymmetric waveform ion mobility spectrometry (FAIMS) interface. Specifications of the LC separation and MS analysis are presented in Supplemental Table S13.

### BONCAT Protein Pulldown SIP-Metaproteomics

The second BONCAT-enabled metaproteomics approach described in this study enriched proteins (rather than whole cells) using click-chemistry and a bead-based biotin-streptavidin affinity assay. HPG labeled and unlabeled bioreactor samples were lyophilized overnight (~18 hours) until dry. Once dry, the samples were milled with a SPEX Genogrinder at 1750 rpm for 2 minutes. The ground samples were then transferred to fresh 1.7 mL Sorenson tubes and resuspended with 200 µL of water. Proteins were extracted using the MPLEx protocol^93^. After resuspension, a 1:5 (1 mL) volume of a 2:1 (v/v) mixture of chloroform and methanol was added to samples which were then horizontally vortexed for 10 minutes at 4°C. The samples were then placed in a −80°C freezer for 5 minutes. The samples were then sonicated for 30 seconds with a pulse of 1 second on and 1 second off at 60% amplitude on ice. After sonication, the samples were placed back in the −80°C freezer for 5 minutes. Sonication was repeated two more times for a total of three cycles. After three cycles of sonication and cooling, the samples were centrifuged at 4000 × g for 10 minutes at 4°C to separate the phases. Once the phases separated, the bottom layer (lipid phase) was removed using a glass serological pipette. Then, another 300 µL of ice-cold methanol was added to ensure that any remaining protein was washed off the tube walls. After methanol addition, the samples were centrifuged again at 4000 × g for 5 minutes at 4°C and the top layer (metabolite phase) was removed, leaving behind the protein pellet. The pellet was then allowed to dry on the benchtop for 15 minutes.

After the pellets were dry, 500 µL of 0.5% SDS in PBS was added to each sample, and they were sonicated for 30 seconds with a pulse of 1 second on 1 second off at 60% amplitude. The samples were then vortexed for 2 minutes and incubated for 30 minutes at 400 rpm and 50°C to solubilize the protein. The samples were then centrifuged at 4500 × g for 10 minutes at ambient temperature, and the supernatant was collected into a fresh 1.5 mL tube. The pellet was then subjected to another round of protein solubilization, and the supernatant was added to the first extraction for a total of 1 mL solubilized protein. The samples were then centrifuged at 8000 x g for 10 minutes at ambient temperature to remove any particulate humic substances, and the supernatant was transferred to a fresh 1.7 mL Sorenson tube.

To visualize and confirm the success of this method prior to mass spectrometry, a 50 µL aliquot from each sample was removed for SDS-PAGE fluorescence characterization (described in the next section). The remaining volumes were combined with 2.15 µL of 14 mM biotin azide, 5 µL of 500 mM sodium ascorbate, 5 µL of 280 mM THPTA, and 10 µL of 275 mM copper sulfate and incubated for 1 hour at 500 rpm and 37°C. After the click reaction, the samples were transferred to a 15 mL Olympus tube with 4 mL of ice-cold methanol and allowed to precipitate at −80 °C overnight. After methanol precipitation, the samples were centrifuged at 14,000 × g for 10 minutes at ambient temperature. Supernatant was then removed, and the pellet was allowed to dry upside down on the benchtop for 15 minutes or until completely dry. After the pellet was completely dry, 520 µL of 1.2% SDS in PBS was added to the samples and heated for 2 minutes at 95 °C and 500 rpm. After heating, the samples were then sonicated for 6 seconds with pulses of 1 second on 1 second off and 60% amplitude. After the solutions were completely homogenized, the samples were centrifuged at 14,000 × g for 5 minutes at 24 °C to pellet any insoluble material and protein concentrations were determined with a Pierce^TM^ BCA protein assay (Thermo Fisher Scientific, 23227).

The samples were then normalized with 1.2% SDS in 1× PBS to a final volume of 500 μL, to the lowest protein concentration amongst the samples determined by the BCA protein assay in 4 mL cryovials. All subsequent washes were performed using a vacuum manifold (VWR, PAA7231). Bio-Rad chromatography columns of 1.2 mL bed volume (Bio-Spin ^®^, 7326025) were washed twice with 1 mL 1× PBS and 100 μL of streptavidin-agarose resin (Thermo Scientific, 20353) were added to the columns. The beads were washed 2× with 1 mL 0.5% SDS in 1× PBS, 2× with 1 mL 5 M urea in 25 mM HEPES (prepared fresh, pH 8), and 4× with 1 mL 1× PBS. After washing, beads were transferred with two portions of 1 mL of 1× PBS to the 4 mL cryovials containing the normalized proteome samples. After transferring the beads, 500 µL of 1× PBS was added to the cryovials for a final approximate concentration of 0.2% SDS. The tubes were then rotated for 1 hour at 37°C.

After incubation, the protein-bound bead mixture was poured back into the columns. The cryovials were then rinsed with 1× PBS to transfer any remaining beads to the columns. This was repeated until there were no beads left in the vials. The protein bound beads were then washed 2× with 1 mL of 0.5% SDS in 1× PBS, 2× with 1 mL of 5 M urea in 25 mM HEPES (prepared fresh, pH 8), 2× with 1 mL of MilliQ water, 8× with 1 mL 1× PBS, and 4× with 1 mL of 25 mM HEPES (pH 8). After washing, the protein-bound beads were transferred to 1.7 mL low binding tubes (Sorenson BioScience, 11500-E) with 2× 350 μL 5 M urea in 25 mM HEPES (prepared fresh, pH 8). Then, 28 μL of 100 mM TCEP-HCL were added to each tube, and samples were reduced by incubating at 1,200 rpm, 37°C for 30 minutes. Samples were then alkylated by adding 28 μL of 200 mM iodoacetamide and shaking at 1,200 rpm, 50 °C, for 45 minutes in the dark. After reduction and alkylation, the protein-bound beads were transferred back to the corresponding column with 2× 500 μL volumes of 1× PBS. After transferring, the bound beads were washed 8× with 1 mL of 1× PBS and 4× with 1 mL of 25 mM ammonium bicarbonate (pH 8). After the final washing, the protein-bound beads were transferred to fresh 1.7 mL low binding tubes with 2× 500 μL of 25 mM ammonium bicarbonate (pH 8). The samples were then centrifuged at 10,500 × g for 5 minutes at 24 °C and the supernatant was discarded.

The protein-bound beads were resuspended in 200 μL of 25 mM ammonium bicarbonate (pH 8) and sufficient 0.25 μg/μL trypsin (Thermo Scientific, 90057) was added to achieve a 1:4000 (m/m) ratio of trypsin to protein (pre-enrichment). Protein digestion was conducted overnight at 1,200 rpm and 37°C. The following day, the samples were centrifuged at 10,500 × g for 5 minutes at room temperature. After centrifugation, exactly 200 μL of the supernatant was transferred to low retention individually wrapped 1.5 mL tubes (Fisher Scientific, 02-681-331). Then 150 μL of 25 mM ammonium bicarbonate (Fisher Scientific, 14-650-357) was added to the beads and mixed at 1,200 rpm and 37°C for 10 minutes. After mixing, the samples were centrifuged at 10,500 × g and 4°C for 5 minutes, and the resulting supernatant was added to the corresponding samples in the individually wrapped tubes. The tubes were covered with Breathe-Easier membranes (Electron Microscopy Sciences, 7053720) and allowed to dry completely. The samples were then reconstituted in 40 µL of 25 mM ammonium bicarbonate and heated at 37°C and 1000 rpm for 5 minutes. The samples were then transferred to polycarbonate centrifuge tubes (Beckman Coulter 343776) and spun at 53,000 rpm in a TLA 120.1 rotor at 4 °C for 20 minutes. After ultracentrifugation, 25 µL of the supernatant was transferred to MS vials.

The peptide solution was separated with a Nano-Acquity dual pumping UPLC system (Waters, USA) and analyzed using an Q Exactive Plus Orbitrap MS (Thermo Scientific, USA). Specifications of the LC separation and MS analysis are presented in Supplemental Table S13.

### SDS-PAGE

The 50 μL aliquot taken from the extracted, solubilized protein solution in the BONCAT-Protein Pulldown SIP-Metaproteomics workflow was used for SDS-PAGE fluorescence characterization with copper-catalyzed click chemistry (Supplemental Figure S17). Rhodamine-azide (30 μM final concentration), sodium ascorbate (2.5 mM), THPTA (1 mM), and copper sulfate (2 mM) were added to each aliquot. The reactions were incubated and protected from light for 1 hour at 37°C and 500 rpm. Following the incubation, excess rhodamine was removed by precipitating the protein using cold methanol (400 μL). Samples were placed in the −80 °C freezer overnight, followed by centrifugation at 10,500 × g at 4 °C for 5 minutes, after which the supernatant was discarded. The samples were allowed to air dry for 15 minutes. Probe-labeled, rhodamine-clicked proteins were resolubilized in water (16 μL), 2× SDS running buffer (20 μL) and 4 μL of 10× reducing agent (Invitrogen). The samples were sonicated for 2 seconds on, 1 second off with 60% amplitude using a probe sonicator. The samples were heated for 10 minutes at 95°C followed by spinning for 10-15 seconds in a tabletop centrifuge to pellet any remaining insoluble material. 20 μL of each sample was added to the wells of Novex™ WedgeWell™ 16% mini protein gels (Invitrogen). Gels were run at 135 V for 2.5 hours. Gels were imaged with a Typhoon FLA 9500 laser scanning imager (General Electric) and then fixed for 30 minutes in SYPRO fix solution (50% methanol with 7% acetic acid in MilliQ water). Fixed gels were rinsed with water, stained with GelCode blue for 2 hours to visualize total protein loading, and de-stained with water overnight. Stained gels were imaged using a Gel Doc EZ imager (BioRad Laboratories; Supplemental Figure S17).

### Metabolomics analysis

HPG unlabeled bioreactor samples were lyophilized overnight until dry. Once dry, 70 mg of the samples were then transferred to new 1.7 mL Sorenson tubes and extracted with a 4:1 methanol:water solution. The extracts were dried under vacuum and after this, they were chemically derivatized as previously reported^94^. Briefly, the extracts were derivatized by methoxyamination and trimethylsilyation (TMS), then the samples were analyzed by GC-MS. Samples were acquired in an Agilent GC 7890A using a HP-5MS column (30 m × 0.25 mm × 0.25 μm; Agilent Technologies, Santa Clara, CA) coupled with a single quadrupole MSD 5975C (Agilent Technologies). One microliter of sample was injected into a splitless port at constant temperature of 250°C. The GC temperature gradient started at 60°C, with a hold of temperature for 1 minute after injection, followed by increase to 325°C at a rate of 10°C/minute and a 10-minute hold at this temperature. A fatty acid methyl ester standard mix (C8-28) (Sigma-Aldrich) was analyzed in parallel as standard for retention time calibration.

GC-MS raw data files were processed using the Metabolite Detector software^95^. Retention indices (RI) of detected metabolites were calculated based on the analysis of a FAMEs mixture, followed by their chromatographic alignment across all analyses after deconvolution. Metabolites were initially identified by matching experimental spectra to a PNNL augmented version of Agilent GC-MS Metabolomics Library, containing spectra and validated retention indices for over 1200 metabolites. Then, the unknown peaks were additionally matched with the NIST20/Wiley11 GC-MS library. All metabolite identifications and quantification ions were validated and confirmed to reduce deconvolution errors during automated data-processing and to eliminate false identifications. Peak areas for identified metabolites were reported and subjected to statistical analysis for comparisons.

### Metaproteomics Data Processing

Mass spectra data were processed using quantms v1.3.1^96^ of the nf-core collection^97^ of workflows, utilizing reproducible software environments from the Bioconda^98^ and Biocontainers^99^ projects and tools from OpenMS v3.2.0^100^. The workflow was executed with Nextflow v24.04.4^101^. Both ^12^C-acetate (non-SIP) and ^13^C-acetate (SIP) samples from all metaproteomic methods were treated identically for peptide identification. Mass spectra were searched against the metagenome-derived protein database using both MSGF+ v2021.03.22^102^ and Comet v2023.01^103^. A target-decoy competition strategy was employed to estimate the rate of false positive peptide-spectrum matches (PSMs)^104^. Mass tolerances were set to 5 ppm for precursor (peptide) ions and 0.03 Da for fragment ions. Up to two missed cleavages were allowed and peptide termini were required to match the tryptic cleavage conditions fully. The precursor charge was limited to 2-4 and the peptide length was limited to 6-40. The PSMs were then rescored using the support-vector machine learning program Percolator v3.05.0^105^. The two search engine results were combined by retaining only the peptide sequence with the highest E-value for each spectrum and the consensus list of PSMs was filtered with a false discovery rate (FDR) threshold of 10% to reduce decoy and false positive matches.

Quantification of peptides was performed on the ^12^C-acetate (non-SIP) samples using the OpenMS tool, proteomicsLFQ. First, the PSM identity information extracted from the MS^2^ spectra was used to infer peptide quantity information from the MS^1^ spectra. Peptide intensities were scaled such that the median peptide intensity in every sample was equal. The peptides are then evaluated for “uniqueness” and used for protein inference. The quantification results were filtered with a protein FDR of 1% and a PSM FDR of 1%. Peptides which did not map uniquely to a protein but mapped to multiple proteins with shared characteristics (e.g. belonging to MAGs from the same phylum) were used for aggregated comparisons for the shared characteristics.

The peptide quantification data was then imported into R v4.2.0^106^ for processing and visualization with the tidyverse package v2.0.0^107^. The data was first filtered to remove erroneous or low confidence results. Peptides matching the cRAP database were considered contaminants and removed. Peptides that were only quantified in 1 of 3 replicates for a given time point and method were considered unreliable and removed. The remaining LFQ intensity was then summed and used as the “total intensity” of each sample. Relative biomass contributions of MAGs or higher taxonomic groups were computed as the sum of peptide intensities from all uniquely mapping peptides divided by the total intensity of the sample^29–31,108,109^.

Detection of isotopically labeled peptides was performed using MetaProSIP v2.3.0^110^ as implemented in OpenMS v2.8.0^100^ in a KNIME Analytics Platform v4.6.3^111^ workflow. MetaProSIP was chosen as it is capable of detecting background levels of ^13^C in unlabelled samples, as well as high levels (>10%) of isotopic labeling^16^. Briefly, MetaProSIP works by using information obtained from the paired, unlabeled condition (^12^C-acetate) to identify coeluting isotopologues in the labeled condition (^13^C-acetate). Using the list of monoisotopic peptides obtained from the peptide identification workflow described above, the workflow first performs retention time alignment between the ^13^C sample and all unlabeled samples from the same time point. For each identified peptide, MetaProSIP searches the mass spectrum of the ^13^C sample for all MS^1^ peaks starting at the mass of the monoisotopic peptide up to the mass of the peptide with every carbon substituted for ^13^C. The distribution of discovered peaks is compared to a theoretical distribution of isotopologue peaks and erroneous peaks are removed; the default parameters for this calculation were used. The intensity data is finally used to determine the global labeling ratio and relative isotope abundance of the peptide. Labeling ratio (LR) is defined as the ratio of labeled to unlabeled isotopologue intensities of a peptide. Relative isotope abundance (RIA) is defined as the percent labeling of possible atoms in the peptide and is an aggregate measure of all isotopologues observed^110^. Again, peptides which did not map uniquely to a protein but mapped to multiple proteins with shared characteristics (e.g. belonging to MAGs from the same phylum) were used for aggregated comparisons for the shared characteristics.

Relative isotopic protein abundance was calculated with the following formula:

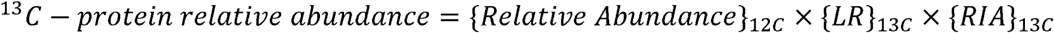

where: {*Relative Abundance*}_12C_ is the mean relative abundance of a protein across the unlabeled acetate incubation biological replicates, {*LR*}_13C_ is the mean peptide labeling ratio of all peptides observed for the protein across the universally-^13^C-labeled acetate incubation biological replicates, and {*RIA*}_13C_ is the mean peptide relative isotope abundance of all peptides observed for the protein across the universally-^13^C-labeled acetate incubation biological replicates. This number represents the ^13^C-protein abundance relative to all observed protein (g ^13^C-protein/g protein). To evaluate the accuracy of this measure, a bootstrapping procedure was used to estimate the 95% confidence interval of the difference in means between pairwise comparisons of the proteomic methods. The mean and standard deviation of the relative isotopic protein abundance were used to sample 3 simulated values from a normal distribution for two methods from which the difference between simulated sample means was determined. The procedure was repeated 10000 times and the 95% and 5% quantiles were taken to represent the 95% confidence interval.

### Fluorescence In Situ Hybridization (FISH)

Biomass samples (1 mL) for FISH were collected for fixation from the SIP microcosms at 96 hrs in 2 mL tubes and immediately placed on ice. Sample aliquots were then centrifuged at 15,000 x *g* to pellet the cells, and the supernatant was decanted. The cell pellet was then washed three times by addition of 500 µL PBS followed by centrifugation at 15,000 x *g* for 2 min and decanting the supernatant. The pellet was then resuspended in 500 µL PBS, then 500 µl of 4% paraformaldehyde (PFA) was added, the sample was inverted to mix, and was incubated at 4°C overnight. The sample was then centrifuged at 16,000 x *g* and the supernatant decanted. The cell pellet was then washed via resuspension in 1 mL of PBS followed by centrifugation at 16,000 x *g* for 3 minutes. After decanting the supernatant, the pellet was resuspended in 1 mL of 1:1 PBS:EtOH (100%) and stored at −20°C until FISH analysis.

For FISH, 5 µL of fixed sample was spotted onto a Teflon-coated slide and let air dry at room temperature. The slide was then dehydrated by flooding in a graded ethanol series (50%, 80%, and 96%, 3 min each). Then 9 µL of hybridization buffer was added to the sample followed by 1 µL of probe mix. The hybridization buffer contained (per 1 mL): 200 µL formamide, 600 µL Milli-Q water, 20 µL 1 M Tris/HCl, 180 µL 5 M NaCl, and 1 µL 10% SDS. The probe mix contained: Arch915: 5’-GTGCTCCCCCGCCAATTCCT-3’^112^ (Cy3; 30 ng/µL) targeting *Archaea*, LGC354A: 5’-TGGAAGATTCCCTACTGC-3’ + LGC354B: 5’-CGGAAGATTCCCTACTGC-3’ + LGC354C: 5’-CCGAAGATTCCCTACTGC-3’^113^ (Fluos; equimolar at 10 ng/µL each) targeting *Bacillota* (i.e. *Firmicutes*). Probe hybridization occurred at 46°C in a humid chamber for 90 minutes. The slide was then immersed in a wash buffer at 48°C for 10 minutes, followed by brief immersion in ice-cold MilliQ water and air dried. The wash buffer contained (per 50 mL): 2.15 mL 5 M NaCl, 1 mL 1 M Tris/HCl, 0.5 mL 0.5 M EDTA, 50 µL 10% SDS, and 46.3 mL MillQ water. Cells were counterstained with DAPI (10 µL of 1 ug/mL) for 3 minutes in the dark. Slides were then briefly rinsed with ice-cold MilliQ water followed by 96% EtOH, and air dried in the dark. Samples were then mounted with Citifluor (Electron Microscopy Sciences) and a coverslip and stored at 4°C until microcopy. Cells were analyzed by confocal microscopy with a Leica SP8 confocal laser scanning microscope (CLSM, Leica Microsystems) at the Centre for Microbiology and Environmental Systems Science of the University of Vienna. The CLSM instrument was equipped with a 405 nm diode laser and a white light laser for excitations between 470 and 670 nm. The Lightning Deconvolution Software package was used for image analysis (Leica Microsystems).

### Metabolic Modeling

Microbial communities were modeled using CobraPy^114^, with each organism represented by a separate compartment within the model. Pathways for central carbon and energy metabolism were reconstructed based on gene annotations for each MAG. The objective function for all simulations was to maximize the total ATP produced by the microbial community. Flux distributions were predicted using parsimonious flux balance analysis^115^ while constraining each organism’s ATP production to be proportional to the mass of ^13^C labeled proteins 24 hrs and 72 hrs after adding ^13^C labeled acetate. Model fluxes are based on consumption of one mol of acetate and thus represent the stoichiometry across the metabolic networks at two discrete time points. The metabolic models are available for download at the <u>OSF.io</u> repository: https://osf.io/u5vyc/?view_only=fde700d17bd74c5eb6a2d05ad73d33d5.

## Data Availability

All sequencing reads from the time series metagenomes, PacBio metagenomes, and mini-metagenomes and the 466 high-quality MAGs are available through the NCBI BioProject identifier: PRJNA1180090. Details of all metagenome samples, including individual NCBI accession numbers, can be found in Supplemental Table S1. NCBI accessions for individual MAGs from this study are provided in Supplemental Table S2. All proteomic data is available through the ProteomeXchange accession: PXD057167. Additional data and metadata used in the analysis can be found in the <u>OSF.io</u> repository: https://osf.io/u5vyc/?view_only=fde700d17bd74c5eb6a2d05ad73d33d5.

## Code Availability

Computational code used to analyze the metagenomic and metaproteomic data, conduct metabolic modeling, and generate the figures shown in this paper, are included in the OSF.io repository: https://osf.io/u5vyc/?view_only=fde700d17bd74c5eb6a2d05ad73d33d5.

## Acknowledgements

This work was performed under the auspices of the Natural Sciences and Engineering Research Council (NSERC) of Canada (Grant No. CRDPJ 543962-19 and RGPIN-2018-04585, both to R.M.Z.), Genome British Columbia (Grant No. SIP027 to R.M.Z. and S.J.H), the Canada Foundation for Innovation (CFI), and the US Department of Energy (DOE) Joint Genome Institute (JGI) and Environmental Molecular Science Laboratory (EMSL) Facilities Integrating Collaborations for User Science (JGI-EMSL FICUS) program (10.46936/fics.proj.2020.51515/60000205) and used resources at the JGI (https://ror.org/04xm1d337) and EMSL (https://ror.org/04rc0xn13) under Contract Nos. DE-AC02-05CH11231 (JGI) and DE-AC05-76RL01830 (EMSL). A portion of this research was supported under the EMSL user project award https://doi.org/10.46936/intm.proj.2022.60473/60008546, for developing technologies for and using capabilities operating at the Environmental Molecular Sciences Laboratory, a DOE Office of Science User Facility, sponsored by the Biological and Environmental Research program under Contract No. DE-AC05-76RL01830. The LABGeM (CEA/Genoscope & CNRS UMR8030), the France Génomique and French Bioinformatics Institute national infrastructures (funded as part of Investissement d’Avenir program managed by Agence Nationale pour la Recherche, contracts ANR-10-INBS-09 and ANR-11-INBS-0013) are acknowledged for support within the MicroScope annotation platform. We are grateful to Carrie Nicora for her advice and guidance with sample milling. We thank Ron Moore and Priscila Lalli for performing proteomics runs. We also want to acknowledge Markus Schmid for help with confocal microscopy. We also want to thank Maryna Kazak and Felicia Crozier for assisting with sample collection from the full-scale bioenergy facility.

## Author Contributions

R.M.Z and S.J.H conceived the research. S.F. and R.M.Z. processed the metagenomic and metaproteomic data and analyzed the results. S.F. and E.M. performed the microcosm incubation experiment. M.M. and K.W. performed the time series sample collection and advised on the microcosm incubation. M.K.D.D. performed the global distribution analysis based on 16S rRNA gene amplicon sequencing data from the MiDAS project. R.R.M. and D.G. designed and performed the mini-metagenome FACS protocol. W.C. performed the metagenome FACS protocol. L.J.G., C.J.K., L.H.G., H.M.O., and V.S.L designed and performed the BONCAT-protein pulldown assay and mass spectrometry. H.M.O performed protein extraction for the bulk metaproteome. J.T. and N.M. performed the bulk metabolomics extraction and untargeted metabolomics analysis by GC-MS. S.W. performed mass spectrometry for the bulk and BONCAT-FACS samples. S.B.L. performed IRMS analysis. N.T. and L.P-T. performed the µPOTS method. M.L. advised on the project formation. R.M.Z. and N.T. advised on the proteomic data analysis. S.F. and R.M.Z. drafted the manuscript. All authors discussed the results and commented on the manuscript.

## Competing Interests

With the exception of SJH, the authors declare no competing interests associated with this work. SJH is a co-founder of Koonkie Inc., a bioinformatics consulting company that designs and provides scalable algorithmic and data analytics solutions in the cloud. Koonkie Inc. was not involved in any aspect of this research.

## Notes

### Summary of Updates

The bacterial species name has been updated to "Syntrophacetatiphaga salishiae" to reflect the final endorsed naming in the SeqCode database.

## References

1. Briones, A. & Raskin, L. Diversity and dynamics of microbial communities in engineered environments and their implications for process stability. Curr. Opin. Biotechnol. 14, 270– 276 (2003).

2. Ducklow, H. Microbial services: challenges for microbial ecologists in a changing world. Aquat. Microb. Ecol. 53, 13–19 (2008).

3. Sampara, P., Lawson, C. E., Scarborough, M. J. & Ziels, R. M. Advancing environmental biotechnology with microbial community modeling rooted in functional ‘omics. Curr. Opin. Biotechnol. 88, 103165 (2024).

4. Hatzenpichler, R., Krukenberg, V., Spietz, R. L. & Jay, Z. J. Next-generation physiology approaches to study microbiome function at single cell level. Nat. Rev. Microbiol. 18, 241– 256 (2020).

5. Hua, Z.-S. et al. Ecological roles of dominant and rare prokaryotes in acid mine drainage revealed by metagenomics and metatranscriptomics. ISME J. 9, 1280–1294 (2015).

6. Jia, X., Dini-Andreote, F. & Salles, J. F. Unravelling the interplay of ecological processes structuring the bacterial rare biosphere. ISME Commun. 2, 96 (2022).

7. Dyksma, S., Jansen, L. & Gallert, C. Syntrophic acetate oxidation replaces acetoclastic methanogenesis during thermophilic digestion of biowaste. Microbiome 8, 105 (2020).

8. Lueders, T., Pommerenke, B. & Friedrich, M. W. Stable-Isotope Probing of Microorganisms Thriving at Thermodynamic Limits: Syntrophic Propionate Oxidation in Flooded Soil. Appl. Environ. Microbiol. 70, 5778–5786 (2004).

9. Schink, B. Energetics of syntrophic cooperation in methanogenic degradation. Microbiol. Mol. Biol. Rev. 61, 262–280 (1997).

10. McInerney, M. J., Sieber, J. R. & Gunsalus, R. P. Syntrophy in anaerobic global carbon cycles. Curr. Opin. Biotechnol. 20, 623–632 (2009).

11. Westerholm, M., Dolfing, J. & Schnürer, A. Growth Characteristics and Thermodynamics of Syntrophic Acetate Oxidizers. Environ. Sci. Technol. 53, 5512–5520 (2019).

12. Angelidaki, I. & Batstone, D. J. Anaerobic Digestion: Process. in Solid Waste Technology & Management (ed. Christensen, T. H.) 583–600 (John Wiley & Sons, Ltd, Chichester, UK, 2010). doi:10.1002/9780470666883.ch37.

13. Conrad, R. Contribution of hydrogen to methane production and control of hydrogen concentrations in methanogenic soils and sediments. FEMS Microbiol. Ecol. 28, 193–202 (1999).

14. Kleiner, M. et al. Assessing species biomass contributions in microbial communities via metaproteomics. Nat. Commun. 8, 1558 (2017).

15. Mosbæk, F. et al. Identification of syntrophic acetate-oxidizing bacteria in anaerobic digesters by combined protein-based stable isotope probing and metagenomics. ISME J. 10, 2405–2418 (2016).

16. Kleiner, M. et al. Ultra-sensitive isotope probing to quantify activity and substrate assimilation in microbiomes. Microbiome 11, 24 (2023).

17. Hatzenpichler, R., et al. *In situ* visualization of newly synthesized proteins in environmental microbes using amino acid tagging and click chemistry. Environ. Microbiol. 16, 2568–2590 (2014).

18. Couradeau, E. et al. Probing the active fraction of soil microbiomes using BONCAT-FACS. Nat. Commun. 10, 2770 (2019).

19. Hellwig, P. et al. Tracing active members in microbial communities by BONCAT and click chemistry-based enrichment of newly synthesized proteins. ISME Commun. 4, ycae153 (2024).

20. Madill, M. B. W., Luo, Y., Sampara, P., Ziels, R. M. & Gilbert, J. A. Activity-Based Cell Sorting Reveals Resistance of Functionally Degenerate Nitrospira during a Press Disturbance in Nitrifying Activated Sludge. mSystems vol. 6 e00712–21 (2021).

21. Scott, M. & Hwa, T. Shaping bacterial gene expression by physiological and proteome allocation constraints. Nat. Rev. Microbiol. 21, 327–342 (2023).

22. Lawson, C. E. et al. Rare taxa have potential to make metabolic contributions in enhanced biological phosphorus removal ecosystems. Environ. Microbiol. 17, 4979–4993 (2015).

23. Madsen, E. L. Identifying microorganisms responsible for ecologically significant biogeochemical processes. Nat. Rev. Microbiol. 3, 439–446 (2005).

24. The Genome Standards Consortium et al. Minimum information about a single amplified genome (MISAG) and a metagenome-assembled genome (MIMAG) of bacteria and archaea. Nat. Biotechnol. 35, 725–731 (2017).

25. Campanaro, S. et al. New insights from the biogas microbiome by comprehensive genome-resolved metagenomics of nearly 1600 species originating from multiple anaerobic digesters. Biotechnol. Biofuels 13, 25 (2020).

26. Wirth, R. et al. Inter-kingdom interactions and stability of methanogens revealed by machine-learning guided multi-omics analysis of industrial-scale biogas plants. ISME J. 17, 1326–1339 (2023).

27. Steward, K. F. et al. Metabolic Implications of Using BioOrthogonal Non-Canonical Amino Acid Tagging (BONCAT) for Tracking Protein Synthesis. Front. Microbiol. 11, (2020).

28. Mulat, D. G. et al. Quantifying Contribution of Synthrophic Acetate Oxidation to Methane Production in Thermophilic Anaerobic Reactors by Membrane Inlet Mass Spectrometry. Environ. Sci. Technol. 140130145609003 (2014) doi:10.1021/es403144e.

29. Wiśniewski, J. R. et al. Extensive quantitative remodeling of the proteome between normal colon tissue and adenocarcinoma. Mol. Syst. Biol. 8, 611 (2012).

30. Wiśniewski, J. R. & Rakus, D. Multi-enzyme digestion FASP and the ‘Total Protein Approach’-based absolute quantification of the Escherichia coli proteome. J. Proteomics 109, 322–331 (2014).

31. Delogu, F. et al. Integration of absolute multi-omics reveals dynamic protein-to-RNA ratios and metabolic interplay within mixed-domain microbiomes. Nat. Commun. 11, 4708 (2020).

32. Oren, A. The Family Methanotrichaceae. in The Prokaryotes (eds. Rosenberg, E., DeLong, E. F., Lory, S., Stackebrandt, E. & Thompson, F.) 297–306 (Springer Berlin Heidelberg, Berlin, Heidelberg, 2014). doi:10.1007/978-3-642-38954-2_277.

33. Conklin, A., Stensel, H. D. & Ferguson, J. Growth Kinetics and Competition Between Methanosarcina and Methanosaeta in Mesophilic Anaerobic Digestion. Water Environ. Res. 78, 486–496 (2006).

34. Hao, L. et al. Novel syntrophic bacteria in full-scale anaerobic digesters revealed by genome-centric metatranscriptomics. ISME J. 14, 906–918 (2020).

35. Singh, A., Schnürer, A., Dolfing, J. & Westerholm, M. Syntrophic entanglements for propionate and acetate oxidation under thermophilic and high-ammonia conditions. ISME J. 17, 1966–1978 (2023).

36. Westerholm, M., Calusinska, M. & Dolfing, J. Syntrophic propionate-oxidizing bacteria in methanogenic systems. FEMS Microbiol. Rev. 46, fuab057 (2022).

37. Campanaro, S., Treu, L., Kougias, P. G., Luo, G. & Angelidaki, I. Metagenomic binning reveals the functional roles of core abundant microorganisms in twelve full-scale biogas plants. Water Res. 140, 123–134 (2018).

38. Sedano Núñez, V. T., Boeren, S., Stams, A. J. M. & Plugge, C. M. Comparative proteome analysis of propionate degradation by *Syntrophobacter fumaroxidans* in pure culture and in coculture with methanogens. Environ. Microbiol. 20, 1842–1856 (2018).

39. Schuchmann, K. & Müller, V. Autotrophy at the thermodynamic limit of life: a model for energy conservation in acetogenic bacteria. Nat. Rev. Microbiol. 12, 809–821 (2014).

40. Sánchez-Andrea, I. et al. The reductive glycine pathway allows autotrophic growth of Desulfovibrio desulfuricans. Nat. Commun. 11, 5090 (2020).

41. Fink, C. et al. A Shuttle-Vector System Allows Heterologous Gene Expression in the Thermophilic Methanogen Methanothermobacter thermautotrophicus ΔH. mBio 12, e02766–21 (2021).

42. McDaniel, E. A. et al. Diverse electron carriers drive syntrophic interactions in an enriched anaerobic acetate-oxidizing consortium. ISME J. 17, 2326–2339 (2023).

43. Müller, V. & Hess, V. The Minimum Biological Energy Quantum. Front. Microbiol. 8, (2017).

44. Yi, Y. et al. Thermodynamic restrictions determine ammonia tolerance of methanogenic pathways in *Methanosarcina barkeri*. Water Res. 232, 119664 (2023).

45. Westerholm, M., Roos, S. & Schnürer, A. Syntrophaceticus schinkii gen. nov., sp. nov., an anaerobic, syntrophic acetate-oxidizing bacterium isolated from a mesophilic anaerobic filter. FEMS Microbiol. Lett. 309, 100–104 (2010).

46. Manzoor, S., Bongcam-Rudloff, E., Schnürer, A. & Müller, B. Genome-Guided Analysis and Whole Transcriptome Profiling of the Mesophilic Syntrophic Acetate Oxidising Bacterium Syntrophaceticus schinkii. PLOS ONE 11, e0166520 (2016).

47. Dueholm, M. K. D. et al. MiDAS 5: Global diversity of bacteria and archaea in anaerobic digesters. Nat. Commun. 15, 5361 (2024).

48. Westerholm, M., Müller, B., Singh, A., Karlsson Lindsjö, O. & Schnürer, A. Detection of novel syntrophic acetate-oxidizing bacteria from biogas processes by continuous acetate enrichment approaches. Microb. Biotechnol. 11, 680–693 (2018).

49. Hedlund, B. P. et al. SeqCode: a nomenclatural code for prokaryotes described from sequence data. Nat. Microbiol. 7, 1702–1708 (2022).

50. Kieft, B. et al. Genome-resolved correlation mapping links microbial community structure to metabolic interactions driving methane production from wastewater. Nat. Commun. 14, 5380 (2023).

51. Nobu, M. K. et al. Microbial dark matter ecogenomics reveals complex synergistic networks in a methanogenic bioreactor. ISME J. 9, 1710–1722 (2015).

52. Zhu, X. et al. Metabolic dependencies govern microbial syntrophies during methanogenesis in an anaerobic digestion ecosystem. Microbiome 8, 22 (2020).

53. Nobu, M. K. et al. Catabolism and interactions of uncultured organisms shaped by eco-thermodynamics in methanogenic bioprocesses. Microbiome 8, 111 (2020).

54. Lambrecht, J. et al. Flow cytometric quantification, sorting and sequencing of methanogenic archaea based on F420 autofluorescence. Microb. Cell Factories 16, 1–15 (2017).

55. McKay, L. J. et al. Activity-based, genome-resolved metagenomics uncovers key populations and pathways involved in subsurface conversions of coal to methane. ISME J. 16, 915–926 (2022).

56. Amann, R. & Fuchs, B. M. Single-cell identification in microbial communities by improved fluorescence in situ hybridization techniques. Nat. Rev. Microbiol. 6, 339–348 (2008).

57. Yilmaz, S., Haroon, M. F., Rabkin, B. A., Tyson, G. W. & Hugenholtz, P. Fixation-free fluorescence *in situ* hybridization for targeted enrichment of microbial populations. ISME J. 4, 1352–1356 (2010).

58. Harris, R. L. et al. FISH-TAMB, a Fixation-Free mRNA Fluorescent Labeling Technique to Target Transcriptionally Active Members in Microbial Communities. Microb. Ecol. 84, 182– 197 (2022).

59. Anthonisen, A. C., Loehr, R. C., Prakasam, T. B. & Srinath, E. G. Inhibition of nitrification by ammonia and nitrous acid. J. - Water Pollut. Control Fed. 48, 835–852 (1976).

60. Kolmogorov, M. et al. metaFlye: scalable long-read metagenome assembly using repeat graphs. Nat. Methods 17, 1103–1110 (2020).

61. Kalikar, S., Jain, C., Vasimuddin, M. & Misra, S. Accelerating minimap2 for long-read sequencing applications on modern CPUs. Nat. Comput. Sci. 2, 78–83 (2022).

62. Kang, D. D. et al. MetaBAT 2: an adaptive binning algorithm for robust and efficient genome reconstruction from metagenome assemblies. PeerJ 7, e7359 (2019).

63. Pan, S., Zhu, C., Zhao, X.-M. & Coelho, L. P. A deep siamese neural network improves metagenome-assembled genomes in microbiome datasets across different environments. Nat. Commun. 13, 2326 (2022).

64. Lamurias, A., Sereika, M., Albertsen, M., Hose, K. & Nielsen, T. D. Metagenomic binning with assembly graph embeddings. Bioinformatics 38, 4481–4487 (2022).

65. Sieber, C. M. K. et al. Recovery of genomes from metagenomes via a dereplication, aggregation and scoring strategy. Nat. Microbiol. 3, 836–843 (2018).

66. Chklovski, A., Parks, D. H., Woodcroft, B. J. & Tyson, G. W. CheckM2: a rapid, scalable and accurate tool for assessing microbial genome quality using machine learning. Nat. Methods 20, 1203–1212 (2023).

67. Nurk, S., Meleshko, D., Korobeynikov, A. & Pevzner, P. A. metaSPAdes: a new versatile metagenomic assembler. Genome Res. 27, 824–834 (2017).

68. Langmead, B. & Salzberg, S. L. Fast gapped-read alignment with Bowtie 2. Nat. Methods 9, 357–359 (2012).

69. Nawrocki, E. P. & Eddy, S. R. Infernal 1.1: 100-fold faster RNA homology searches. Bioinformatics 29, 2933–2935 (2013).

70. Olm, M. R., Brown, C. T., Brooks, B. & Banfield, J. F. dRep: a tool for fast and accurate genomic comparisons that enables improved genome recovery from metagenomes through de-replication. ISME J. 11, 2864–2868 (2017).

71. Parks, D. H. et al. A complete domain-to-species taxonomy for Bacteria and Archaea. Nat. Biotechnol. 38, 1079–1086 (2020).

72. Chaumeil, P.-A., Mussig, A. J., Hugenholtz, P. & Parks, D. H. GTDB-Tk v2: memory friendly classification with the genome taxonomy database. Bioinformatics 38, 5315–5316 (2022).

73. Shaw, J. & Yu, Y. W. Fast and robust metagenomic sequence comparison through sparse chaining with skani. Nat. Methods 20, 1661–1665 (2023).

74. Eren, A. M. et al. Community-led, integrated, reproducible multi-omics with anvi’o. Nat. Microbiol. 6, 3–6 (2020).

75. Hug, L. A. et al. A new view of the tree of life. Nat. Microbiol. 1, 16048 (2016).

76. Letunic, I. & Bork, P. Interactive Tree of Life (iTOL) v6: recent updates to the phylogenetic tree display and annotation tool. Nucleic Acids Res. 52, W78–W82 (2024).

77. Hyatt, D. et al. Prodigal: prokaryotic gene recognition and translation initiation site identification. BMC Bioinformatics 11, 119 (2010).

78. Shaffer, M. et al. DRAM for distilling microbial metabolism to automate the curation of microbiome function. Nucleic Acids Res. 48, 8883–8900 (2020).

79. Kanehisa, M., Furumichi, M., Tanabe, M., Sato, Y. & Morishima, K. KEGG: new perspectives on genomes, pathways, diseases and drugs. Nucleic Acids Res. 45, D353– D361 (2017).

80. Mistry, J. et al. Pfam: The protein families database in 2021. Nucleic Acids Res. 49, D412– D419 (2021).

81. Sayers, E. W. et al. Database resources of the national center for biotechnology information. Nucleic Acids Res. 50, D20–D26 (2022).

82. Fu, L., Niu, B., Zhu, Z., Wu, S. & Li, W. CD-HIT: accelerated for clustering the next-generation sequencing data. Bioinformatics 28, 3150–3152 (2012).

83. Vallenet, D. et al. MicroScope: an integrated platform for the annotation and exploration of microbial gene functions through genomic, pangenomic and metabolic comparative analysis. Nucleic Acids Res. gkz926 (2019) doi:10.1093/nar/gkz926.

84. Karp, P. D. et al. Pathway Tools version 23.0 update: software for pathway/genome informatics and systems biology. Brief. Bioinform. 22, 109–126 (2021).

85. Ong, W. K., Midford, P. E. & Karp, P. D. Taxonomic weighting improves the accuracy of a gap-filling algorithm for metabolic models. Bioinformatics 36, 1823–1830 (2020).

86. Plugge, C. M. Anoxic Media Design, Preparation, and Considerations. in Methods in Enzymology vol. 397 3–16 (Elsevier, 2005).

87. Valero, D., Montes, J. A., Rico, J. L. & Rico, C. Influence of headspace pressure on methane production in Biochemical Methane Potential (BMP) tests. Waste Manag. 48, 193– 198 (2016).

88. Rinke, C. et al. Obtaining genomes from uncultivated environmental microorganisms using FACS–based single-cell genomics. Nat. Protoc. 9, 1038–1048 (2014).

89. Brent, R. P. *Algorithms for Minimization without Derivatives*. (Dover Publications, Mineola, N.Y, 2002).

90. Hatzenpichler, R. & Orphan, V. J. Detection of Protein-Synthesizing Microorganisms in the Environment via Bioorthogonal Noncanonical Amino Acid Tagging (BONCAT). in Hydrocarbon and Lipid Microbiology Protocols (eds. McGenity, T. J., Timmis, K. N. & Nogales, B.) 145–157 (Springer Berlin Heidelberg, Berlin, Heidelberg, 2015). doi:10.1007/8623_2015_61.

91. Love, M. I., Huber, W. & Anders, S. Moderated estimation of fold change and dispersion for RNA-seq data with DESeq2. Genome Biol. 15, 550 (2014).

92. Xu, K. et al. Benchtop-compatible sample processing workflow for proteome profiling of < 100 mammalian cells. Anal. Bioanal. Chem. 411, 4587–4596 (2019).

93. Nicora, C. D. et al. The MPLEx Protocol for Multi-omic Analyses of Soil Samples. J. Vis. Exp. 57343 (2018) doi:10.3791/57343.

94. Kim, Y.-M. et al. Diel metabolomics analysis of a hot spring chlorophototrophic microbial mat leads to new hypotheses of community member metabolisms. Front. Microbiol. 6, (2015).

95. Hiller, K. et al. MetaboliteDetector: Comprehensive Analysis Tool for Targeted and Nontargeted GC/MS Based Metabolome Analysis. Anal. Chem. 81, 3429–3439 (2009).

96. Dai, C. et al. quantms: a cloud-based pipeline for quantitative proteomics enables the reanalysis of public proteomics data. Nat. Methods (2024) doi:10.1038/s41592-024-02343-1.

97. Ewels, P. A. et al. The nf-core framework for community-curated bioinformatics pipelines. Nat. Biotechnol. 38, 276–278 (2020).

98. The Bioconda Team et al. Bioconda: sustainable and comprehensive software distribution for the life sciences. Nat. Methods 15, 475–476 (2018).

99. Da Veiga Leprevost, F., et al. BioContainers: an open-source and community-driven framework for software standardization. Bioinformatics 33, 2580–2582 (2017).

100. Pfeuffer, J. et al. OpenMS 3 enables reproducible analysis of large-scale mass spectrometry data. Nat. Methods 21, 365–367 (2024).

101. Di Tommaso, P. et al. Nextflow enables reproducible computational workflows. Nat. Biotechnol. 35, 316–319 (2017).

102. Kim, S. & Pevzner, P. A. MS-GF+ makes progress towards a universal database search tool for proteomics. Nat. Commun. 5, 5277 (2014).

103. Eng, J. K., Jahan, T. A. & Hoopmann, M. R. Comet: An open-source MS/MS sequence database search tool. PROTEOMICS 13, 22–24 (2013).

104. Elias, J. E. & Gygi, S. P. Target-decoy search strategy for increased confidence in large-scale protein identifications by mass spectrometry. Nat. Methods 4, 207–214 (2007).

105. Käll, L., Canterbury, J. D., Weston, J., Noble, W. S. & MacCoss, M. J. Semi-supervised learning for peptide identification from shotgun proteomics datasets. Nat. Methods 4, 923– 925 (2007).

106. R Core Team. R: A language and environment for statistical computing. R Foundation for Statistical Computing https://www.R-project.org/ (2022).

107. Wickham, H. et al. Welcome to the Tidyverse. J. Open Source Softw. 4, 1686 (2019).

108. Zhang, X. et al. Metaproteomics reveals associations between microbiome and intestinal extracellular vesicle proteins in pediatric inflammatory bowel disease. Nat. Commun. 9, 2873 (2018).

109. Li, L. et al. Revealing proteome-level functional redundancy in the human gut microbiome using ultra-deep metaproteomics. Nat. Commun. 14, 3428 (2023).

110. Sachsenberg, T. et al. MetaProSIP: Automated Inference of Stable Isotope Incorporation Rates in Proteins for Functional Metaproteomics. J. Proteome Res. 14, 619–627 (2015).

111. Berthold, M. R. et al. KNIME - the Konstanz information miner: version 2.0 and beyond. ACM SIGKDD Explor. Newsl. 11, 26–31 (2009).

112. Stahl, D. A. Development and application of nucleic acid probes in bacterial systematics. Seq. Hybrid. Tech. Bact. Syst. (1991).

113. Meier, H., Amann, R., Ludwig, W. & Schleifer, K. H. Specific Oligonucleotide Probes for in situ Detection of a Major Group of Gram-positive Bacteria with low DNA G+C Content. Syst. Appl. Microbiol. 22, 186–196 (1999).

114. Ebrahim, A., Lerman, J. A., Palsson, B. O. & Hyduke, D. R. COBRApy: COnstraints-Based Reconstruction and Analysis for Python. BMC Syst. Biol. 7, 74 (2013).

115. Holzhütter, H.-G. The principle of flux minimization and its application to estimate stationary fluxes in metabolic networks. Eur. J. Biochem. 271, 2905–2922 (2004).

116. Edgar, R. C. Search and clustering orders of magnitude faster than BLAST. Bioinformatics 26, 2460–2461 (2010).

