## Supplemental Figures for "Activity-targeted metaproteomics uncovers rare syntrophic bacteria central to anaerobic community metabolism"

### Activity-targeted metaproteomics enhances the ecophysiological characterization of cryptic syntrophic metabolisms

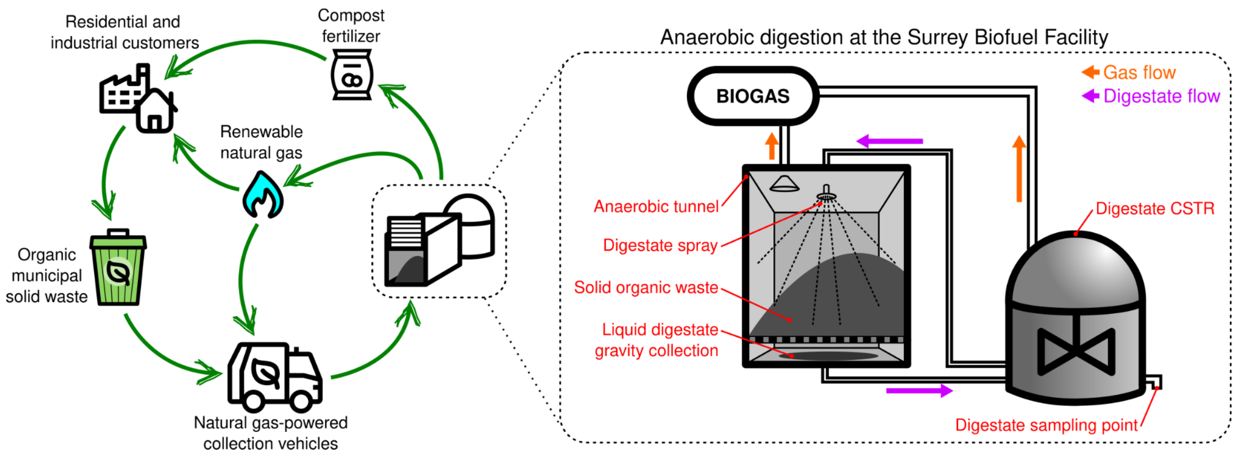


**Supplemental Figure S1.** Process diagram of the Surrey Biofuel Facility. (CSTR = continuous stirred tank reactor)

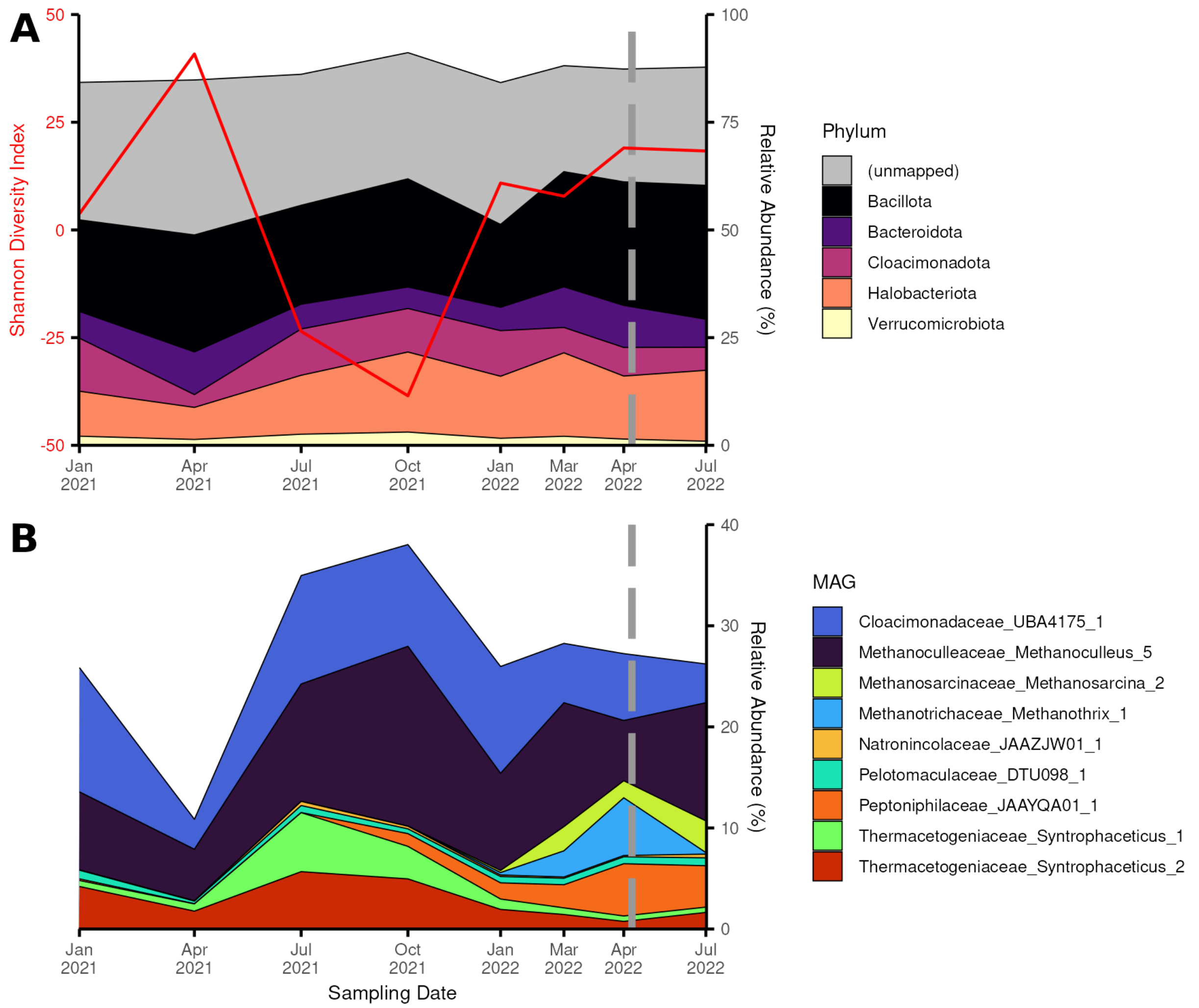


**Supplemental Figure S2.** Taxonomic composition of the time series deep sequencing reads. Dashed line indicates the date of digestate collection for the microcosm incubation. (a) The left axis and red line indicate the Shannon Diversity Index at each sampling event. The right axis and colored areas represent the relative abundance of the five most abundant phyla across the time series. The proportion of unmapped reads are also shown and lower abundance phyla are excluded. (b) Sequencing read coverage of a subset of MAGs in the time series samples. Both highly abundant MAGs and MAGs of interest due to the metaproteomic results are shown. A notable exclusion is *Pelotomaculaceae_Pelotomaculum_C_1* which had an abundance so low in every sample that it could not be displayed.

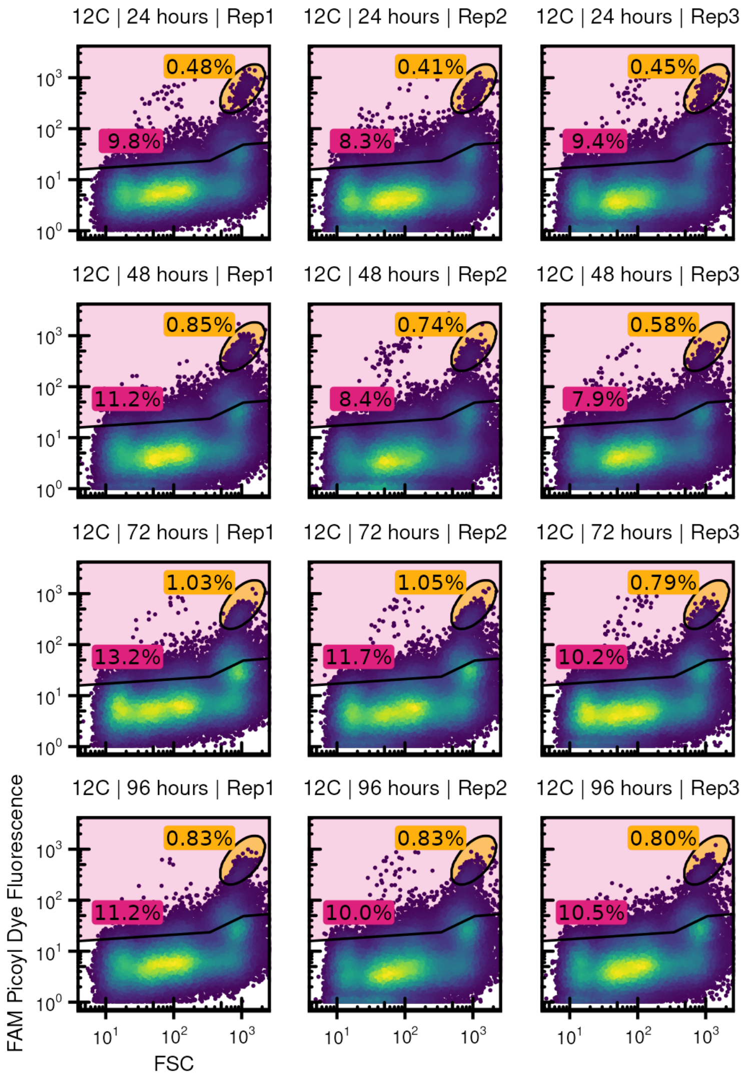


**Supplemental Figure S3.** Gating results for all BONCAT Sorted Cells mini-metagenome samples. Percentages in pink and yellow represent the fractions of SYTO+ events passing the BONCAT+ and Bright++ gates respectively.

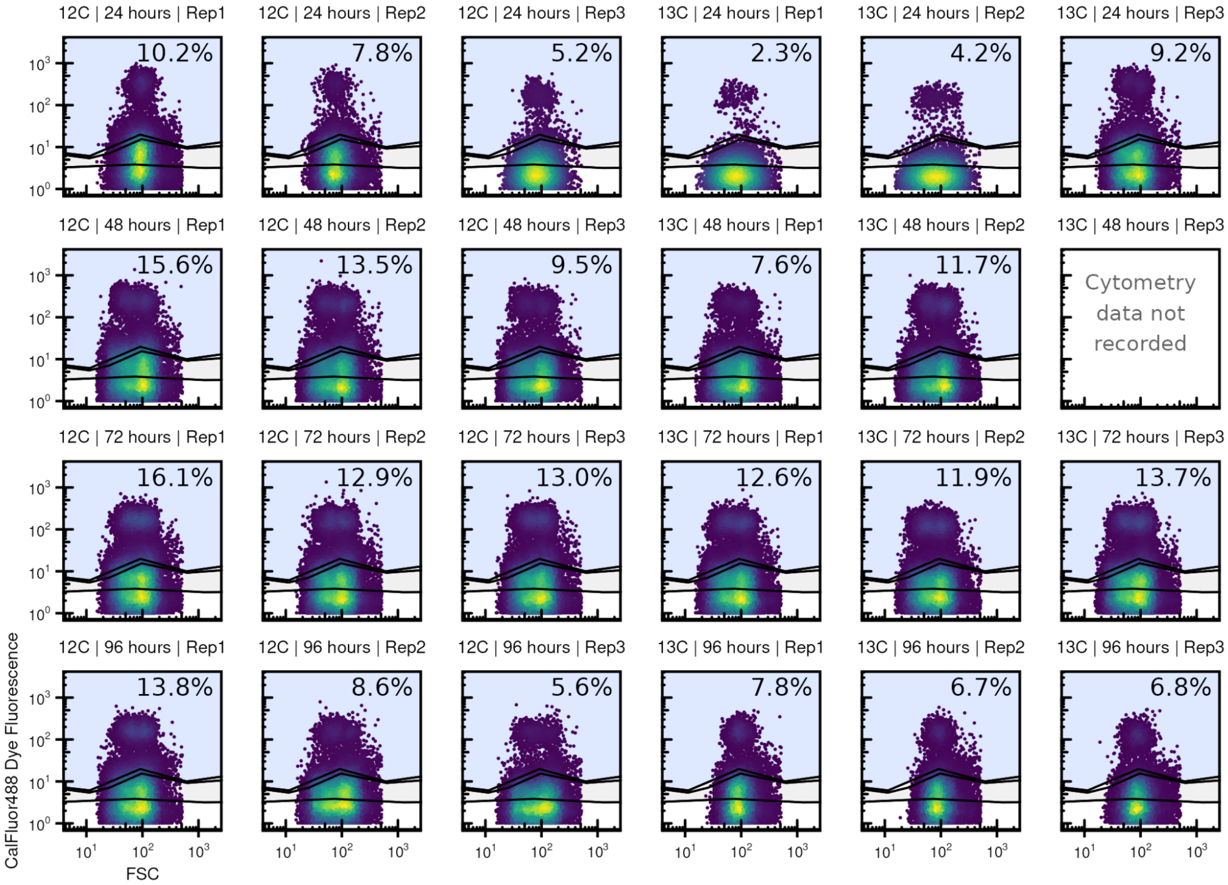


**Supplemental Figure S4.** Gating results for all BONCAT Sorted Cells metaproteomics samples. Percentages represent the fraction of SYTO+ events passing the BONCAT+ gate.

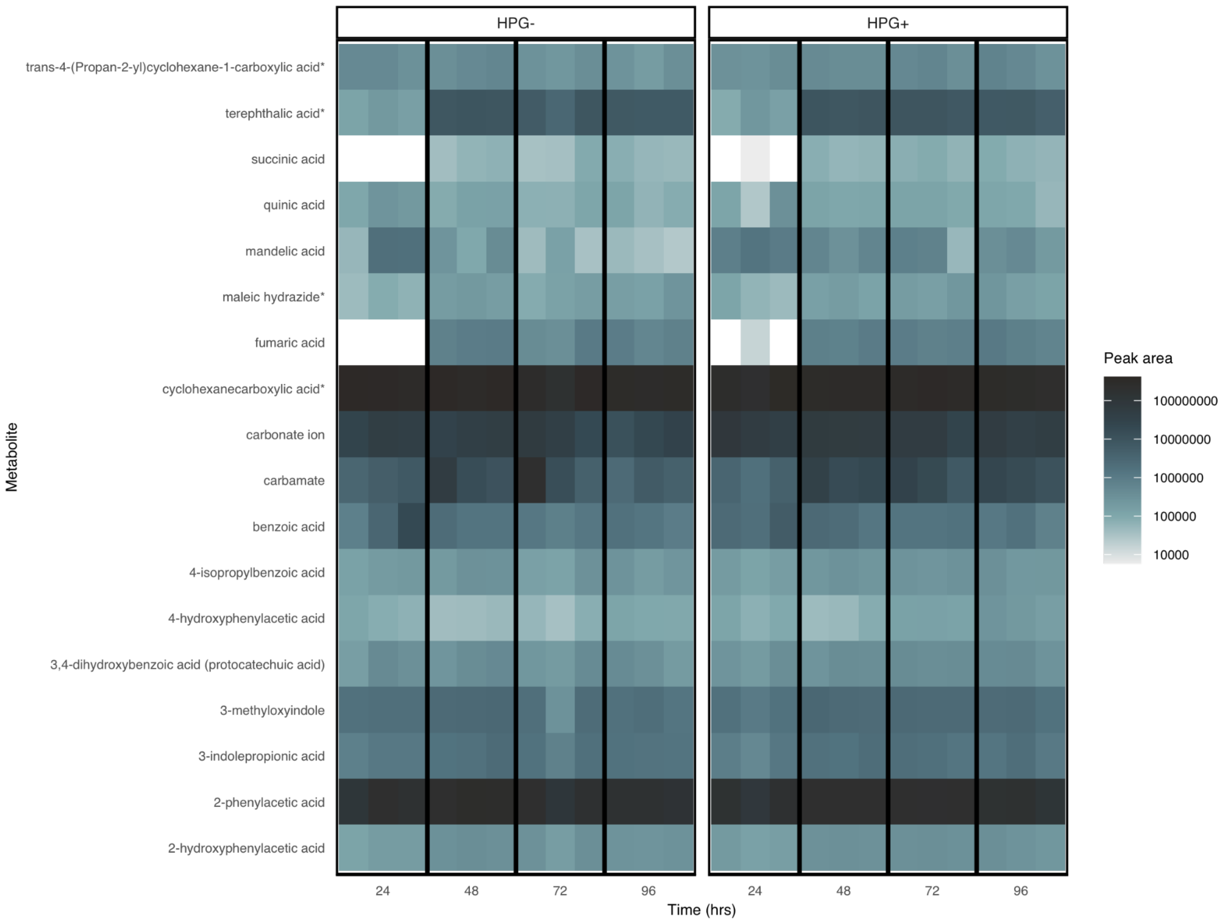


**Supplemental Figure S5:** Relative levels of metabolites measured by GC-MS in the SIP microcosms with $^{12}C$-acetate. Metabolites with a * next to the name were identified based on fragmentation patterns only (NIST library); all others were identified based on fragmentation and retention indices from PNNL augmented version of Agilent GC-MS metabolomics Library.

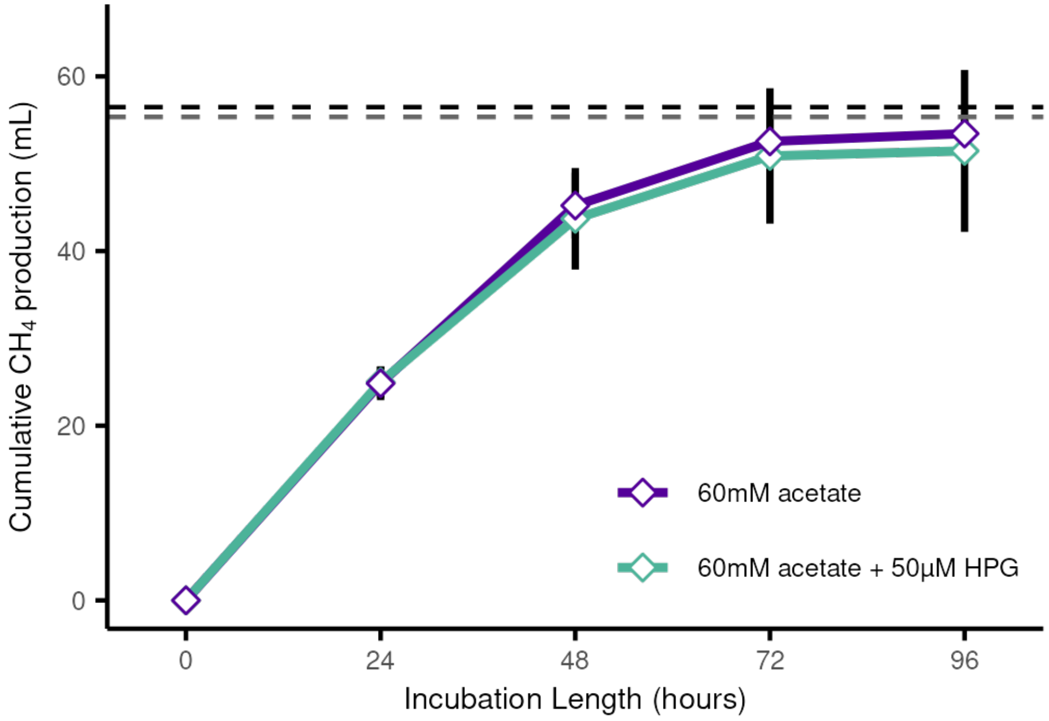


**Supplemental Figure S6.** Methane production over time in the SIP incubations with and without HPG. The black dashed line represents the maximum theoretical methane yield based on 1 mol acetate / 1 mol methane conversion of 2.52 mmol sodium acetate amendment (vertical bars represent standard deviation, n varies from 6 - 24). The dashed grey line represents the maximum expected methane yield after accounting for biomass yield (0.02 $gCOD_{biomass}/gCOD_{removed}$ for SAO consortia[1]).

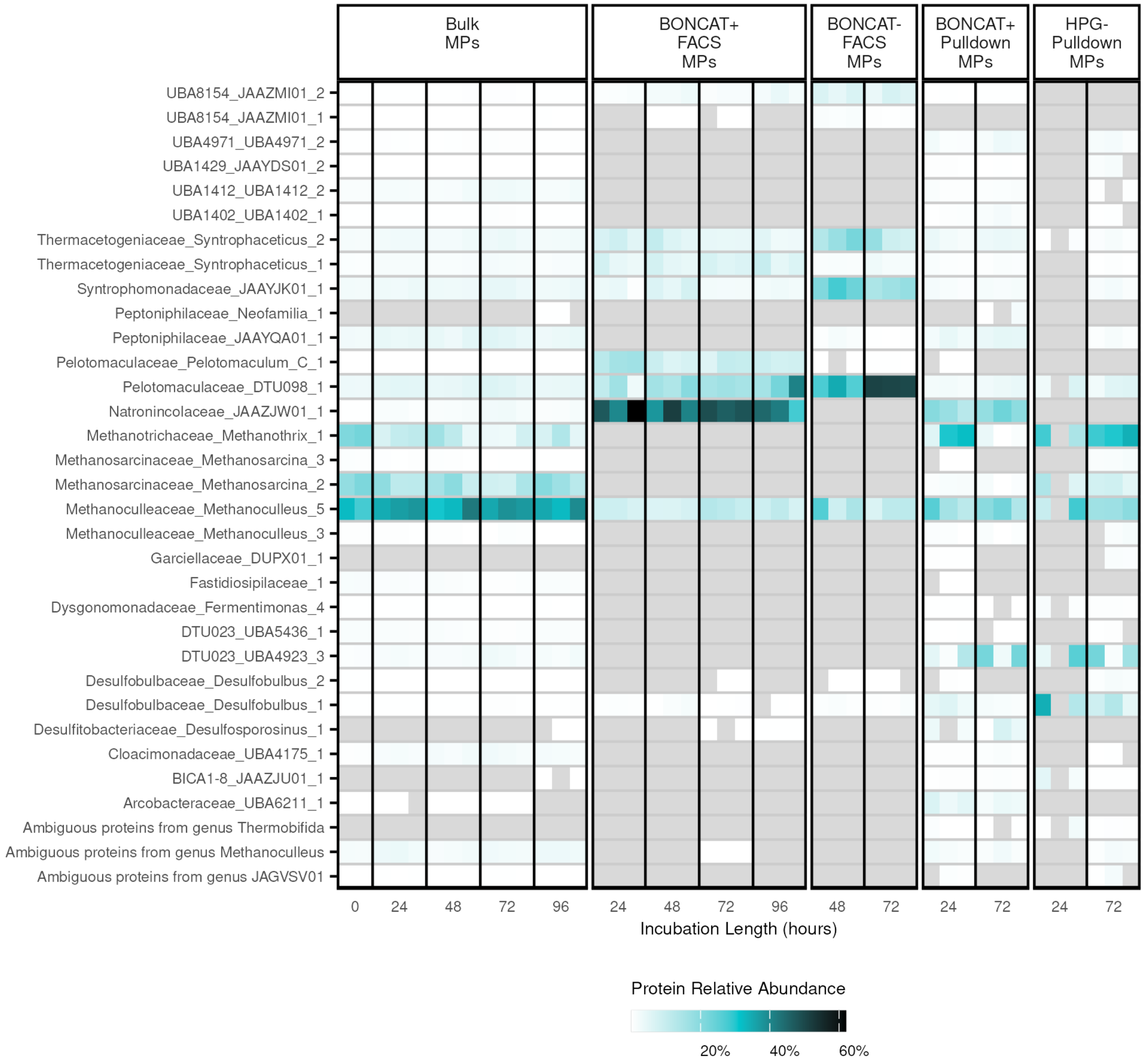


**Supplemental Figure S7.** Relative protein abundance of detected MAGs (> 1% relative abundance in at least 1 sample) across all samples. Missing values are shown in grey.

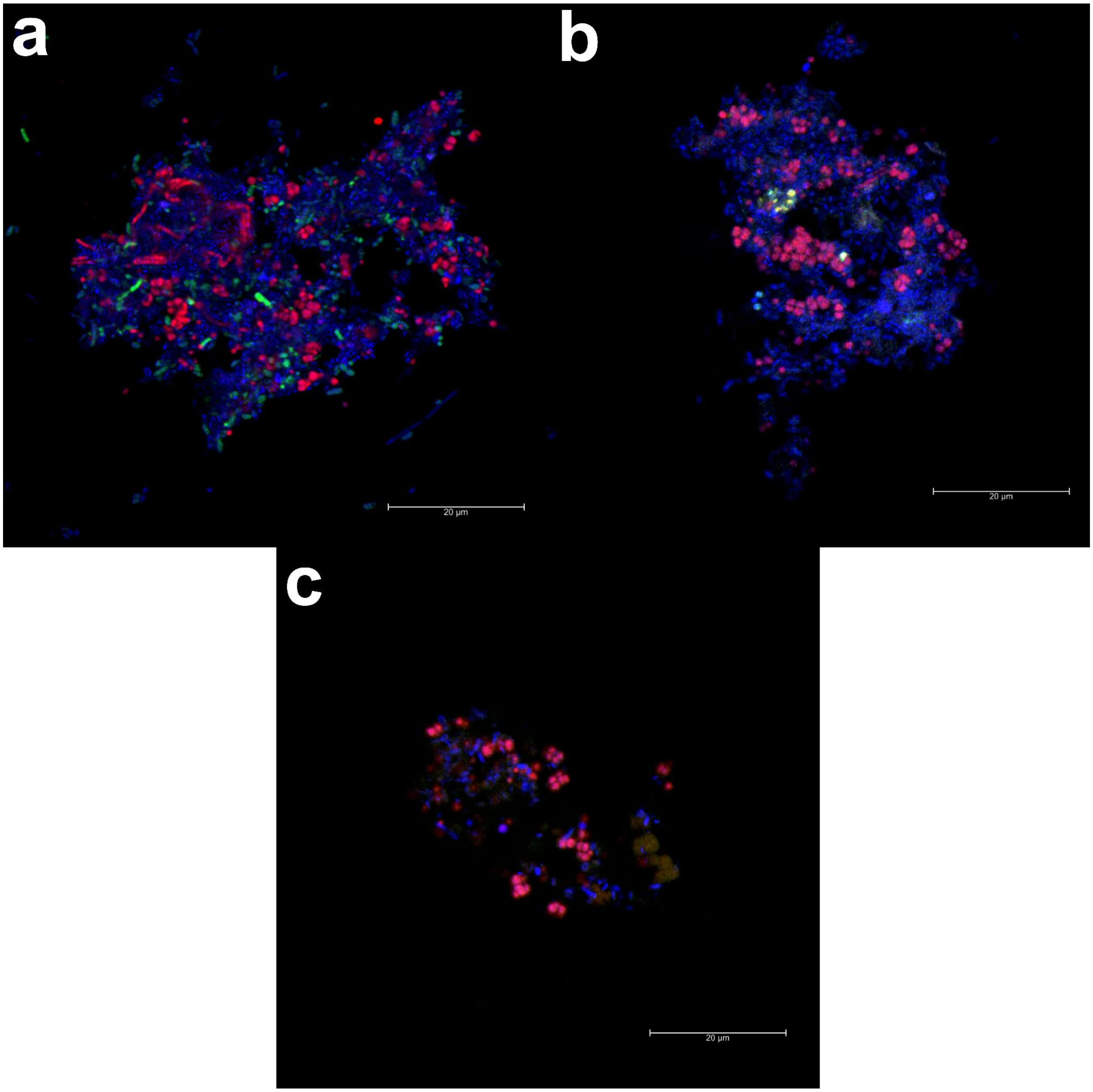


**Supplemental Figure S8.** Fluorescence in situ hybridization (FISH) images of the microbial biomass in the $^{12}C$-acetate microcosm incubations collected at 96 hours. The probes and stains used were: Arch915F targeting *Archaea* (Cy3; red); LGC mix targeting *Bacillota* (i.e., *Firmicutes*) (Fluos; green), and DAPI (blue). The images were obtained using confocal microscopy (see Online Methods).

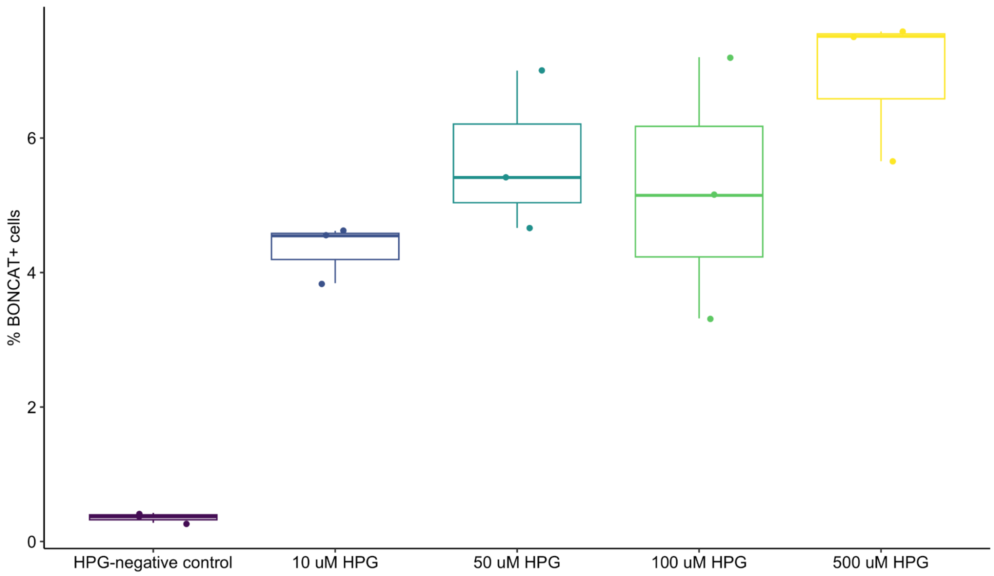


**Supplemental Figure S9:** The impact of HPG concentration on the fraction of SYTO-stained cells that were BONCAT-positive in biomass from the Surrey Biofuel Facility incubated in anoxic conditions with 50 mM of acetate. Cell staining and click chemistry was performed in the same was as the BONCAT-FACS MPs described in Supplemental Table S12, and cell counts were determined using flow cytometry on a BD FACSJazz cell sorter (BD Biosciences, USA) with the gating strategy outlined in the Online Methods.

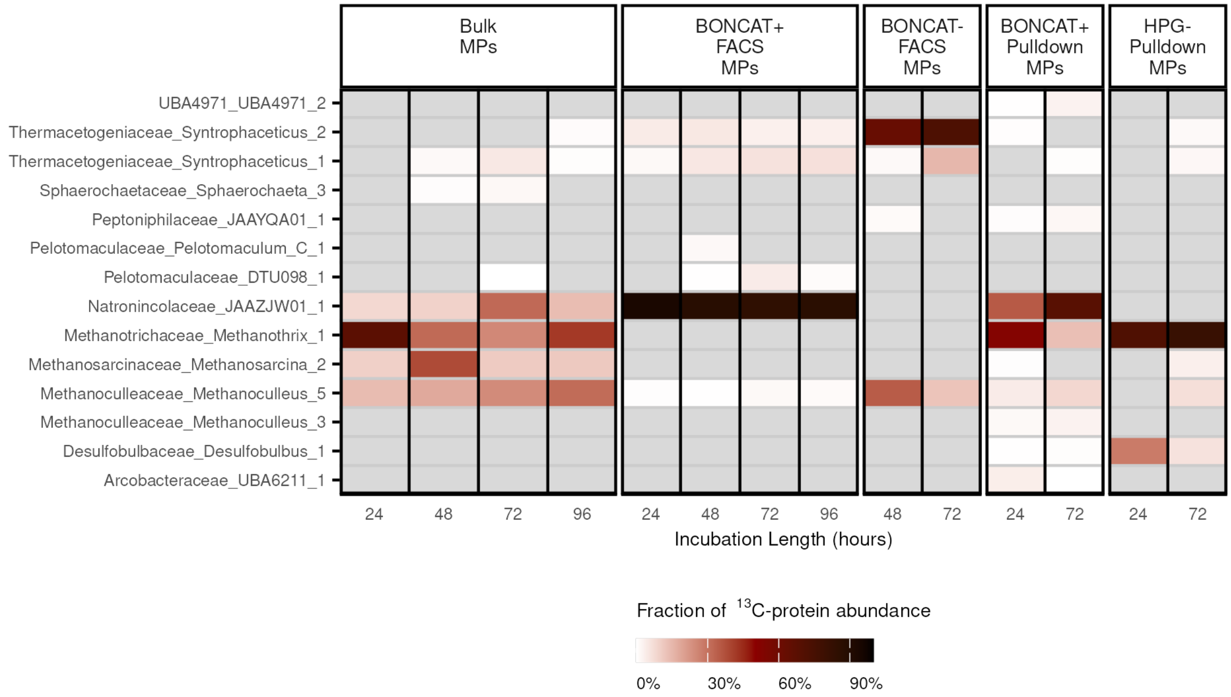


**Supplemental Figure S10.** Fraction of $^{13}$C-protein abundance from detected MAGs (> 1% of $^{13}$C-protein abundance in at least one sample) across all samples. Missing values are shown in grey.

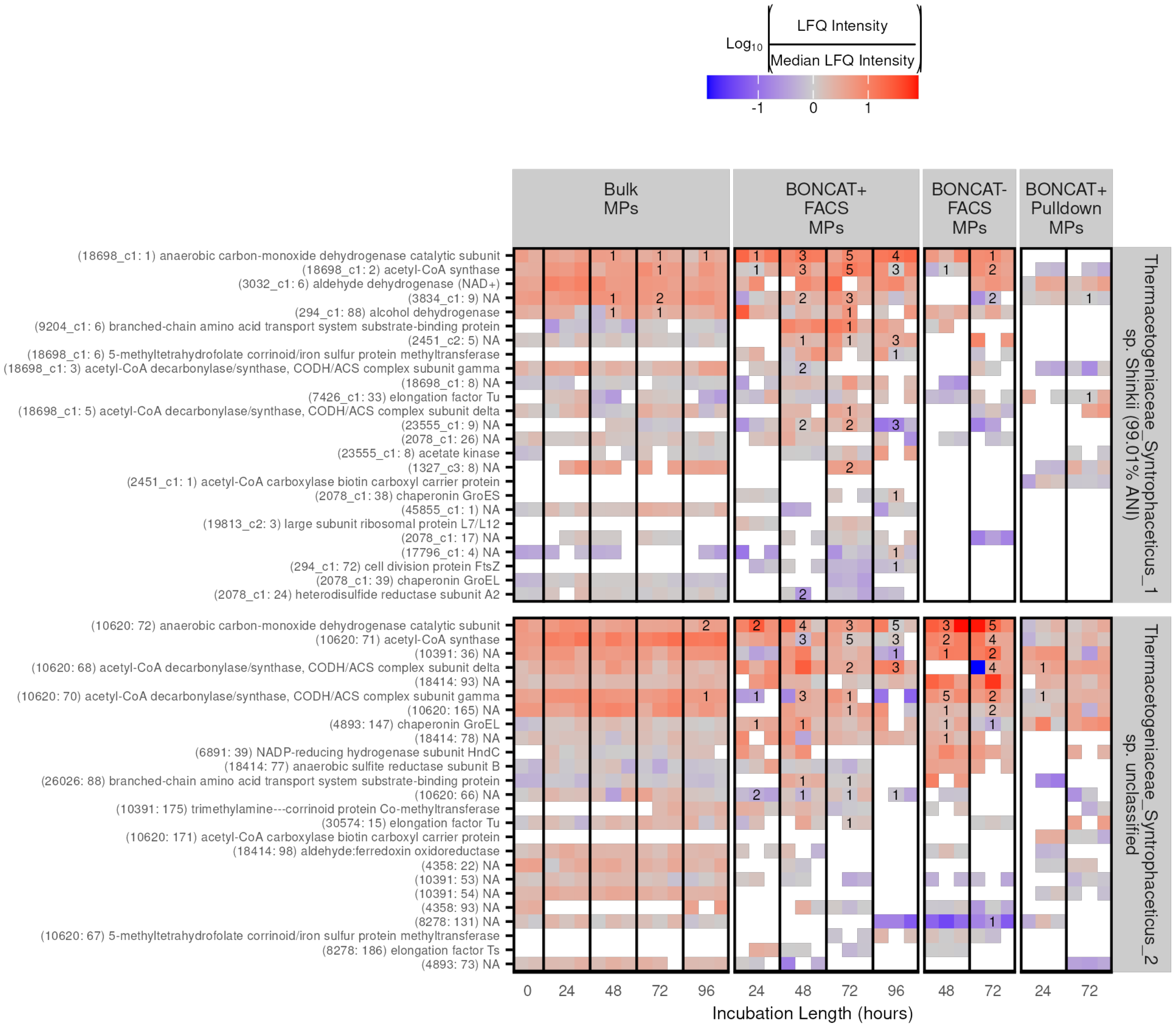


**Supplemental Figure S11.** Top 25 highest expression proteins in MAGs from genus *Syntrophaceticus.* Protein expression was calculated as the mean MS intensity of the top 3 peptides detected from that protein. Protein expression was scaled by dividing by the median protein expression from the MAG in that sample and log-transformation. Numbers represent $^{13}$C-labeled peptides detected across $^{13}$C-incubation samples (n = 3) at the given time point. Labels are “(contig: orf) KEGG annotation”.

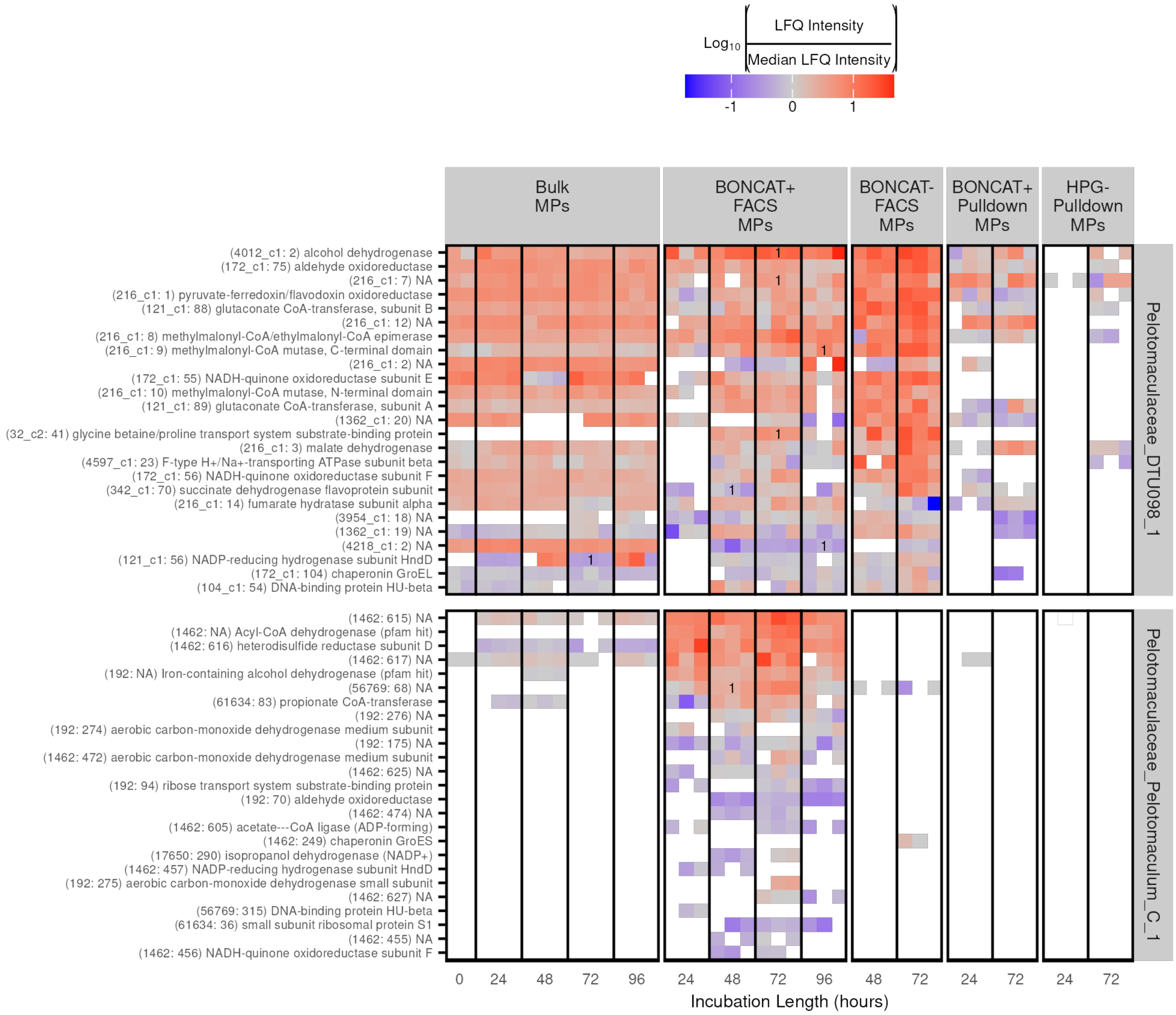


**Supplemental Figure S12.** Top 25 highest expression proteins from two potential SPOB that were enriched by the BONCAT methods but did not incorporate substantial $^{13}C$-labeled acetate. Protein expression was calculated as the mean MS intensity of the top 3 peptides detected from that protein. Protein expression was scaled by dividing by the median protein expression from the MAG in that sample and log-transformation. Numbers represent $^{13}C$-labeled peptides detected across $^{13}C$-incubation samples (n = 3) at the given time point. Labels are “(contig: orf) KEGG annotation”.

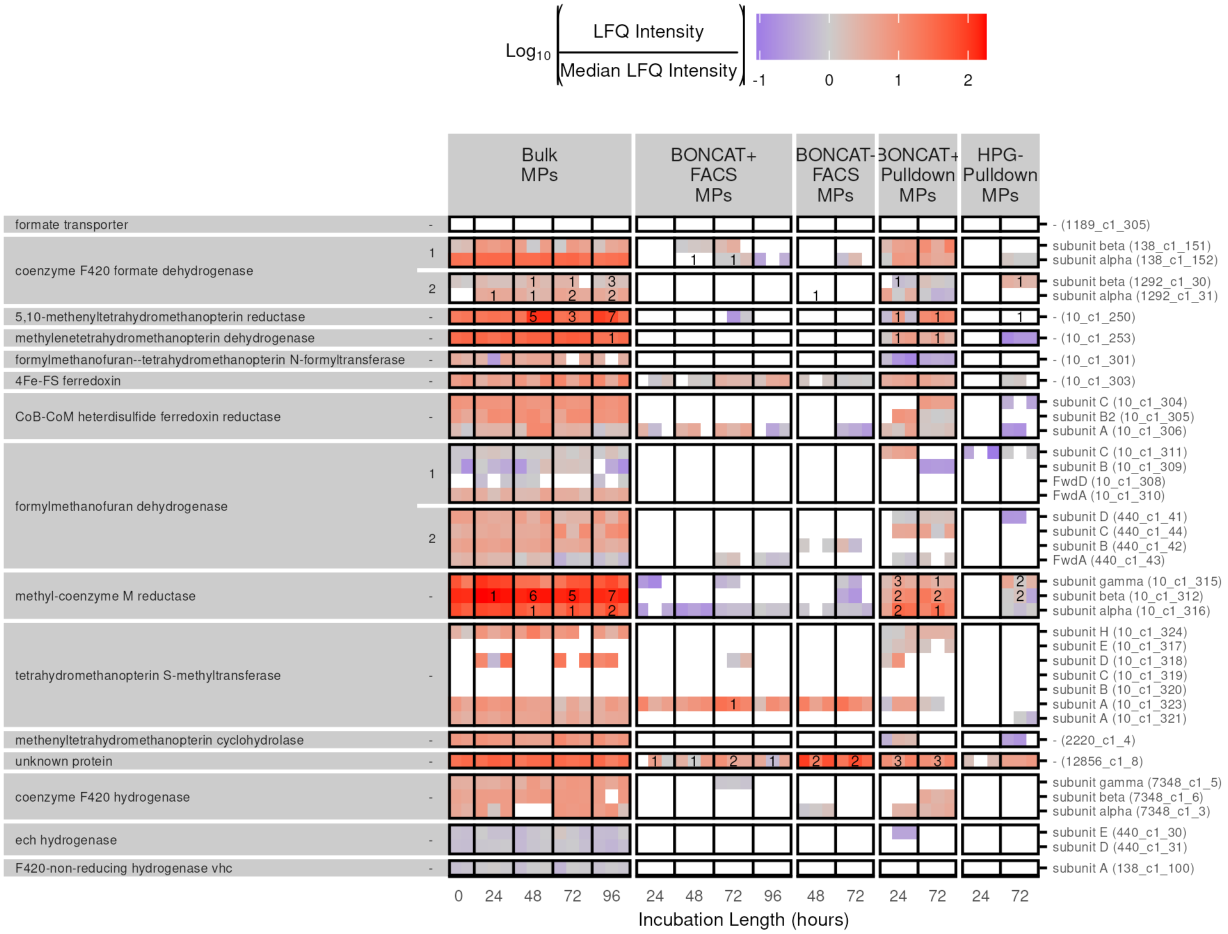


**Supplemental Figure S13.** Genes related to $CO_2$ metabolism in the hydrogenotrophic methanogen *Methanoculleaceae_Methanoculleus_5*. This MAG encodes two separate formate dehydrogenase clusters that were highly expressed indicating the likelihood that it accepts formate from *Natronincolaceae_JAAZJW01_1* as an electron carrier. Numbers on the left between the enzyme name and the heatmap indicate distinct gene clusters. Protein expression was calculated as the mean MS intensity of the top 3 peptides detected from that protein. Protein expression was scaled by dividing by the median protein expression from the MAG in that sample and log-transformation. Numbers represent unique $^{13}C$-labeled peptides detected across $^{13}C$-incubation samples (n = 3) at the given time point. Labels from left to right are: enzyme complex (MicroScope annotation), distinct genome cluster (or “-” if only one), subunit (or “-” if none), and finally “contig: orf”.

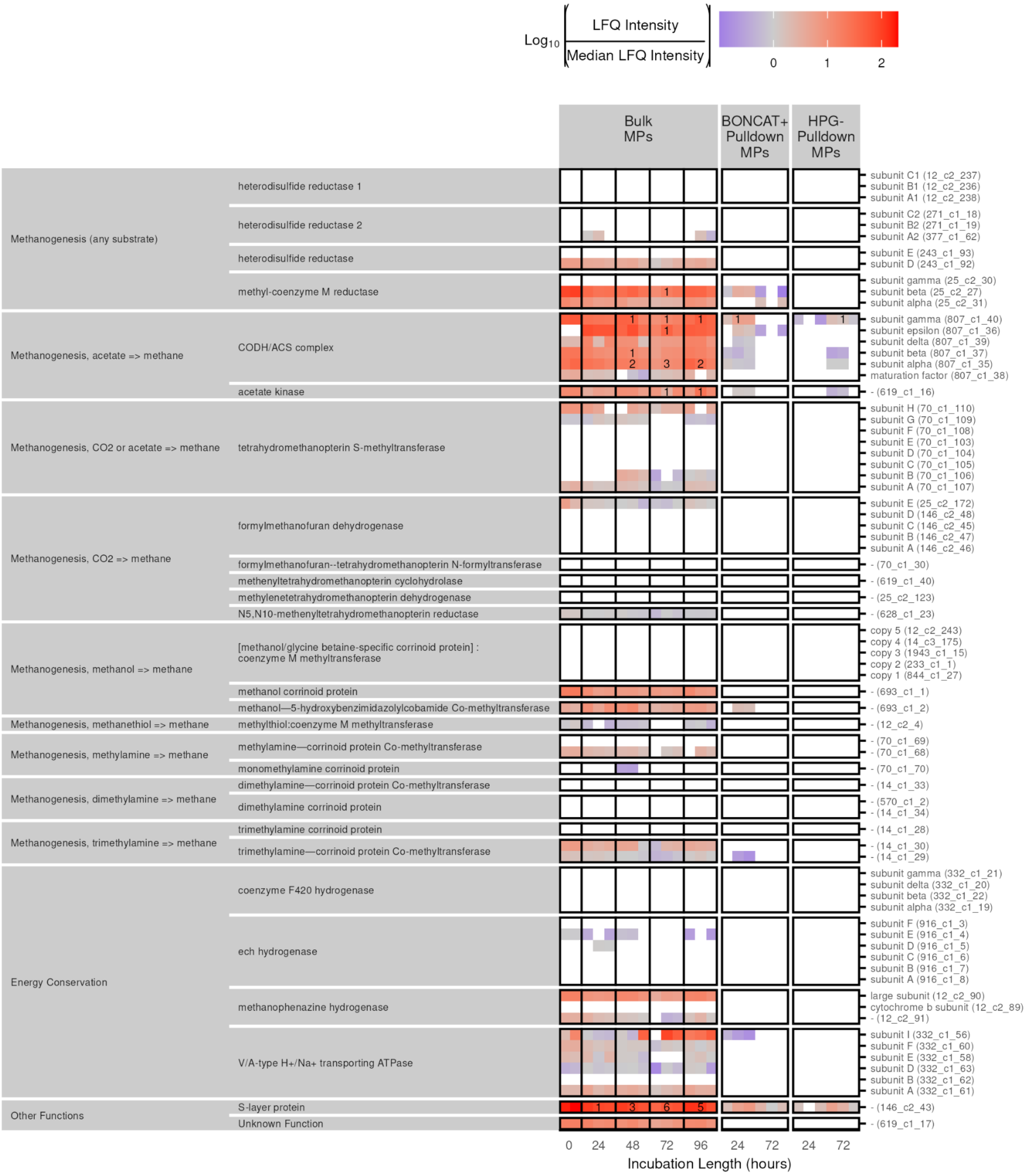


**Supplemental Figure S14.** Genes related to methanogenic metabolisms in *Methanosarcinaceae_Methanosarcina_2*. This MAG showed the highest expression of the acetoclastic pathway and many enzymes required for $CO_2$ or formate utilization were not detected proteomically. Protein expression was calculated as the mean MS intensity of the top 3 peptides detected from that protein. Protein expression was scaled by dividing by the median protein expression from the MAG in that sample and log-transformation. Numbers represent unique $^{13}C$-labeled peptides detected across $^{13}C$-incubation samples (n = 3) at the given time point. Labels from left to right indicate: metabolic pathway (KEGG), enzyme complex (MicroScope annotation), subunit or identical copy (or "-" if none), and finally "contig: orf".

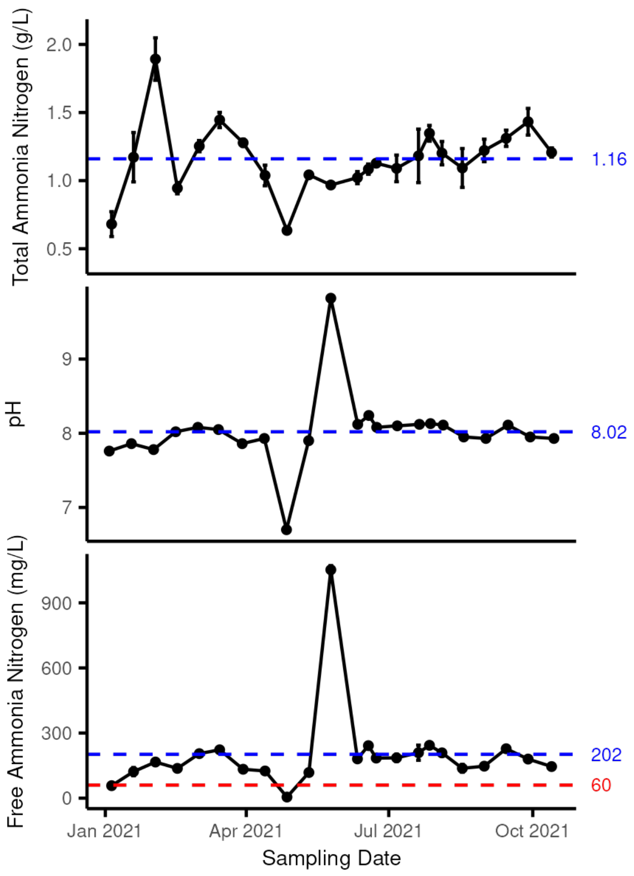


**Supplemental Figure S15.** Measured total ammonia nitrogen, measured pH and calculated free ammonia nitrogen (Anthonisen et. al. 1976) in liquid digestate from the Surrey Biofuel Facility across 10 months of sampling (vertical bars represent standard deviation, n varies from 3 - 6). The temperature of the reactor is maintained at 37°C. Blue lines indicate mean across shown sampling events. The red line indicates free ammonia nitrogen threshold at which acetoclastic methanogenesis is reduced to 75% activity (Jiang et al., 2015)[2].

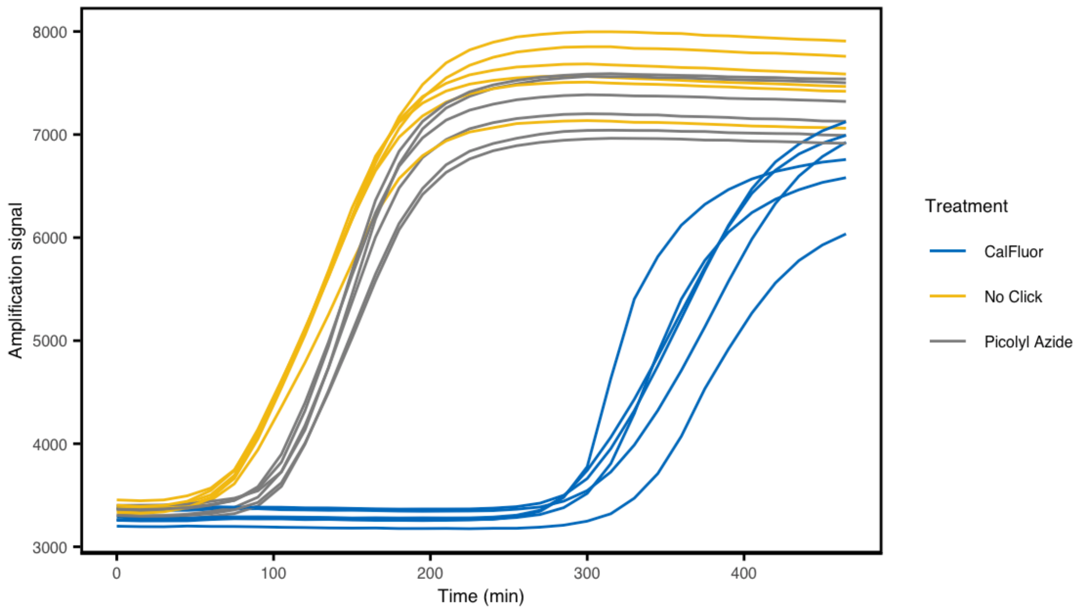


**Supplemental Figure S16:** Multiple displacement amplification results on cells of *Escherichia coli* grown in the presence of HPG, followed by click-chemistry with either CalFluor or picolyl azide dye (with no click-chemistry as a negative control). MDA of the *E. coli* DNA was performed in the presence of SYTO9 stain in a real-time PCR machine with the filter set to SYBR Green fluorescence to measure the amplification signal. The inhibition of MDA caused by CalFluor dye is apparent by the delayed amplification curve relative to picolyl azide dye and the no-click control.

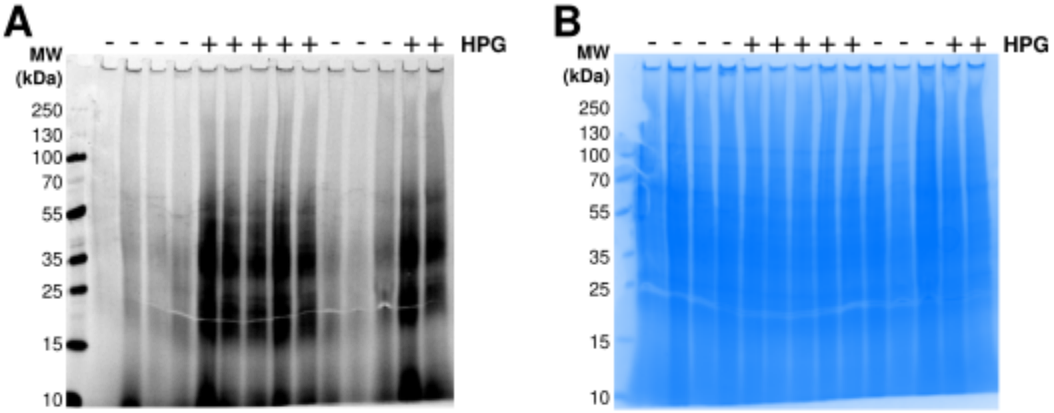


**Supplemental Figure S17.** Images of SDS-PAGE gels containing extracted protein following click-chemistry with rhodamine-azide. (A) Representative fluorescence gel and (B) corresponding gel after total protein stain with GelCode blue.

### References


1. McDaniel, E. A. *et al.* Diverse electron carriers drive syntrophic interactions in an enriched anaerobic acetate-oxidizing consortium. *ISME J.* **17**, 2326–2339 (2023).
2. Jiang, Y. *et al.* Ammonia inhibition and toxicity in anaerobic digestion: A critical review. *J. Water Process Eng.* **32**, 100899 (2019).
